# The host range and the role of O-antigen in P1 transduction with its alternative S’ tail fibre

**DOI:** 10.1101/2022.07.24.501295

**Authors:** Yang W. Huan, Jidapha Fa-arun, Baojun Wang

## Abstract

Enterobacteria phage P1 expresses two types of tail fibre, S and S’. Despite the wide usage of phage P1 for transduction, the host range and the receptor for its alternative S’ tail fibre was never determined. Here, a Δ*S-cin* Δ*pac E. coli* P1 lysogenic strain was generated to allow packaging of phagemid DNA into P1 phage having either S or S’ tail fibre. P1(S’) could transduce phagemid DNA into *Shigella flexneri* 2a 2457O, *Shigella flexneri* 5a M90T and *Escherichia coli* O3 efficiently. Mutational analysis of the O-antigen assembly genes and LPS inhibition assays indicated that P1(S’) transduction requires at least one O-antigen unit. *E. coli* O111:B4 LPS produced a high neutralising effect against P1(S’) transduction, indicating that this *E. coli* strain could be a host for P1(S’). Mutations in the O-antigen modification genes of *S. flexneri* 2a 2457O and *S. flexneri* 5a M90T did not cause significant changes to P1(S’) transduction efficiency. A higher transduction efficiency of P1(S’) improved the delivery of a *cas9* antimicrobial phagemid into both *S. flexneri* 2457O and M90T. These findings provide novel insights into P1 tropism-switching, by identifying the host range of P1(S’) and demonstrating its potential for delivering a sequence-specific Cas9 antimicrobial into clinically relevant *S. flexneri*.

## INTRODUCTION

Enterobacteria phage P1 has a wide host range of gram-negative *Enterobacteriaceae*, including multiple strains of *Escherichia coli* (Bertani, 1951; Bertani, 1958; Goldhar et al., 1973; Goldberg et al., 1974; Murooka and Harada, 1979; Westwater et al., 2002), *Shigella flexneri* (Westwater et al., 2002), *S. dysenteriae* (Bertani, 1951; Bertani, 1958; Goldhar et al., 1973; Westwater et al., 2002), *Klebsiella pneumoniae* (Goldberg et al., 1974; Murooka and Harada, 1979; Westwater et al., 2002), *Citrobacter freundii* (Goldberg et al., 1974; Murooka and Harada, 1979; Westwater et al., 2002) and *Pseudomonas aeruginosa* (Goldberg et al., 1974; Murooka and Harada, 1979; Westwater et al., 2002). Genes that determine P1 phage adsorption and host specificity are associated with the *cin* operon, as well as the tail fibre operon encoding for three genes, *R*, *S*, *U* under the regulation of late promoter LP_*Sit*_ (Guidolin et al., 1989a; Guidolin et al., 1989b; Łobocka et al., 2004). S is a structural protein that extends along the tail fibre with a constant N-terminus (Sc) attached to the baseplate while its variable C-terminus (Sv) binds to a specific receptor on the host bacterium allowing for phage adsorption (Yarmolinsky and Sternberg, 1988). *U* was predicted to encode an essential chaperone protein for the assembly of a distal part of the tail fibre (Łobocka et al., 2004). The coding sequences of *Sv* and *U* are arranged in an opposite orientation to the alternative variants, namely *Sv’* and *U*’, which together form the invertible C segment flanked by inverted repeats, *cixL* and *cixR* (Iida et al., 1982; Hiestand-Nauer and Iida, 1983; Iida, 1984). Inversion of the C segment at the *cix* sites is mediated by the Cin recombinase, which allows for expression of the alternative S’ tail fibre and its assembly protein U’, changing the host range specificity of P1 (Iida et al., 1982). Inversion of the C segment, however, is thought to be infrequent hence most of the P1 phage progeny generated lytically (i.e., phage progeny from a single plaque or from continuous lysis of cell culture) would have the same tail fibre as the infecting phage whereas the induction of lysogenic host cell would generate a mixture of P1 phage with either the S or S’ tail fibre (Iida et al., 1982; Hiestand-Nauer and Iida, 1983; Iida, 1984).

The host range of P1 with its S tail (termed P1(S) hereafter) include *E. coli* K12 (Iida et al., 1982; Hiestand-Nauer and Iida, 1983; Iida, 1984), *E. coli* C (Iida et al., 1982; Hiestand-Nauer and Iida, 1983; Iida, 1984) and *S. dysenteriae* strain “Sh” (Iida, 1984). Previous studies of *E. coli* and *Salmonella enterica* serovar Typhimurium lipopolysaccharide (LPS)-defective mutants suggested that glucose molecule(s) on the outer core of LPS might serve as the receptor for P1 S tail fibre (Franklin, 1969; Ornellas and Stocker, 1974; Sandulache et al., 1984). P1 with its S’ tail fibre (termed P1(S’) in this study), however, failed to transduce into either *E. coli* K12 or *E. coli* C and its bacterial host range remained unknown (Iida et al., 1982; Iida, 1984). On the contrary, a mutant strain of *E. coli*strain C that supports transduction and plaque formation with both P1(S) and P1(S’) was identified in a previous study, although the nature of the mutation(s) was not known (Iida, 1984).

Analysis of P1 *S*, *U* and the alternative *S*’, *U* genes sequences suggested extensive homologies of these proteins to the genes’ products of Bacteriophage Mu invertible G segment (Łobocka et al., 2004). Consistently, the P1 *S* and *U* genes were shown to complement and rescue transduction of Mu *S* and *U* mutants at varying efficiencies (Toussaint et al., 1978). The genes of Mu G segment, however, failed to complement amber mutations within the P1 C segment, suggesting functional differences between Mu and P1 tail fibres and chaperones despite high DNA sequence homology. This was consistent with discrepancies in the host range of Mu(S) and P1(S), as Mu(S) can only infect *E. coli* K12 but not *E. coli* C while P1(S) can infect both strains of *E. coli* (Sandulache et al., 1984).

In this study, we have created a mutant strain of an *E. coli* P1 lysogen in which the entire C segment and the *cin* recombinase coding sequence has been deleted. The mutations were complemented *in trans* by providing the *S* and *U* or the *S*’ and *U’* genes from a plasmid, giving phage progeny with homogenous S or S’ tail fibres. Our data demonstrated that P1(S’) phage could transduce phagemid DNA into *S. flexneri* 2a 2457O, *S. flexneri* 5a M90T and *E. coli* O3 at a high efficiency. Transduction of phagemid DNA by P1(S’) was inhibited on Δ*waaL* and Δ*rfe* mutant strains of these bacteria lacking O-antigen. Furthermore, mutant strains of *S. flexneri* 2a 2457O, *S. flexneri* 5a M90T and *E. coli* O3 with the O-antigen polymerase gene *wzy/rfc* deleted were susceptible to P1(S’) transduction, indicating that P1(S’) transduction requires at least one O-antigen unit. Consistently, LPS extracted from wildtype *S. flexneri* 2a 2457O, *S. flexneri* 5a M90T, *E. coli* O3 and *E. coli* O111:B4 can inhibit P1(S’) transduction, which demonstrated that P1(S’) could adsorb to LPS with these O-antigens. Taken together, this study fills the knowledge gap of P1 transduction with its alternative S’ tail fibre, with evidence supporting the potential use of P1(S’) to improve the transduction efficiency of a sequence-specific Cas9 antimicrobial system in *S. flexneri*, which is a clinically relevant human pathogen.

## RESULTS

### *S. flexneri* 2a 2457O, *S. flexneri* 5a M90T and *E. coli* O3 supported P1(S’) transduction of phagemid DNA

We have established a J72114 P1 phagemid system in Δ*pac E. coli* K12 EMG16 P1kc lysogenic strain that allows the packaging of phagemid DNA into infectious P1 phage particles upon arabinose induction (**Supplementary Figures 1a**, **1b**, **1c**). The chloramphenicol acetyltransferase *(cat)* gene of the phagemid allows quantification of transduction events by enumerating the chloramphenicol resistant colony forming units (Cm^R^ CFU) recovered after P1 infection. The P1 phagemid contains a constitutively expressed *cas9* system with a DNA sequence-specific antimicrobial effect in the presence of a chromosomal-targeting crRNA guide (Bikard et al., 2014; Citorik et al., 2014; Gomaa et al., 2014; Cui and Bikard, 2016) (**Supplementary Figure 1d**). Our preliminary results suggested that P1 can transduce the *cas9* phagemid into *E. coli* K12 NCM3722, *S. flexneri* 2a 2457O and *S. flexneri* 5a M90T at a high efficiency, giving at least 10^9^ phagemid transductants per mL lysate (TU/mL) (data not shown). Since the induction of a P1 lysogen is expected to produce a mixed population of P1 progeny with either the S or S’ tail fibre, we sought to compare the transduction efficiency of P1(S) and P1(S’) on different strains of *E. coli* and *S. flexneri*. The aim was to identify the P1 tail fibre that could give a higher transduction efficiency, particularly in targeting different strains of *S. flexneri*. P1 *cas9* phagemid with a randomly generated spacer sequence (non-chromosomal targeting, *cas9*-NT) was used for the preparation of lysates, as the phagemid can be stably maintained in *E. coli* and *S. flexneri*, which allowed quantification of phagemid transductants.

To obtain a homogeneous population of P1 phage particles with either the S or S’ tail fibre, the coding sequences for *Sc, Sv*’, *U*, *U*’, *Sv*’ and *cin* recombinase were deleted via the lambda Red recombineering system (**Figures 1a, b**). The Δ*S-cin* mutation was then complemented *in trans*, by transforming the mutant strain with a low-copy number plasmid containing genes encoding the S tail fibre and U chaperone (plasmid termed pP1SU) or the S’ tail fibre and U’ chaperone (plasmid pP1S’U’) (**Figure 1b**). Δ*pac* EMG16 cells co-transformed with a *cas9*-NT phagemid and the pP1SU or pP1S’U’ plasmid were used for phagemid lysate preparation. The phagemid lysates were then used to treat *E. coli* K12 NCM3722, *E. coli* BL21, *E. coli* 24377A, *E. coli* O3, *S. flexneri* 2a 2457O and *S. flexneri* 5a M90T, and the number of phagemid transductants recovered were quantified. Infection of *E. coli* K12 NCM3722 and *E. coli* B BL21 with P1(S) but not P1(S’) yielded Cm^R^ CFU (**Figure 1c**). The lack of phagemid transductant indicated that P1(S’) could not transduce phagemid DNA into *E. coli* K12 and *E. coli* B, which was consistent with results from previous studies (Lennox, 1955; Iida et al., 1982; Iida, 1984; Sandulache et al., 1984). On the contrary, infection of *E. coli* O3 with P1(S’) but not P1(S) yielded Cm^R^ CFU, indicating that *E. coli* O3 was permissive for P1(S’) transduction yet resistant to P1(S) transduction. The lack of phagemid transductant was not due to cell death after P1 infection, as we could recover approximately 5 X 10^8^ to 1 X 10^9^ *E. coli* BL21, *E. coli* NCM3722 and *E. coli* O3 CFU (per mL reaction mix) on non-selective LB (data not shown). P1(S’) infection of *S. flexneri* 5a M90T and *S. flexneri* 2a 2457O yielded a 207-fold (*s* = 31, *n*= 12, p < 0.005) and a 1.61-fold (*s*= 0.24, *n*= 12, p > 0.05) increase in the number of phagemid transductant respectively, when compared to P1(S) infection of these bacteria. P1(S’) and P1(S) infection of *E. coli* E24377A yielded a similar number of phagemid transductant, which was approximately 1 x 10^6^ transductants per mL lysate (TU/mL) (*s*= 2.6 X 10^5^ TU/mL, *n*= 24, p > 0.05). To verify that differences in the number of P1(S) and P1(S’) phagemid transductant were not due to variations in the titre of phagemid transducing particles, all phagemid lysates were treated with DNase I and quantified for P1-packaged phagemid DNA via quantitative PCR (qPCR). qCPR quantification of the *cas9* DNA sequence in DNase I-treated lysates indicated a similar concentration of phagemid DNA that were packaged into phage particles (**Supplementary Figures 2a, 2b**). Since all lysates contained a similar concentration of P1-packaged phagemid DNA, differences in the number of *S. flexneri* and *E. coli* phagemid transductants were indicative of variations in the transduction efficiency of phagemid DNA.

**Figures 1:**
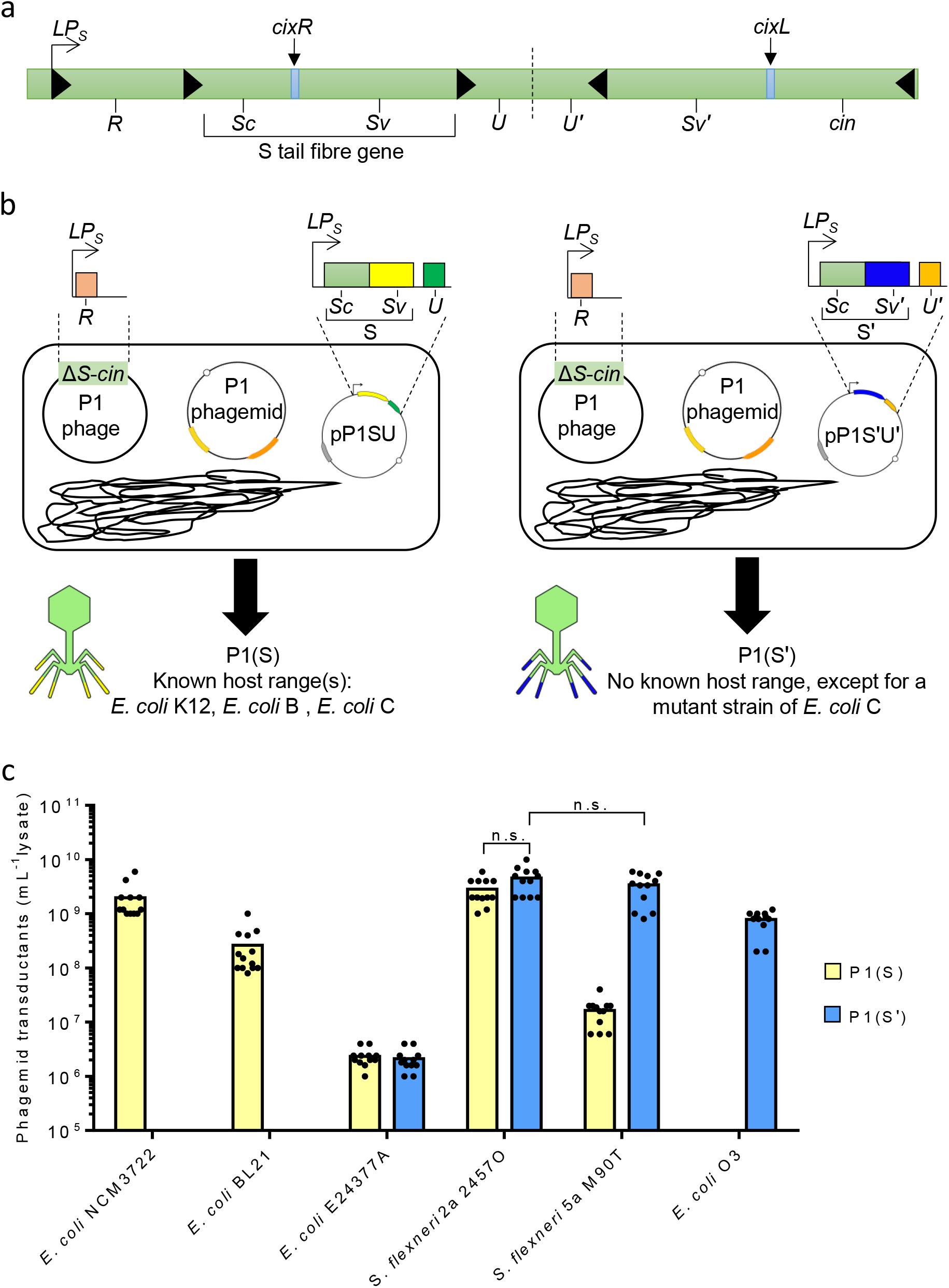
*S. flexneri* 2a 2457O, *S. flexneri* 5a M90T and *E. coli* O3 supported P1(S’) transduction of phagemid DNA. (**a**) The organisation of P1 tail fibre operon. Black triangles mark the start site for the coding sequences. The expression of *R* (hypothetical tail assembly protein), *S* (tail fibre protein) and *U* (hypothetical chaperone protein) are controlled by the P1 late promoter, LP_*Sit*_. The tail fibre is divided into a constant Sc region as well as a variable Sv region. The coding sequence for the alternative variable region, *Sv*’, and its chaperone, *U* are located adjacent to *Sv* and *U*, in an opposite orientation. Cin recombinase mediates inversion of the C segment at the *cix* sites (in blue), allowing expression of the *S*’ tail fibre and *U* chaperone. (**b**) Schematic diagram showing the packaging of a phagemid into P1 phage particles using the Δ*S-cin E. coli* EMG16 P1 lysogen. The tail fibre operon and the *cin* recombinase coding sequence were deleted. The Δ*S-cin*mutation was complemented *in trans*, by providing either a combination of *S* and *U* (plasmid termed pP1SU) or *S*’ and *U* (plasmid termed pP1S’U’) genes, controlled by the P1 late promoter LP_*Sit*_. The lysates prepared is expected to contain a homogenous population of P1 phage with either S (P1(S)) or S’ (P1(S’)) tail fibre. P1(S) (with yellow tail fibre terminals) was shown to transduce strains of *E. coli* K12, *E. coli* B, *E. coli* C, while the host range of P1(S’) (with blue tail fibre terminals) was not determined (Lennox, 1955; Iida et al., 1982; Iida, 1984; Sandulache et al., 1984). A mutant strain of *E. coli* C could support transduction by both the P1(S) and P1(S’), although the nature of mutation was not identified (Iida, 1984). (**c**) The number of phagemid transductant recovered, after P1(S) (yellow bars) and P1(S’) (blue bars) phagemid lysates treatment of *E. coli* K12 NCM3722, *E. coli* BL21, *E. coli* E24377A, *S. flexneri* 2a 2457O, *S. flexneri* 5a M90T and *E. coli* O3. The p-values were determined by Welch’s ANOVA. n.s. represented no significant difference(s) in p-values for comparison of the number of *S. flexneri* 2457O phagemid transductant recovered between P1(S) and P1(S’) infection, as well as comparison between the number of *S. flexneri* 2a 2457O and *S. flexneri* 5a M90T phagemid transductant recovered after P1(S’) infection.

Taken together, P1(S’) transduced phagemid DNA into *S. flexneri* 5a M90T, *S. flexneri* 2a 2457O and *E. coli* O3 at a high efficiency and therefore identified the host range of P1(S’) which was not determined in previous studies.

### P1(S’) transduction has a less stringent requirement for Ca^2+^ compared to P1(S)

Historic protocol for generalised P1 transduction stated that free calcium ions are needed for P1 phage adsorption and transduction (Lennox, 1955). To verify the role of Ca^2+^ in P1 transduction, the transduction efficiency of P1(S’) and P1(S) were assessed in the absence of Ca^2+^ or with the addition of citrate which sequesters free Ca^2+^. The relative transduction efficiency of P1(S’) and P1(S) were compared to that of phage infection in PLM medium, in which Ca^2+^ was present during the phage adsorption stage. The absence of Ca^2+^ or the addition of citrate during phage adsorption stage completely blocked P1(S) transduction of *S. flexneri* 5a M90T at both multiplicity of infection (MOI) of 0.1 and 0.01 (**Supplementary Figures 3a, 3b**). At a MOI of 0.1, the lack of Ca^2+^ supplement or the addition of citrate caused an average 278-fold reduction (-Ca^2+^, 264-fold, *s*= 22, *n*= 9; -Ca^2+^, + Citrate, 293-fold, *s*= 31, *n*= 9) in the transduction efficiency of P1(S) in *S. flexneri* 2a 2457O. Similarly, the absence of Ca^2+^ supplement or the addition of citrate reduced the transduction efficiency of P1(S) in *E. coli* K12 NCM3722 by approximately 22-fold (*s*= 3.0, *n*= 9, p < 0.0005) and 267-fold (*s*= 27.8, *n*= 9, p < 0.0005) respectively, at a MOI of 0.1.

Contrastingly, the transduction efficiency of P1(S’) in *E. coli* O3 was not significantly affected by the lack of Ca^2+^ nor the presence of citrate at both MOI of 0.1 and 0.01 (p > 0.05) (**Supplementary Figures 3a, 3b**). Neither the presence of citrate nor the lack of Ca^2+^ supplement caused significantly changes to the transduction efficiency of P1(S’) in *S. flexneri* 5a M90T and *S. flexneri* 2457O, at a MOI of 0.1 (p > 0.05) (**Supplementary Figures 3a)**. The addition of citrate at a lower MOI of 0.01, however, significantly reduced the transduction efficiency of P1(S’) in *S. flexneri 5a* M90T and *S. flexneri* 2a 2457O by approximately 9.8-fold (*s*= 2.6, *n*= 9, p < 0.0005) and 2.1-fold (*s*= 2.1, *n*=9, p < 0.005) respectively (**Supplementary Figures 3b)**. Similarly, at a MOI of 0.01, the lack of Ca^2+^ or the addition of citrate caused at least 50-fold reduction in P1(S) transduction efficiency in S*. flexneri* 2a 2457O (-Ca^2+^, 921-fold, *s*= 134, *n*= 9; -Ca^2+^, + Citrate, 2447fold, *s*= 616, *n*= 9) and *E. coli* NCM3722 (-Ca^2+^, 72-fold, *s*= 6.7, *n*= 9; -Ca^2+^, + Citrate, 216-fold, *s*= 72, *n*= 9).

Taken together, a lower reduction in P1(S’) transduction efficiency without Ca^2+^ or in the presence of citrate indicated that P1(S’) transduction has a less stringent requirement for Ca^2+^ when compared to P1(S).

### P1(S’) transduction of phagemid DNA required O-antigen

Since the host range and the receptor of P1(S’) were not known, we sought to identify the factor(s) required for P1(S’) transduction of phagemid DNA into *S. flexneri* 2a 2457O, *S. flexneri* 5a M90T and *E. coli* O3. Previous data indicated that only *E. coli* and *S. flexneri* strains with O-antigen supported P1(S’) transduction (**Figure 1c**). To determine if the O-antigen of *S. flexneri* and *E. coli* O3 were required for P1(S) and P1(S’) transduction, the relative transduction efficiency of both phages was assessed on wildtype (WT) and O-antigen deficient mutant strains of *S. flexneri* 2a 2457O, *S. flexneri* 5a M90T and *E. coli* O3. The genes selected for generating the O-antigen deficient mutant strains were *waaL, rfe* and *rfc/wzy* (**Figure 2a**). *waaL* and *rfe* genes encode the O-antigen ligase and the WecA integral membrane protein, which are required for the ligation of O-antigen onto the LPS core (Yao and Valvano, 1994) and the biosynthesis of *E. coli* as well as *S. flexneri* O-antigens containing a *N*-acetylglucosamine residue (Alexander and Valvano, 1994), respectively. Δ*waaL* and Δ*rfe* strains of *S. flexneri* (Yao and Valvano, 1994; Sandlin et al., 1995; West et al., 2005) and *E. coli* (Yao and Valvano, 1994; Alexander and Valvano, 1994) were shown to give a rough type LPS without any O-antigen. On the contrary, the gene product of *rfc/wzy* is required for the polymerisation of O-antigen polysaccharide and a loss-of-function mutation in the *rfc/wzy* gene of *S. flexneri* (Nath et al., 2015; Kim et al., 2018) as well as *E. coli* O3 (Ren et al., 2008) were shown to give a semi-rough type LPS that has only one O-antigen unit. Δ*waaL*, Δ*rfe* and Δ*rfc/wzy* strains of *S. flexneri* 2a 2457O, *S. flexneri* 5a M90T and *E. coli* O3 were generated via lamda Red recombineering, and the effects of these mutations on the bacterial LPS were visualised via silver staining. LPS extracted from WT cells gave a ladder-like pattern of multiple bands on the silver-stained gel, indicating the presence of O-antigen repeating units of varying lengths (**Figures 2b, 2c**). LPS extracted from the Δ*waaL* and Δ*rfe* mutants gave a single band, indicating the absence of O-antigen. The LPS of the Δ*rfc* mutants gave two bands on the silver-stained gel, in which one was a unit higher than the bands observed for Δ*waaL* and Δ*rfe* mutants, which verified the semi-rough type LPS of Δ*rfc* mutants with a single O-antigen unit.

**Figures 2:**
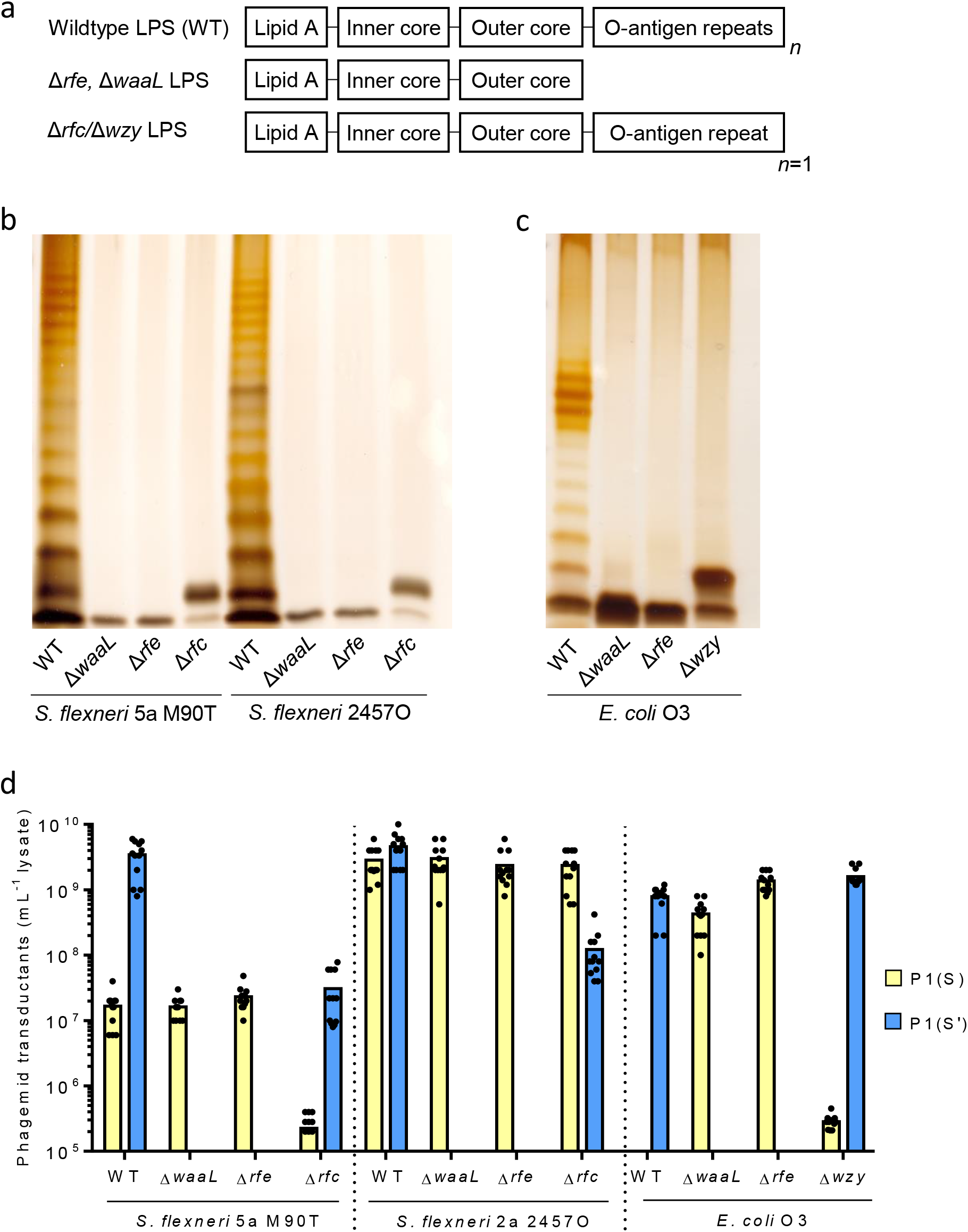
P1(S’) transduction of *S. flexneri* 5a M90T, *S. flexneri* 2a 2457O and *E. coli* O3 required O-antigen. (**a**) Schematic diagram showing the expected LPS structure of wildtype (WT), Δ*rfe, ΔwaaL* and Δ*rfc* strains of *S. flexneri* 5a M90T, *S. flexneri* 2a 2457O and *E. coli* O3. The LPS of Δ*rfe* and Δ*waaL* mutants contains Lipid A, inner and outer core oligosaccharide structures without any O-antigen unit(s), while that of Δ*rfc* mutant has only 1 O-antigen unit compared to the multiple O-antigen repeating units on the LPS of WT cells. Analysis of LPS extracted from wildtype (WT), Δ*waaL*, Δ*rfe* and Δ*rfc* strains of (**b**) *S. flexneri* 5a M90T (*S. flexneri* M90T), *S. flexneri* 2a 2457O (*S. flexneri* 2457O) and (**c**) *E. coli* O3. LPS samples were analysed on 12 % precast polyacrylamide gels and silver stained. LPS of WT strains of *S. flexneri* 5a M90T, *S. flexneri* 2a 2457O and *E. coli* O3 showed a ladder-like pattern characteristic of smooth type LPS, while the LPS of the Δ*waaL* and Δ*rfe* mutants gave a single band characteristic of rough type LPS without any O-antigen. LPS of the Δ*rfc* mutants gave two bands, indicating semi-rough type LPS consisting of lipid A, complete oligosaccharide core and a single O-antigen unit. (**d**) The number of phagemid transductant recovered, after P1(S) (yellow bars) and P1(S’) (blue bars) phagemid lysates treatment of wildtype (WT), Δ*waaL*, Δ*rfe* and Δ*rfc* strains of *S. flexneri* 5a M90T, *S. flexneri* 2a 2457O and *E. coli* O3.

To determine the relative transduction efficiency of phagemid in WT, Δ*waaL*, Δ*rfe* and Δ*rfc/wzy* strains of *S. flexneri* 2a 2457O, *S. flexneri* 5a M90T and *E. coli* O3, the bacteria cells were treated with either P1(S) or P1(S’) phagemid lysates and the number of phagemid transductants was enumerated. No phagemid transductant was recovered after P1(S’) infection of the Δ*waaL* and Δ*rfe* mutants (**Figure 2d**). The lack of phagemid transductant was not due to cell death after P1(S’) infection, as we recovered approximately 1 X 10^8^ to 5 X 10^8^ CFU (per mL reaction mix) on non-selective LB (data not shown). Complementation of the Δ*waaL* and Δ*rfe* mutations *in trans*, by providing a *waaL* and a *rfe* gene respectively, restored the smooth type LPS as well as the recovery of phagemid transductants after P1 (S’) infection of the Δ*waaL* and Δ*rfe* mutants (**Supplementary Figures 4a, 4b, 4c**). On the contrary, P1(S’) infection of Δ*rfc S. flexneri* 5a M90T and Δ*rfc S. flexneri* 2a 2457O yielded a 216-fold (*s*= 44, *n*= 12, p < 0.0005) and 58-fold (*s*= 9.8, *n*= 12, p < 0.0005) lower number of phagemid transductant respectively, when compared to P1(S’) infection of WT *S. flexneri* (**Figure 2d**). P1(S’) infection of Δ*rfc E. coli* O3 yielded a 2-fold higher (*s*= 0.18, *n*= 12, p < 0.0005) number of phagemid transductant when compared to P1(S’) infection of *E. coli* O3.

There were no significant differences in the number of phagemid transductant recovered between P1(S) infection of WT, Δ*waaL* and Δ*rfe* strains of *S. flexneri* 2a 2457O and *S. flexneri* 5a M90T (p > 0.05) (**Figure 2d**). Although P1 (S) infection of WT and Δ*rfc S. flexneri* 2a 2457O yielded a similar number of phagemid transductant (p > 0.05), P1(S) infection of the Δ*rfc S. flexneri* 5a M90T yielded a 321-fold (*s*= 148, *n*= 12, p < 0.005) lower number of phagemid transductant when compared to infection of WT *S. flexneri* 5a M90T. While no phagemid transductant was recovered after P1(S) infection of WT *E. coli* O3, P1(S) infection of its isogenic Δ*waaL* and Δ*rfe* mutants yielded approximately 4.3 X10^8^ (*s*= 6.6 X 10^7^, *n*=12) and 1.3 X 10^9^ (*s*= 1.8 X 10^8^, *n*=12) phagemid transductants per mL lysate, respectively. Δ*rfc E. coli* O3, however, was permissive for P1(S) transduction yet the number of phagemid transductant was approximately 1500-fold (*s*= 93, *n*= 12, p < 0.0005) and 5000-fold (*s*= 5000, *n*= 297, p < 0.0005) lower, when compared to that recovered after P1(S) infection of its isogenic Δ*waaL* and Δ*rfe* mutants, respectively.

Overall, these data indicated that P1(S’) transduction of *S. flexneri* 2a 2457O, *S. flexneri* 5a M90T and *E. coli* O3 required at least one O-antigen unit while the O-antigen deficient mutants were still susceptible to P1(S) transduction.

### P1(S’) transduction of *S. flexneri* 2a 2457O and *S. flexneri* 5a M90T was not significantly affected by mutations in the O-antigen modification genes

All known serotypes of *S. flexneri*, except for serotype 6, share the same O-antigen backbone with modifications such as glucoslylation and acetylation, which give rise to variation in serotypes (Kenne et al., 1978; Foster et al., 2011; Perepelov et al., 2012). Despite the variations in O-antigen modifications, P1(S’) could transduce phagemid DNA into *S. flexneri* 2a 2457O and *S. flexneri* 5a M90T at a similar efficiency (**Figure 1c**). We sought to determine if changes to the O-antigen modifications of *S. flexneri* would affect P1(S’) and P1(S) transduction efficiency. We have generated 8 mutant strains of *S. flexneri* 2a 2457O and *S. flexneri* 5a M90T that were defective in combinations of O-antigen acetylation and glycosylation, by deleting the O-antigen modification gene(s) via lamda Red recombination (**Figure 3a**) (West et al., 2005; Teh et al., 2020). The effects, if any, of these mutations on the LPS structures of *S. flexneri* were visualised by silver stain. All the LPS samples showed a ladder-like pattern on the silver-stained gel, indicating the presence of O-antigen repeating units (**Supplementary Figure 5a**). Only the LPS of Δ*gtr* mutants lacking a glucose modification showed shift(s) in the band size(s) when compared to the LPS of WT cells (West et al., 2005; Teh et al., 2020).

**Figures 3:**
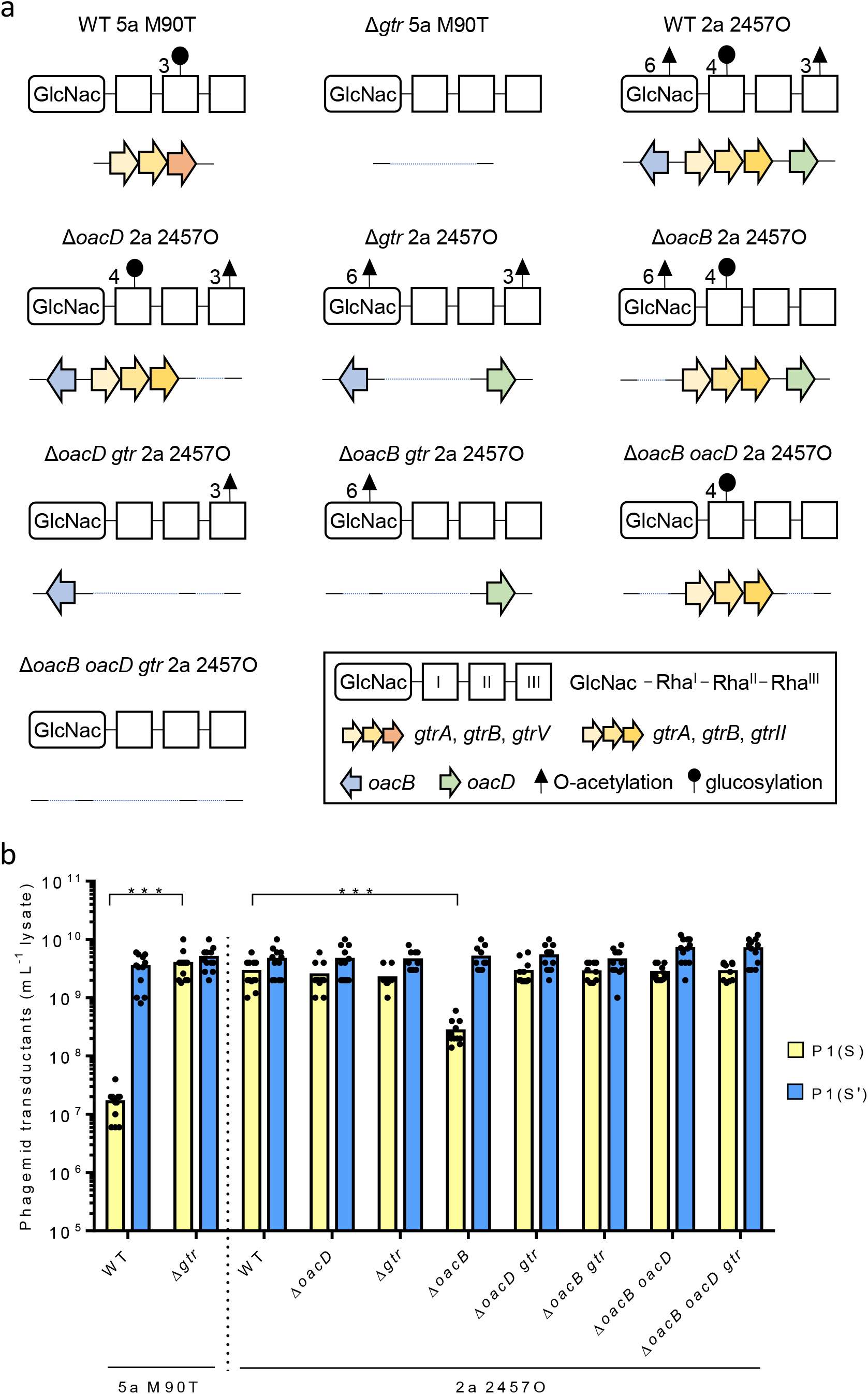
P1(S’) transduction of *S. flexneri* 5a M90T and *S. flexneri* 2a 2457O was not significantly affected by mutations in the O-antigen modification genes. (**a**) Schematic diagrams showing isogenic mutant strains of *S. flexneri* 5a M90T (5a M90T) and *S. flexneri* 2a 2457O (2a 2457O) with the expected change(s) to their glucosylation and/or acetylation of O-antigen, based on the respective genetic deletion(s) (West et al., 2005; Teh et al., 2020). All serotype of *S. flexneri*, except for serotype 6, share a common O-antigen backbone comprised of repeating units of three L-rhamnose residues (Rha^I^-Rha^II^-Rha^III^) and a *N*-acetylglucosamine residue (GlcNac). In terms of *S. flexneri* 2a 2457O and *S. flexneri* 5a M90T, differences in the acetylation and glucosylation of O-antigen give rise to a variation in their serotyping. The genes involved in the acetylation and glucosylation of *S. flexneri* 2a 2457O and *S. flexneri* 5a M90T O-antigens were indicated as arrows (West et al., 2005; Teh et al., 2020). The orientation of the glycosidic linkages between the acetyl and glucose molecules and the O-antigen backbone were shown as Arabic numerals. Blue dotted lines represented genetic deletion(s). (**b**) The number of phagemid transductants recovered after P1(S) (yellow bars) and P1(S’) (blue bars) phagemid lysates treatment of wildtype (WT) and all 8 mutant strains of *S. flexneri* 5a M90T (5a M90T) and *S. flexneri* 2a 2457O (2a 2457O). The p-values were determined by Welch’s ANOVA. p < 0.0005 was indicated as as ***, for comparison of the number of phagemid transductants recovered between P1(S) infection of wildtype and Δ*gtr S. flexneri* 5a M90T, as well as the number of phagemid transductants recovered between P1(S) infection of wildtype and Δ*oacB S. flexneri* 2a 2457O.

WT and all 8 mutant strains of *S. flexneri* 2a 2457O and *S. flexneri* 5a M90T were treated with P1(S) and P1(S’) phagemid lysate and the number of phagemid transductant was enumerated. P1(S’) phagemid lysate treatment of WT and all 8 mutant strains of *S. flexneri* yielded a similar number of phagemid transductant (p > 0.05) (**Figure 3b**). The similar number of P1(S’) phagemid transductant indicated that P1(S’) transduction of *S. flexneri* 2a 2457O and *S. flexneri* 5a M90T was not significantly affected by mutations in the O-antigen modification genes. Contrastingly, P1(S) infection of Δ*gtr S. flexneri* 5a M90T yielded a 235-fold (*s*= 40, *n*= 12, p < 0.0005) higher number of phagemid transductant when compared to P1(S) infection of WT *S. flexneri* 5a M90T (**Figure 3b**). Δ*oacB* S*. flexneri* 2a 2457O mutant yielded a 12-fold (*s*= 1.4, *n*= 12, p < 0.0005) lower number of phagemid transductant after P1(S) infection, when compared to the infection of WT S*. flexneri* 2a 2457O. Complementation of the Δ*oacB* mutation *in trans* increased the number of phagemid transductant recovered after P1(S) infection (**Supplementary Figure 5b**). Unexpectedly, P1(S) infection of Δ*oacB gtr, ΔoacB oacD* and Δ*oacB oacD gtr* strains of *S. flexneri* 2a 2457O yielded a similar number of phagemid transductant when compared to P1(S) infection of WT *S. flexneri* 2a 2457O (p > 0.05) (**Figure 3b,** refer to **Supplementary Materials 2** for p-values).

Overall, the results indicated that P1(S’) transduction of phagemid DNA was not significantly affected by mutations in the O-antigen modifications genes of *S. flexneri* 2a 2457O and *S. flexneri* 5a M90T, when compared to P1(S) transduction.

### The LPS of *S. flexneri* 2a 2457O, *S. flexneri* 5a M90T and *E. coli* O3 with at least one O-antigen unit could act as an inhibitor against P1(S’) transduction

P1(S’) transduction of *S. flexneri* 2a 2457O, *S. flexneri* 5a M90T and *E. coli* O3 required the presence of at least one O-antigen unit (**Figure 2d**). We therefore hypothesised that the O-antigens of *S. flexneri* 2a 2457O, *S. flexneri* 5a M90T and *E. coli* O3 might be important for P1(S’) adsorption. If this is true, an excess *S. flexneri* 2a 2457O, *S. flexneri* 5a M90T and *E. coli* O3 LPS in phagemid lysates could sequester P1(S’) phage and this might reduce its transduction efficiency. To verify this hypothesis, LPS extracted from WT and O-antigen deficient mutant strains of *S. flexneri* 2a 2457O, *S. flexneri* 5a M90T and *E. coli* O3 were assessed for their inhibitory effect against P1(S) and P1(S’) transduction (**Figure 4a**).

**Figure 4:**
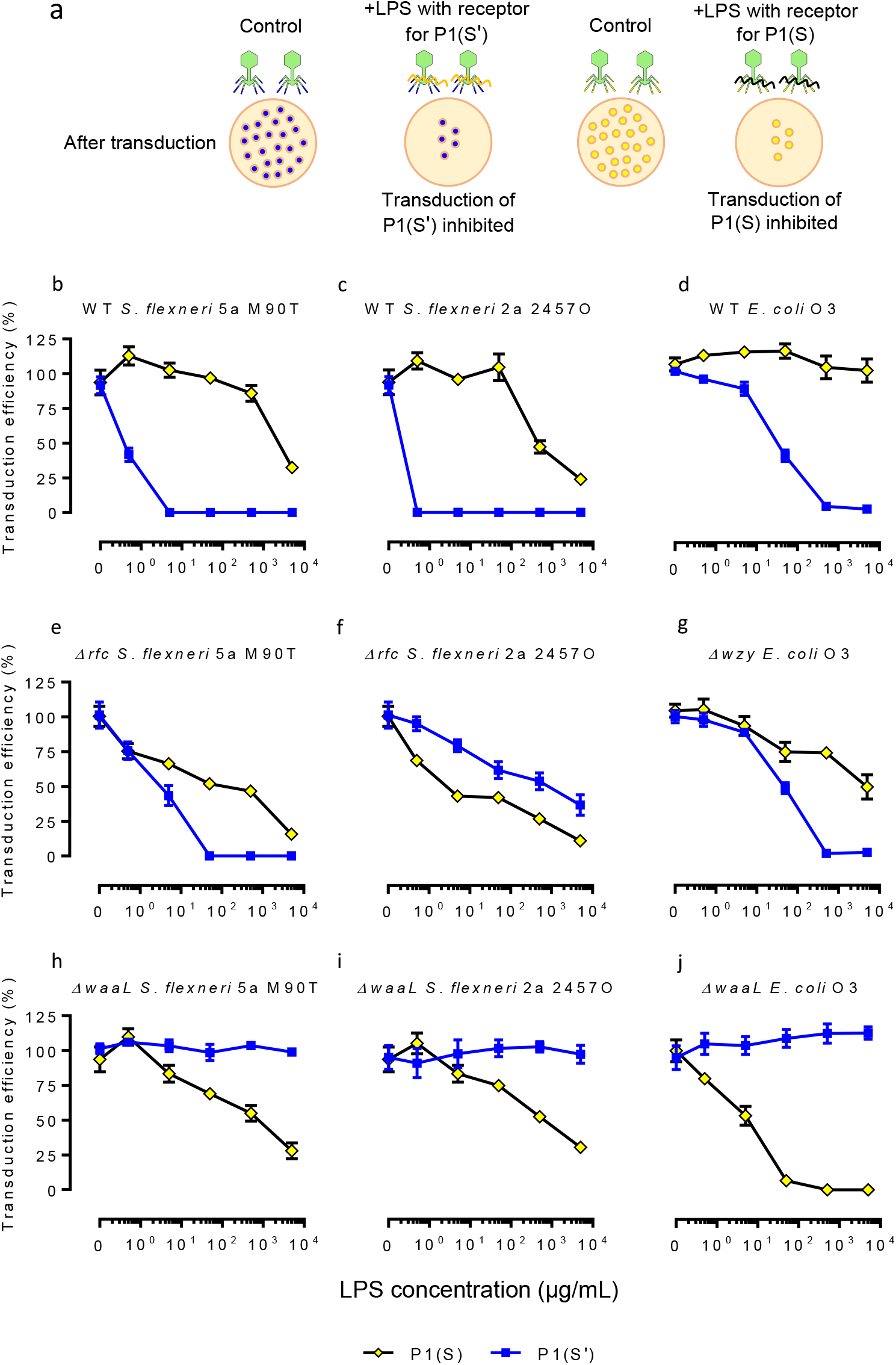
The LPS of *S. flexneri* 2a 2457O, *S. flexneri* 5a M90T and *E. coli* O3 with at least one O-antigen unit could act as an inhibitor against P1(S’) transduction. (**a**) Schematic diagram showing a LPS inhibition assay. When applied at a certain concentration, LPS that allows the adsorption of P1(S) and/or P1(S’) would sequester the phage thus reduce its relative transduction efficiency on *E. coli* K12 NCM3722 (yellow) and *S. flexneri* 2a 2457O (blue), respectively (Sandulache et al., 1984). A reduced transduction efficiency would give a lower number of phagemid transductant when compared to that recovered with non-LPS treated phagemid lysates. The transduction efficiency of P1(S) (yellow diamonds, black lines) and P1(S’) (blue squares, blue lines) after pre-incubation with 0.5 μg/mL, 5.0 μg/mL, 50 μg/mL, 500 μg/mL and 5000 μg/mL of LPS extracted from (**b**) WT *S. flexneri* 5a M90T, (**c**) WT *S. flexneri* 2a 2457O, (**d**) WT *E. coli* O3, (**e**) Δ*rfc S. flexneri* 5a M90T, (**f**) Δ*rfc S. flexneri* 2a 2457O, (**g**) Δ*wzy E. coli* O3 2457O, (**h**) Δ*waaL S. flexneri* 5a M90T, (**i**) Δ*waaL S. flexneri* 2a 2457O, (**j**) Δ*waaL E. coli* O3. All the graphs share the same y-axis for transduction efficiency (%) and an x-axis for LPS concentration (μg/mL). Each data point represented results generated using 1 LPS sample, 1 lysate sample, 3 biological repeats (host cells used) and 2 technical repeats. Data were represented as mean ± SEM.

LPS extracted from WT *S. flexneri* 5a M90T and WT *S. flexneri* 2a 2457O completely blocked P1(S’) transduction when applied at concentrations greater or equal to 5 μg/mL (p < 0.0005) and 0.5 μg/mL (p < 0.0005), respectively (**Figures 4b, 4c**). The LPS of Δ*rfc S. flexneri* 5a M90T completely blocked P1(S’) transduction at concentrations greater or equal to 50 μg/mL (p < 0.0005), while 5000 μg/mL of Δ*rfc S. flexneri* 2a 2457O LPS was required to reduce the transduction efficiency of P1(S’) by approximately 64 % (*s*= 7.3, *n*= 6, p < 0.0005) (**Figures 4e, 4f**). 500 μg/mL of WT and Δ*wzy E. coli* O3 LPS reduced P1(S’) transduction efficiency by approximately 94 % (WT *E. coli* O3, *s*= 2.6, *n*= 6, p < 0.0005) and 96 % (Δ*wzy E. coli* O3, *s*= 1.8, *n*=6, p < 0.0005), respectively (**Figures 4d, 4g**). Overall, the inhibitory effect of LPS with at least one O-antigen unit indicated that P1(S’) might adsorb to the LPS of *S. flexneri* 5a M90T, *S. flexneri* 2a 2457O and *E. coli* O3 with at least one O-antigen unit. Contrastingly, treatment of P1(S’) lysate with the LPS of Δ*waaL* mutants yielded an average transduction efficiency of 107 % at all concentrations *(ΔwaaL S. flexneri* M90T, *s*= 3.4, *n* = 5, p > 0.05; Δ*waaL S. flexneri* 2457O, *s*= 4.9, *n* = 30, p > 0.05; Δ*waaL E. coli* O3, *s*= 5.7, *n* = 30, p > 0.05) (**Figures 4h, 4i, 4j**). Similarly, treatment of P1(S’) lysate with *E. coli* K12 LPS yielded an average transduction efficiency of 105 % (*s*= 6.1, *n*= 30, p > 0.05) at all concentrations (**Supplementary Figure 6**). Therefore, the lack of inhibitory effect against P1(S’) transduction indicated that P1(S’) might not adsorb to the LPS of *E. coli* K12 as well as the LPS of the Δ*waaL* mutant strains without O-antigen, which was consistent with the absence of phagemid transductant after P1(S’) infection of these bacteria (**Figures 1c, 2d**).

5 μg/mL of *E. coli* K12 LPS reduced P1(S) transduction efficiency by approximately 76 % (*s*= 3.3, *n*= 6, p < 0.0005), while concentrations greater or equal to 50 μg/mL completely blocked P1(S) transduction **Supplementary Figure 6**). 500 μg/mL to 5000 μg/mL of LPS extracted from all strains of *S. flexneri* reduced the transduction efficiency of P1(S) by at least 50 % (**Figures 4b, 4c, 4e, 4f, 4h, 4i**). The inhibitory effect of *S. flexneri* and *E. coli* K12 LPS against P1(S) transduction indicated phage adsorption to these LPS, which was consistent with the recovery of phagemid transductants after P1(S) infection of WT and O-antigen deficient mutant strains of *S. flexneri*, as well as *E. coli* K12 NCM3722 (**Figure 2d**).

5 μg/mL to 50 μg/mL Δ*rfc S. flexneri* 2a 2457O LPS reduced P1(S) transduction efficiency by approximately 57 % (*s*= 2.0, *n*= 12, p < 0.0005), indicating a greater inhibitory effect of the LPS against P1(S) transduction when compared to the LPS of WT *S. flexneri* 2a 2457O (**Figure 4f**). Treatment of P1(S) lysates with WT *E. coli* O3 LPS yielded an average transduction efficiency of 110 % at all concentrations tested (*s*= 7.5, *n*= 30, p > 0.05), while 5000 μg/mL of Δ*wzy E. coli* O3 LPS was needed to reduce P1(S) transduction efficiency by approximately 54 % (s= 6.2, n= 6, p < 0.0005) (**Figures 4d, 4g**). Contrastingly, 5 μg/mL of Δ*waaL E. coli* O3 LPS reduced P1(S) transduction efficiency by 47 % (s= 4.8, n= 6, p < 0.0005), followed by complete inhibition of its transduction at concentrations greater or equal to 500 μg/mL (**Figure 4j**). Differences in the inhibitory effect of *E. coli* O3 LPS against P1(S) transduction suggested a greater adsorption efficiency of P1(S) to the LPS of Δ*waaL E. coli* O3 when compared to the LPS of WT *E. coli* O3 and its isogenic Δ*wzy* mutant. A greater adsorption efficiency of P1(S) on Δ*waaL E. coli* O3 was consistent with a significantly higher number of phagemid transductants recovered after P1(S) infection of the mutant strain, when compared to that recovered after P1(S) infection of WT and its isogenic Δ*wzy* strains (**Figure 2d**).

Taken together, the LPS of *S. flexneri* 2a 2457O, *S. flexneri* 5a M90T and *E. coli* O3 with at least one O-antigen unit produced inhibitory effects against P1(S’) transduction, indicating that these LPS allowed P1(S’) phage adsorption.

### *E. coli* O111:B4 LPS blocked P1(S’) transduction completely

*S. flexneri* and *E. coli* O3 strains with at least one O-antigen unit were susceptible to P1(S’) transduction, and the LPS of these bacteria allowed P1(S’) adsorption. The LPS inhibiton assay could potentially identify other hosts of P1(S’) and P1(S), by identifying LPS samples that showed neutralising activity against P1(S’) and P1(S) transduction. Hence, we sought to test if P1(S) and P1(S’) could adsorb to the smooth LPS of other *E. coli* strains, by repeating the LPS inhibition assay with *E. coli* O26:B6, *E. coli* O55:B5, *E. coli* O111:B4 and *E. coli* O127:B8 LPS (refer to **Supplementary Figures 7** for the chemical structures of LPS).

5000 μg/mL of *E. coli* O26:B6 and *E. coli* O55:B5 LPS reduced P1(S’) transduction efficiency by 76 % (*s*= 1.4, *n*= 6, p < 0.0005) and 50 % (*s*= 3.4, *n*= 6, p < 0.0005) respectively (**Figures 5a, 5b**). 5 μg/mL to 5000 μg/mL of *E. coli* O127:B8 LPS caused an average 23 % (*s*= 1.6, *n*= 24, p < 0.05) reduction in P1(S’) transduction efficiency (**Figure 5d**). On the contrary, *E. coli* O111:B4 LPS completely blocked P1(S’) transduction when applied at concentrations greater or equal to 5 μg/mL (p < 0.0005) (**Figure 5c**). The high inhibitory effect of *E. coli* O111:B4 LPS was similar to that of WT *S. flexneri* 2a 2457O and WT *S. flexneri* 5a M90T LPS, indicating that P1(S’) could adsorb to these LPS at a high efficiency (**Figures 4b, 4c**). Treatment of P1(S) lysates with *E. coli* O111:B4 and *E. coli* O127:B8 LPS yielded an average transduction efficiency of 105 % at all concentrations (*E. coli* O111:B4, *s*= 4.8, *n*= 30, p > 0.05; *E. coli* O127:B8, *s*= 5.6, *n*= 30, p > 0.05) (**Figures 5c, 5d**). 5000 μg/mL of *E. coli* O26:B6 and *E. coli* O55:B5 LPS reduced P1(S) transduction efficiency by approximately 93 % (*s*= 2.0, *n*= 6, p < 0.0005) and 38 % (*s*= 1.7, *n*= 6, p < 0.0005) respectively (**Figures 5a, 5b**). A higher inhibitory effect of *E. coli* O26:B6 LPS against P1(S) transduction indicated a greater P1(S) adsorption efficiency to this LPS when compared to the LPS of *E. coli* O55:B5, *E. coli* O111:B4 and *E. coli* O127:B8.

**Figures 5:**
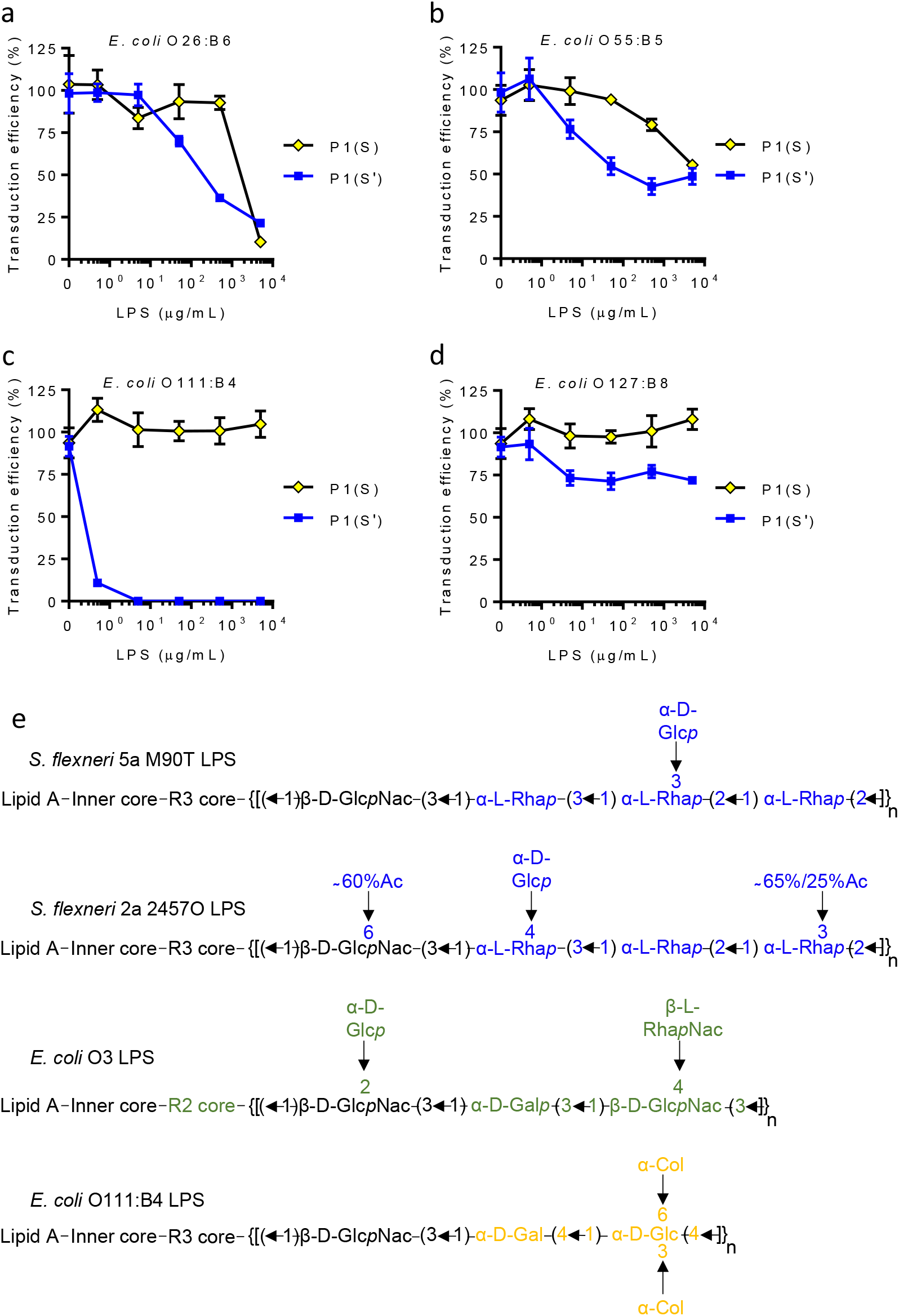
Low concentration of *E. coli* O111:B4 LPS blocked P1(S’) transduction completely. Quantification of P1(S) (yellow diamonds, black lines) and P1(S’) (blue squares, blue lines) transduction efficiency when pre-incubated with 0.5 μg/mL, 5.0 μg/mL, 50 μg/mL, 500 μg/mL and 5000 μg/mL of commercially available (**a**) *E. coli* O26:B6, (**b**) *E. coli* O55:B5, (**c**) *E. coli* O111:B4 and (**d**) *E. coli* O127:B8 LPS samples. Each data point represented results generated from 1 LPS sample, 1 lysates sample, 3 biological repeats (host cells used) and 2 technical repeats. Data were represented as mean ± SEM. (**e**) Schematic diagrams showing the predicted chemical formulas of *S. flexneri* 2a 2457O, *S. flexneri* 5a M90T, *E. coli* O3 and *E. coli* O111:B4 LPS, which were shown to give at least one log reduction in the transduction efficiency of P1(S’) when applied at concentrations lower or equal to 500 μg/mL. The LPS structures were determined in previous studies (Kenne et al., 1978; Stenutz et al., 2006; Ren et al., 2008; Huang et al., 2012; Perepelov et al., 2012). Differences between the outer core and O-antigens were highlighted in blue for the LPS of *S. flexneri* 5a M90T and *S. flexneri* 2457O, green for the LPS of *E. coli* O3 and yellow for the LPS of *E. coli* O111:B4. For simplification purpose, the orientation of glycosidic bond linking the O-antigen to the outer oligosaccharide core of LPS, as well as possible modifications to inner and lipid A core of LPS were not shown. The O-antigens are expected to be linked to the terminal α-D-glucose of both R2 (Heinrichs et al., 1998) and R3 (Kaniuk et al., 2004) type LPS outer cores via a (O-antigen)-1,4-(α-D-glucose of outer core) glycosidic bond. *p*, phosphorylation; *Ac*, acetyl group; GlcNac, *N*-acetylglucosamine; Rha, rhamnose; Glc, glucose, RhaNac; *N*-acetylrhamnosamine; Gal, galactose; Col, colitose.

Taken together, a low concentration of *E. coli* O111:B4 LPS blocked (S’) transduction completely, which indicated a high P1(S’) adsorption efficiency on *E. coli* O111:B4 LPS, and that the *E. coli* O111:B4 strain might be a host for P1(S’).

### P1(S’) lysates produced a higher Cas9-mediated antimicrobial effect on *S. flexneri* 5a M90T and *S. flexneri* 2a 2457O

P1(S’) infection of *S. flexneri* 5a M90T and *S. flexneri* 2457O yielded a higher number of phagemid transductant, indicating that P1(S’) could deliver the phagemid DNA into *S. flexneri* at a higher efficiency when compared to P1(S) (**Figure 1c**). We therefore hypothesised that an increase in the transduction efficiency of a *cas9* phagemid into *S. flexneri* would produce a higher antimicrobial effect on the bacteria. To verify this hypothesis, the *cas9* system was transduced into *S. flexneri* 5a M90T and *S. flexneri* 2a 2457O with either P1(S) or P1(S’), and the differences in the Cas9-mediated antimicrobial effect on both *S. flexneri* strains were assessed. A protospacer sequence targeting the chromosomal *shiA* gene sequence of *S. flexneri* was cloned into the *cas9* phagemid (phagemid termed *cas9-shiA* hereafter) (**Figure 6a**). *cas9-shiA* phagemid as well as the non-chromosomal-targeting, *cas9*-NT phagemid, were used for preparation of P1(S) and P1(S’) phagemid lysates. To verify if all the lysates yielded a similar titre of phagemid transducing particles, phagemid lysates were treated with DNase I and quantified for P1-packaged phagemid DNA via qPCR. All the DNase I-treated lysates yielded a similar concentration of P1-packaged phagemid DNA, suggesting that differences, if any, in the Cas9 antimicrobial effect on *S. flexneri* would be attributed to the relative transduction efficiency of phagemid DNA (**Supplementary Figures 2a, 2b**). *S. flexneri* 5a M90T and *S. flexneri* 2a 2457O were then treated with varying volumes of the *cas9-shiA* or *cas9*-NT phagemid lysates and the number of CFU recovered on non-selective agar was enumerated. The Cas9-antimicrobial effect was represented by a reduction in the number of CFU recovered after *cas9-shiA* phagemid lysates treatment, when compared to the use of *cas9*-NT phagemid lysates.

**Figure 6:**
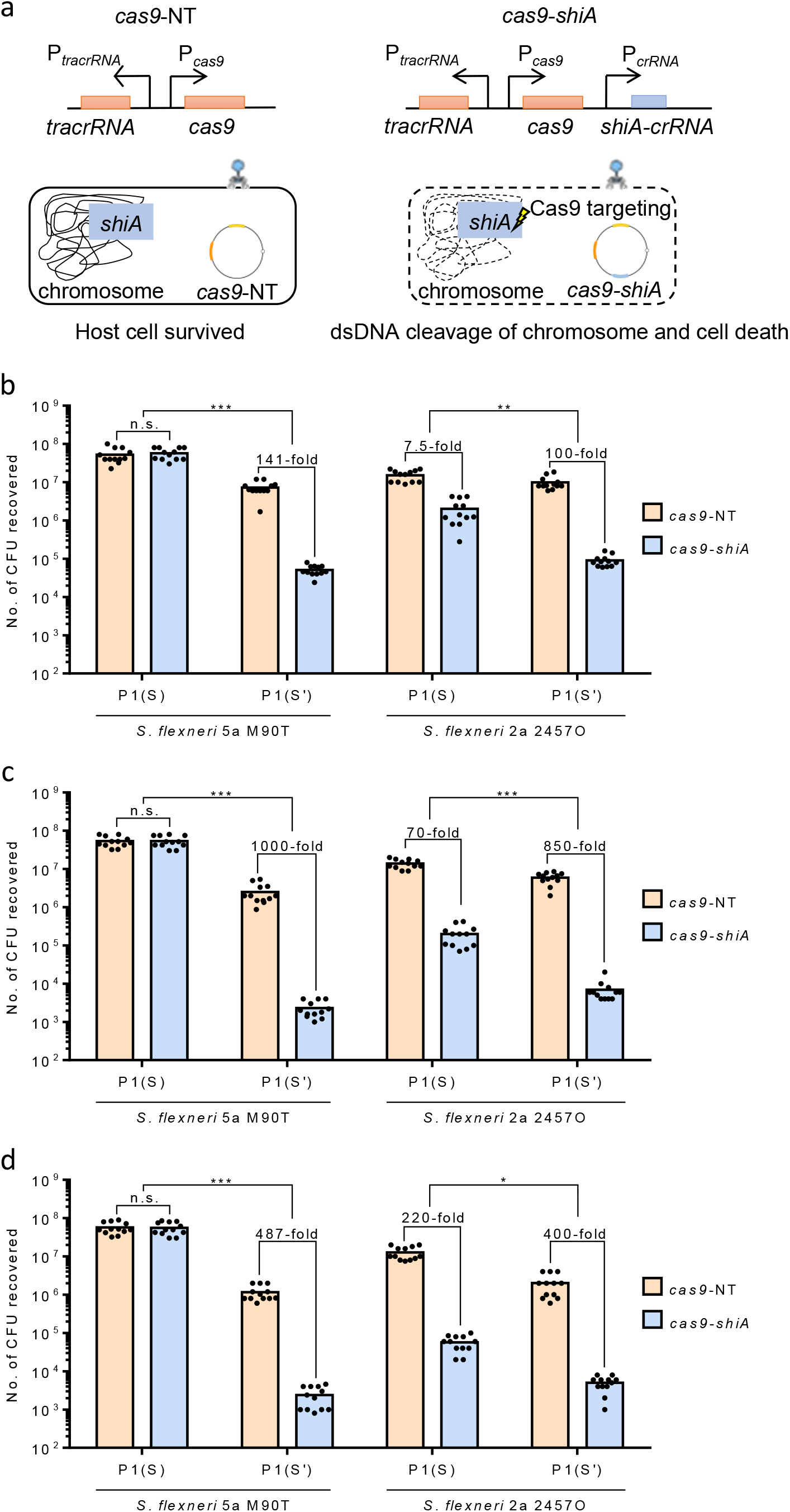
A higher antimicrobial effect of P1(S’) lysates on *S. flexneri* 5a M90T and *S. flexneri* 2a 2457O. **a)** Schematic diagram showing the *cas9* genetic construct of P1 J72114 phagemid, with protospacer sequence targeting chromosomal gene, *shiA* of *S. flexneri* (in light blue). Following transduction and circularisation of the phagemid, the presence of a CRISPR guide RNA (crRNA) with spacer sequence complementary to the chromosomal *shiA* gene of *S. flexneri* would cause dsDNA cleavage of the chromosome via Cas9 endonuclease, leading to cell death (shown as dashed lines). *cas9*-NT phagemid with a randomly-generated spacer sequence can be stably maintained in transduced cells (host cell survival indicated by filled lines). A reduction in the number of *S. flexneri* CFU recovered after *cas9-shiA* lysates treatment when compared to the use of *cas9*-NT lysates represented the Cas9-mediated antimicrobial effect. Quantification of the number of *S. flexneri* 5a M90T and *S. flexneri* 2a 2457O CFU recovered after treatments with **b)** 4 μL, **c)** 8 μL and **d)** 12 μL of P1(S) or P1(S’) non-targeting *cas9*-NT (light brown bars) and chromosomal-targeting *cas9-shiA* (light blue bars) lysates. Approximately 10^7^ CFU was used for each experiment. The differences in *S. flexneri* CFU recovered, between treatments with *cas9*-NT and *cas9*-*shiA* phagemid lysates were shown. n.s. represents no significant difference in the number of *S. flexneri* 5a M90T CFU recovered between P1(S) *cas9*-NT and P1(S) *cas9-shiA* lysates treatments. p < 0.0005 are shown as ***, for comparisons of the antimicrobial effects between P1(S) and P1(S’) *cas9* lysates on *S. flexneri* 5a M90T at all three volumes, as well as that on *S. flexneri* 2a 2457O at 8 μL of lysates. p < 0.005 is shown as **, for comparisons of the antimicrobial effects between 4 μL of P1(S) and 4 μL of P1(S’) *cas9* lysates on *S. flexneri* 2a 2457O. p < 0.05 is shown as *, for comparisons of the antimicrobial effects between 12 μL of P1(S) and 12 μL of P1(S’) *cas9* lysates on *S. flexneri* 2a 2457O.

4 μL to 12 μL of P1(S’) *cas9*-*shiA* lysates reduced the number of *S. flexneri* 5a M90T CFU by 141-fold to 1000-fold, when compared to the number of CFU recovered after treatment with P1(S’) *cas9*-NT lysates (**Figures 6b, 6c, 6d**). There were no significant differences in the number of *S. flexneri* 5a M90T CFU recovered between P1(S) *cas9-shiA* and P1(S) *cas9*-NT lysates treatment (p > 0.05), which indicated a significantly lower Cas9-mediated antimicrobial effect of P1(S) lysates on *S. flexneri* 5a M90T when compared to that of P1(S’) lysates. P1(S’) *cas9-shiA* lysates reduced the number of *S. flexneri* 2a 2457O CFU by 100-fold to 850-fold when compared to the number of CFU recovered after treatment with P1(S’) *cas9*-NT lysates (**Figures 6b, 6c, 6d**). P1(S) *cas9-shiA* treatment caused 7.5-fold to 220-fold reduction in the number of *S. flexneri* 2a 2457O CFU when compared to the number of CFU recovered P1(S) *cas9*-NT treatment, which indicated a lower Cas9-mediated antimicrobial effect of P1(S) lysates on *S. flexneri* 2a 2457O when compared to P1(S’) lysates.

Taken together, P1(S’) *cas9* phagemid lysates produced a greater antimicrobial effect on both *S. flexneri* 5a M90T and *S. flexneri* 2a 2457O, indicating that P1(S’) improved the transduction efficiency of *cas9* phagemids into these bacteria when compared to P1(S).

## DISCUSSION

P1 bacteriophage has an invertible C segment, encoding two sets of tail fibres namely the S and S’ (Iida et al., 1982; Hiestand-Nauer and Iida, 1983; Iida, 1984). Despite the wide usage of P1 for generalised transduction, the host range of P1(S’) was never determined. Here, we showed evidence of P1(S’) transduction on *S. flexneri* 2a 2457O, *S. flexneri* 5a M90T and *E. coli* O3, as well as characterised several interesting properties of this process, such as the requirement for O-antigen and a less stringent need of Ca^2+^ for transduction. Data presented in this study provided novel insights into P1 tropism-switching and showed promise as a tool for improving the transduction efficiency of a *cas9* antimicrobial phagemid into *S. flexneri*.

### The effect(s) of O-antigen on P1 transduction

The presence of O-antigen was reported to reduce the transduction efficiency of bacteriophage P1. Previous studies have shown that restoring the O-antigen expression of *E. coli* K12 MG1655 inhibited P1 phage transduction (Browning et al., 2013), while *gal* mutants of *E. coli* O157:H7 lacking O-antigen were permissive for P1 infection (Ho and Waldor, 2007). The inhibitory effect of O-antigen against P1 transduction contradicts with an earlier study by Godard et al., (1971), which showed that P1kc *vir* infected smooth strains of *S. flexneri* F6S. Notably, P1 lysates used in all these studies were prepared lytically on *E. coli* K12, which would have given a majority of P1 progeny with the S tail fibre, since only P1(S) phage can infect *E. coli* K12 and that inversion of the C segment is infrequent. Our results indicated that both the P1(S’) and P1(S) can transduce phagemid DNA into wildtype *S. flexneri* 5a M90T and *S. flexneri* 2a 2457O with intact O-antigen, albeit with varying efficiencies. Therefore, the inhibitory effects of O-antigen against P1 transduction might be dependent on the bacterial strains used, discrepancies between transduction of phagemid DNA versus P1 genomic DNA, as well as the tail fibre of P1 phage, in which the host range(s) of P1 alternative S’ tail fibre was not determined from previous studies (Iida et al., 1982; Iida, 1984).

### The host range of P1(S’) and the requirement of O-antigen for its transduction

Our results indicated that a lack of O-antigen inhibited P1(S’) transduction of *S. flexneri* 2a 2457O, *S. flexneri* 5a M90T and *E. coli* O3. Indeed, rough LPS extracted from the O-antigen deficient mutant strains lacked P1(S’) neutralising effect, highlighting the possible role of these O-antigens in mediating P1(S’) adsorption. O-antigen chains serve as receptors for the tailspike proteins of multiple bacteriophages, including bacteriophage P22 that infects *Samonella spp*. (Steinbacher et al., 1996) and bacteriophage Sf6 which infects *S. flexneri* serotypes X and Y (Freiberg et al., 2003). Most of these tail spike proteins promote the degradation and/or the modification of O-antigen chains thus orientating the bacteriophage to a secondary receptor for irreversible binding. Such mechanism, however, is expected to be different to that of the P1(S’) tail fibre, due to a lack of significant amino acid sequence homology to any known tail spike protein targeting *S. flexneri* O-antigen, nor to any known enzymatic domain involved in degrading glycosidic bond.

Comparative analysis of P1 S’ tail fibre amino acid sequence (NCBI, NC_005856.1, 32225..33782) indicated a high sequence similarity (≈ 99 %) to the protein product of multiple phage tail fibre genes that were annotated in *E. coli, S. enterica* and *E. albertii* genomes (data not shown). One such homologue of P1 *S’* gene is the *S*’ tail fibre gene of Mu bacteriophage (NCBI, NC_000929.1, 34067..35053). The Mu *S*’ gene shows a high sequence similarity (≈ 90 %) to the 3’-end (639^th^ to 2919^th^ DNA sequence) of P1 S’ gene, which encodes the predicted host-range-determining-region (HRDR) of P1 S’ tail fibre (NCBI, NC_005856.1, 32225..33782) (data not shown). The Mu S’ tail fibre, however, was shown to recognise a glucose-β 1,6-glucose structure of *Erwinia chrysanthemi* (B37) LPS (Sandulache et al., 1985), and this structure is not found on the LPS of *S. flexneri* 2a 2457O, *S. flexneri* 5a M90T, *E. coli* O3 and *E. coli* O111:B4. Hence, P1 S’ might have a different receptor compared to Mu S’, which is consistent with the fact that neither Mu(S) nor Mu(S’) formed plaques on 12 different serotypes of *S. flexneri* (Jakhetia and Verma, 2015). Interestingly, Jakhetia and Verma (2015) reported a Mu-like phage, SfMu which formed plaques on *S. flexneri* serotypes 3b and Y. The differences between Mu and SfMu host range is attributed to 18 amino acid changes in the tail fibre of SfMu when compared to Mu S’ tail fibre. Comparisons between the C-terminus of P1 S’ tail fibre, the tail fibre of SfMu (NCBI, KP010268, ORF49) and the Mu S’ tail fibre indicated that P1 S’ tail fibre contains additional amino acid changes at multiple sites, which might contribute towards differences in the host range of these phages (data not shown). A comparative analysis of the amino acid sequence between P1 S’ tail fibre and its homologs could determine how changes in specific amino acid residues lead to variations in the phages host range.

The LPS of *S. flexneri* 2a 2457O, *S. flexneri* 5a M90T, *E. coli* O3 and *E. coli* O111:B4 with O-antigen (termed O-PS hereafter) produced a high neutralising activity against P1(S’) transduction, which indicated common feature(s) shared among these O-PS might be the receptor for P1(S’). The predicted similarity between *S. flexneri, E. coli* O3 and *E. coli* O111 O-PS is a β-D-Glc*p*Nac-1,4-α-D-Glc residue which links the O-antigens to the terminal glucose of LPS outer core oligosaccharide (**Figure 5e**). This residue, however, is also present in the LPS of *E. coli* O26:B6 (Huang et al., 2012; Liu et al., 2020) which yielded a weaker P1(S’) neutralising effect compared to the LPS of wildtype *E. coli* O3 and *S. flexneri*. Therefore, it remained inconclusive if a β - D-Glc*p*Nac-1,4-α-D-Glc residue serves as the receptor for P1(S’) since LPS can adopt different conformations based on their respective side chains, which could have reduced the accessibility of this residue on the LPS of *E. coli* O26:B6 (Rodriguez-Loureiro et al., 2018, Gao et al., 2020). Furthermore, we could not dismiss the role of other cell surface antigens in mediating P1(S’) transduction and/or adsorption. Genomic analysis indicated the presence of a K2ab polysaccharide capsule operon and a H2 flagellar operon on the strain of *E. coli* O3 used for this study. Hence, the effects, if any, of these surface antigens on P1(S’) adsorption and/or transduction remain to be elucidated. Quantification of P1(S’) transduction efficiency on various gram-negative *Enterobactericae*, coupled with structural studies of the P1(S’) tail fibre as well as the use of synthetic glycans mimicking different regions of LPS oligosaccharides, will help in identifying its receptor and the mechanisms behind P1(S’) adsorption.

### P1(S) host range

Earlier studies suggested that P1(S) tail fibre recognises the terminal glucose molecule on the outer LPS core without discriminating the orientation of a glycosidic bond (Franklin, 1969; Ornellas and Stocker, 1974; Sandulache et al., 1984). Indeed, glucose molecule is present in the LPS of *E. coli*and *S. flexneri*strains tested in our study, which is consistent with the fact that P1(S) infected most of these bacteria. A previous study of *S. flexneri* 2a O-antigen indicated that loosing specific acetylation modification(s) affected the sensitivity of *S. flexneri* 2a 2457T towards bacteriophage Sf6c infection (Teh et al., 2020). Similarly, our results indicated that P1(S) transduction was significantly affected by a Δ*gtr* and a Δ*rfc* mutation in *S. flexneri* 5a M90T, as well as a Δ*oacB* mutation in *S. flexneri* 2a 2457O. While P1(S) failed to transduce phagemid DNA into wildtype *E. coli* O3, its isogenic Δ*waaL* and Δ*rfe* mutants were permissive for P1(S) transduction, and the rough LPS of its Δ*waaL* mutant produced a high inhibitory effect against P1(S). Taken together, accessibility of the P1(S) receptor might be affected by mutations in the genes of O-antigen assembly, biosynthesis and its modification. Further structural studies of the P1(S) tail fibre might provide insights into the molecular mechanisms behind P1(S) adsorption to its receptor.

### The role of free Ca^2+^ ion in P1(S) and P1(S’) transduction

Although historic protocol stated that Ca^2+^ is required for P1 transduction (Lennox, 1955), the role of Ca^2+^ in P1(S’) transduction was never determined. Our results indicated a less stringent requirement of free Ca^2+^ for P1(S’) transduction compared to P1(S). Although the exact role for Ca^2+^ in P1 transduction is not known, previous studies suggested that Ca^2+^ might mediate phage adsorption and/or the insertion of phage DNA into its bacterial host (Garen and Puck, 1955; Harada et al., 2013). Since both the P1(S) and P1(S’) would have the same central tail spike for the injection of DNA, discrepancies in the requirement of Ca^2+^ for transduction might be attributed to different receptor(s) between P1(S) and P1(S’) tail fibres. We could not rule out the role of other divalent cations in mediating P1(S) and P1(S’) transduction, especially since Mg^2+^ was supplemented in the preparation of phage lysates. Further studies involving specific ion chelators, the use of minimal media with defined ionic composition and structural studies of both tail fibres could provide insights into the role of Ca^2+^ in P1(S) and P1(S’) transduction.

### The potential use of P1(S’) for delivering an antimicrobial *cas9* system into pathogenic strains of *Enterobacteriaceae*

Besides *S. flexneri* and *E. coli* O3, the high inhibitory effect of *E. coli* O111:B4 LPS against P1(S’) transduction indicated that this *E. coli* strain might be a host for P1(S’). *E. coli* O111:B4, however, was shown to be resistant to P1 phage infection, due to the presence of a P1 or a P1-like resident phage (Coleman et al., 1977). Subsequently, cells cured from the lysogenic phage were susceptible to P1 infection using lysates prepared from a thermoinducible P1 lysogen. Since lysates prepared from an induced lysogen would yield both P1(S) and P1(S’) phages, the factor(s) required for P1(S’) adsorption might be present in *E. coli* O111:B4, which is consistent with the high P1(S’) neutralising effect of *E. coli* O111:B4 LPS. Serotyping analysis showed that *E. coli* O111 is associated with strains of enterotoxigenic *E. coli* (ETEC) (Campos et al., 1994) while *E. coli* O3 is associated with clinical isolates of enteroaggregative (EAEC) *E. coli* (Jenkins et al., 2006; Kai et al., 2010; de Lira et al., 2021). Hence, P1(S’) could be a valuable vector for the delivery of the *cas9* antimicrobial phagemid into clinical isolates of *E. coli* O111 and *E. coli* O3. Furthermore, P1(S’) transduction could also be expanded to target other serotypes of *S. flexneri*, since most serotypes share a common O-antigen backbone, and that P1(S’) transduction might not be significantly affected by changes in the O-antigen modifications of *S. flexneri*. The application of P1(S’) to target other serotypes of *S. flexneri* is relevant, as the development of a broadly protective vaccine against *S. flexneri* is hampered by its serotypic and genomic variations (Mani et al., 2016; Bengtsson et al., 2022). Notably, our preliminary transduction assay indicated that P1(S’) infection of *S. flexneri* serotype 2b ATCC® 12022 yielded a higher number of phagemid transductant when compared to that recovered after P1(S) infection (data not shown) thus highlighting the potential of P1(S’) for delivering the DNA sequence-specific Cas9 antimicrobial system into different *S. flexneri* serotypes.

## MATERIALS AND METHODS

### Bacterial strains, plasmids and media used in this study

All the bacterial strains used for this study were listed in **Supplementary Table 1** while plasmids used were listed in **Supplementary Table 2**. All PCR reactions for molecular cloning was carried out using Q5® High-Fidelity DNA Polymerase (NEB, M0491) while colony PCR was carried out using Taq Polymerase (NEB, M0270L). All plasmids were assembled using Gibson Assembly (NEB, E2621). Bacteria of this study were routinely cultured in Luria–Bertani (LB) broth and agar. Δ*rfc* mutants of *S. flexneri* exhibited very slow growth after lambda Red recombineering. Hence mutants were cultured on Tryptic Soy Broth (TSB) before subsequent culturing in LB media. SM buffer (50 mM Tris-HCl, 8 mM magnesium sulfate, 100 mM sodium chloride, pH 7.5), was routinely used as the buffer for manipulating phage lysates. DNA sequences used in this study were listed in **Supplementary Tables 4**.

### Genomic sequencing of *E. coli* O3 and agglutination assay

Preliminary results suggested that P1(S’) can transduce phagemid DNA into an unknown *E. coli* strain at a high efficiency. Since the identity of this bacterial strain was not determined, genomic sequencing was performed by MicrobesNG (Birmingham, United Kingdom, https://microbesng.com/), with llumina next-generation sequencing data at a coverage of 30X. Cell mass was provided to MicrobesNG. Genomic sequencing data is available at NCBI, under the accession number JALKIH000000000, Bioproject: PRJNA825326, Biosample: SAMN27512004. Artemis (Oxford, United Kingdom) was used to browse the assembly and annotations (Carver et al., 2012). The O-antigen gene cluster of *E. coli* is commonly located between the *galF* and *gnd* genes sequences (Stenutz et al., 2006), which was annotated in the assembly data. BLASTn (Nucleotide BLAST, NIH) was used to identify homologous DNA sequence, and the results suggested a 99 % (14043/14049) identity match to the *E. coli* O3 O-antigen gene cluster identified in Ren et al., study (2008). DNA sequence of the O3 O-antigen gene cluster was deposited onto GenBank, with the accession number ON210125. The expression of *E. coli* O3 O-antigen was confirmed by agglutination assay using anti-*E. coli* O3 antiserum (Oxford Biosystems, 84999). Overnight culture of *E. coli* O3 was boiled for 1 h on heat block at 100 °C. The boiled culture was left to stand at room temperature for 1 h to allow cell debris to sediment. Equal volume of antiserum was added to the boiled cultures in glass tubes, and the reactions were incubated overnight at 52 °C in a water bath. Glass tubes were shaken vigorously before reading the reaction. *E. coli* O3 culture gave a granular texture indicating positive reaction while a homogenous milky turbidity was observed for *S. flexneri* 2a 2457O, *S. flexneri* 5a M90T and *E. coli* E24377A cultures, indicating negative reactions.

### Transformation of *E. coli* and *S. flexneri*

Transformation of *E. coli* and *S. flexneri* was carried out using standard electroporation protocol. Overnight culture of cells was sub-cultured by 1/100 in fresh LB media at 37 °C with shaking until it reached an OD_600_ of approximately 0.35. Cells were centrifuged at 3000 X g for 15 mins at 4 °C. Cell pellet was washed with ice-cold 10 % glycerol (in distilled water), and the washing step was repeated for another 2 times. The final cell pellet was concentrated by 200-fold in distilled water with 10 % glycerol (e.g., 250 μL of 10 % glycerol for cell pelleted from a 50 mL culture). 1 μL of DNA was electroporated into 40 μL of cells using Gene Pulser/MicroPulser Electroporation Cuvettes, 0.1 cm gap (Biorad, 1652083), and immediately transferred into 1 mL of ice-cold SOC. Cells were incubated at 30 °C for temperature sensitive plasmids, pKD46 and pCP20, or 37 °C for other DNA substrate, with shaking for 1 h. Cells were platted onto LB agar plates with the appropriate antibiotics, followed by overnight incubation at 30 °C or 37 °C.

### P1 phagemid lysate preparation

Preparation of phagemid lysates was carried out based on a protocol from Kittleson et al. (2012) study, with slight modifications. Overnight cultures of *E. coli* P1 lysogenic EMG16 cells transformed with phagemid and/or with plasmid, were sub-cultured in fresh PLM media (LB supplemented with 5 mM of Ca^2+^ and 10 mM Mg^2+^) at 37 °C, with shaking until it reached an OD_600_ between 0.15 to 0.2, which took approximately 2 hours. L-arabinose was added to the culture, giving a final concentration of 0.02 % or approximately 1.3 mM. Induction was carried out at 37 °C, with shaking, until clear culture was obtained, which took approximately 2 h to 3 h. Lysates were transferred into 1.5 mL polypropene tubes and chloroform was then added to the culture, giving a final concentration of 2.5 % (e.g., 25 μL for 1 mL of lysate). Cultures were incubated at 37 °C, with shaking for at least 15 mins. Samples were centrifuged at maximum speed for 3 mins. The supernatant was carefully collected and filtered using a 0.22 μm Syringe filter (Millipore). Lysates can be stored at 4 °C.

For Polythene Glycol 6000 (PEG 6000) precipitation, sodium chloride and chloroform were added to lysates, giving a final concentration of 0.3 M and 2.5 % respectively. Cultures were incubated at 37 °C with shaking for at least 15 mins. Lysates were centrifuged at 5000 rpm for 30 mins and the supernatant was carefully collected. PEG 6000 was added to a final concentration of 4 % (e.g., 4 g in 100 mL of lysate). Lysates were shaken until all the PEG 6000 dissolved, and incubated overnight at 4 °C. The PEG precipitate was spun down at 5000 rpm for 45 mins, supernatant was then removed, and SM buffer was added to give 100-fold concentration of phage precipitate (e.g., 500 μL SM buffer added to the pellet obtained from a 50 mL lysates). Pellet was dissolved by leaving it in SM buffer for 2 h, followed by gentle vortexing and resuspension using Gilson Pipette. The lysates were then passed through a 0.22 μm Syringe filter (Millipore) and stored at 4 °C.

### Quantification of phagemid transductant

Quantification of phagemid transductant was carried out based on a protocol derived from Kittleson et al. (2012) study with slight modifications. Overnight cultures of the host cells were sub-cultured in fresh LB media at 37 °C, with shaking until it reached an OD_600_ of 0.5. Cells were concentrated by 10-fold, and used at 100 μL for transduction, which would give a range of 1 X 10^8^ to 5 X 10^8^ cells per reaction. 10 μL of crude P1 phage lysates, or 10 μL of 10-fold diluted PEG 6000 purified lysate, was diluted in 90 μL of SM buffer supplemented with 5 mM Ca^2+^. 100 μL of host cells were then added to 100 μL of phage lysates, incubated at 37 °C for 30 mins with shaking to maximise phage adsorption. 800 μL of SOC with 10 mM citrate was added to the mixture, and cells were incubated for 1 h at 37 °C with shaking for the expression of chloramphenicol-resistant gene. Serial dilution of the final mixture was made, and 5 μL of each dilution were platted onto LB agar supplemented with 25 μg/mL of chloramphenicol. Plates were incubated at 37 °C overnight and the number of chloramphenicol resistant CFU were enumerated after 14 h to 16 h of incubation.

### Genetic modification of *E. coli* P1 lysogen and *S. flexneri*

All genetic modifications of *E. coli* P1 lysogen, *S. flexneri* 2a 2457O and *S. flexneri* 5a M90T were carried out via lambda Red recombineering technique as described in a previous study by Datsenko and Wanner (2000). The gene knockout template was designed to have homology arms of 60 bps complementary to both upstream and downstream DNA sequence of the target site (refer to **Supplementary Table 3** for primer sequence). The marker gene for selection of mutation was a constitutively expressed kanamycin resistance gene cassette, *npt*, flanked by Flippase recognition sites (FRT). The dsDNA DNA template was generated via PCR, and agarose gel purified. The target bacteria were transformed with pKD46 plasmid. Stationary phase culture of the transformed cells was outgrown in fresh LB at a dilution factor of 100, at 30 °C with shaking, until it reached an OD_600_ of 0.35. 0.1 % or 0.065 M of L-arabinose was added to the culture to induce the expression of Lambda Red genes (*exo*, *bet, gam)*, and cultured for a maximum of 30 mins at 30 °C, with shaking. Cells were chilled on ice for 40 mins and made electrocompetent, using standard protocols for preparing electrocompetent cells. 100 ng of dsDNA substrate was electroporated into the cells, followed by outgrowth in SOC at 30 °C with shaking for 2 h. Cells were plated onto LB agar supplemented with 50 μg/mL of kanamycin, incubated at 37 °C for at least 16 h. Colony PC R was carried out on colonies recovered, using primer pairs that anneal to the junction of integration to verify correct insertion of the *npt* cassette. Positive clones were re-streaked onto new LB supplemented with 50 μg/mL of kanamycin, incubated at 37 °C for at least 16 h, and this re-streaking process was repeated for at least 3 generations. To remove the *npt* cassette, cells were transformed with pCP20, platted onto LB agar supplemented with 100 μg/mL of ampicillin and incubated at 30 °C for at least 16 h. Colonies were picked for PC, to verify the loss of *npt* cassette. PCR products of the correct sizes were gel-purified and sent for sequencing. For curing of the pCP20 plasmid, colonies having the correct PCR band size and DNA sequence were diluted in 5 mL of LB broth without ampicillin and incubated overnight at 43 °C with shaking. Serial dilution of the culture was made, and 25 μL of the dilution range of 10^3^ to 10^6^ was platted onto LB agar, followed by incubation at 30 °C for at least 16 h. Colonies were then platted onto LB agar with or without 50 μg/mL kanamycin or 100 μg/mL ampicillin. Agar plates with or without kanamycin were incubated at 37 °C, while agar plates with ampicillin were incubated at 30 °C, for at least 16 h. Colonies were selected for its sensitivity towards both kanamycin and ampicillin.

### Extraction and visualisation of LPS

LPS of *S. flexneri* and *E. coli* O3 were extracted using the hot aqueous-phenol method as described in Davis et al., (2012) studies with slight modifications. Briefly, bacterial suspension of approximately 10^8^ CFU/mL were centrifuged at 5000 rpm for 10 mins. Bacterial cell pellet was washed twice in PBS (0.15 M, pH= 7.2). For every 10^8^ CFU, cell pellet was gently resuspended in 200 μL of 1 X Laemmli Buffer (Biorad) supplemented with β-mercaptoethanol at a final concentration of 2.5 %. Cells were boiled in a water bath or heat block (at 100 °C) for 15 mins and cooled to room temperature. For every 200 μL of sample, 5 μL of 1 unit/μL DNase I (Sigma, AMP D1) and 5 μL of 10 mg/mL RNase A (Thermo, EN0531) were added and incubated at 37 °C for 1 h. For every 200 μL of sample, 5 μL of 20 mg/mL Proteinase K (Thermo, AM2546) was added, and incubated at 59 °C for 3 h. To extract the LPS, an equal volume of hot (65 °C to 70 °C) UltraPure™ Buffer-Saturated Phenol (Thermo, 15513039) was added to the sample, followed by vigorous mixing and incubation at 65 °C in a water bath or heat block for 20 mins. Samples were vortexed occasionally during the 20 mins incubation time. Samples were transferred to 1.5 mL polypropene tubes, cooled to room temperature. For every 400 mL of samples, 1 mL diethyl ether (Sigma, 91238) was added, and the mixture was vortexed for 5 s to 10 s. The samples were then centrifuged at 20600 X g for 10 mins at 4 °C. The bottom aqueous layer was carefully removed. Phenol extraction was repeated, by using the same heating and centrifugation steps mentioned previously on the aqueous layer. The samples were combined, and LPS was precipitated by adding 10 volumes of 90 % ethanol and 3 M sodium acetate to a final concentration of 0.5 M, and left overnight at −20 °C, as described by Rezania et al., (2011). The mixture was centrifuged at 4000 rpm for 15 mins at 4 °C, supernatant was removed, and the pellet is resuspended in distilled water to give a 10 X concentrated LPS solution (e.g., 0.5 mL of distilled water for pellet centrifuged from 5 mL of sample). To remove contaminating phenol, samples were extensively dialysed in distilled water at 4 °C, using Slide-A-Lyzer™ Dialysis Cassettes, 10K MWCO (Thermo, 66380). The final LPS samples can be stored at 4 °C for short term usage or at −20 °C for long term storage.

LPS samples were visualised using Pierce™ Silver Staining kit (Thermo, 24612), following manufacturer’s protocol. For concentrated LPS, samples were diluted by 10-fold in distilled water. Laemmli buffer was added to samples.10 μL of samples were loaded onto 12 % Mini-P ROTEAN® TGX™ Precast Protein Gels (Biorad). Silver staining was carried out at room temperature for 1 h and gels were developed for 2 to 3 mins, until bands were visible. Gel images were scanned using Canon CanoScan LiDE 120 and images were processed, by adjusting the brightness and contrast of images using ZEN 2 (blue edition) | V1.0 en. Images of 600 dpi resolution were used for this manuscript. Coomassie Blue staining of LPS samples were routinely carried out to determine protein contamination of samples, if any.

### Estimation of LPS concentration and LPS inhibition assay

LPS of *E. coli* K12 (Invivogen, tlrl-eklps), *E. coli* O26:B6 (Sigma, L8274), *E. coli* O55:B5 (Sigma, L2880), *E. coli* O111:B4 (Sigma, L2630) and *E. coli* O127:B8 (Sigma, L3129) were reconstituted in distilled water to give a final concentration of 5 mg/mL following manufacturers’ protocol. Samples were stored at 4 °C for short term usage or at −20 °C for long term storage. Silver staining of the LPS was carried out to verify presence of O-antigen repeating units (data not shown).

The concentration of LPS samples were estimated using a colorimetric assay for estimating carbohydrate concentration, developed by Masuko et al. (2005). A range of diluted LPS samples were made in distilled water to give a final volume of 30 μL. 150 μL of concentrated sulfuric acid (Sigma, 84727) was added to each sample, followed by vigorous mixing and incubation for 15 mins at 90 °C. 30 μL of 5 % phenol solution (in distilled water) was added to the mixture, followed by vigorous mixing, and incubation at room temperature with shaking for 5 mins. The absorption was then measured at a wavelength of A_490_. The concentrations of smooth LPS with O-antigen repeating units were estimated, using standard curves generated from *E. coli* O26:B6, *E. coli* O55:B5, *E. coli* O111:B4 and *E. coli* O127:B8 LPS. The concentrations of rough or semi rough LPS (without O-antigen or with only one O-antigen unit) were estimated using standard curves generated from *E. coli* K12 LPS.

For the LPS inhibition assay, a modified protocol based on Sandulache et al., (1984) study was used. Briefly, 0.5 μg, 5 μg, 50 μg and 500 μg of LPS were resuspended in distilled water, giving a final volume of 100 μL. PEG 6000-purified P1 phage lysates were diluted in SM buffer giving 1 X 10^3^ in 50 μL. Phage lysates and LPS samples were mixed with Ca^2+^ (to a final concentration of 5 mM), incubated at 37 °C for 1 h with shaking. Indicator cells, *E. coli* NCM3722 for P1(S) or *S. flexneri* 2a 2457O for P1 (S’) were prepared as described for **quantification of phagemid transductant**. 50 μL of indicator cells were added to the LPS and phage lysate mixture, followed by incubation at 37 °C for 15 mins with shaking. Cells were centrifuged for 4000 X g for 1 min at room temperature to remove unabsorbed phage. Cell pellet was then washed twice with 500 μL of SOC with 10 mM citrate supplement. The cell pellet was resuspended in a final volume of 1 mL SOC with 10 mM citrate supplement. Cells were incubated at 37 °C for 1 h with shaking. Cells were platted onto LB agar supplemented with 25 μg/mL of chloramphenicol, incubated at at 37 °C for at least 16 h before enumeration of chloramphenicol resistant CFU. The transduction efficiency was calculated by comparing the number of phagemid transductants to that recovered after treatment with non-LPS treated phagemid lysates.

### Quantification of the antimicrobial effect of P1 *cas9-shiA* phagemid

The CRISPR-Cas9 construct of this study was derived from pCas9 plasmid (Addgene plasmid #42876) and assembled onto P1 phagemid via Gibson assembly. The system provides constitutive expression of *cas9*, encoding the endoculease, as well as a *trans*-activating CRISPR RNA, tracRNA and a crRNA guide. The *crRNA* coding sequence contains two BsaI sites, allowing the ligation of a protospacer sequence complementary to chromosomal DNA sequence of the targeted bacterial species. Protospacer sequence complementary to the *shiA* gene sequence of *S. flexneri* was cloned into the *crRNA* coding sequence via BsaI digestion of the phagemid and the ligation of an annealed, oligonucleotide pair (*shiA*-targeting oligonucleotide pair, forward : 5’AAACGCATGACTTCTCCGGCTCTCG-3’; reverse : 5’-AAAACCGAGAGCCGGAGAAGTCATG-3’), as described in by Jiang et al. (2013). Phage lysates were prepared as described previously, using the Δ*pac E. coli* EMG16 P1 lysogen transformed with either the targeting *cas9-shiA* phagemid or non-targeting *cas9*-NT phagemid. Since a *shiA*-targeting phagemid is inherently unstable in *S. flexneri*, the titre of phagemid transducing particle was first assessed using qPCR (**Supplementary Figures 2a, 2b**) as well as estimated on *E. coli* NCM3722 for P1(S) lysates and *E. coli* O3 for P1(S’) lysates (**Supplementary Figures 8a, 8b**). The indicator cells were prepared as described previously for the **quantification of phagemid transductants**. Overnight cultures of *S. flexneri* were sub-cultured in fresh PLM media (LB supplemented with 5 mM of Ca^2+^ and 10 mM Mg^2+^) at 37 °C with shaking. For *S. flexneri* 2a 2457O, cells were cultured until an OD_600_ of approximately 0.3, which was diluted to give an OD_600_ of approximately 0.1. *S. flexneri* 5a M90T exhibited a slower growth rate, hence cells were cultured for 2 h, giving an OD_600_ of approximately 0.12 and cultures were used directly for experiments. 100 μL of the cultures gave approximately 1 X 10^7^ cells per reaction. The *shiA*-targeting and non-targeting phagemids lysates were diluted in SM buffer supplemented with 5 mM Ca^2^ to a final volume of 100 μL. 100 μL of host cells were then added to 100 μL of phage lysates, incubated at 37 °C for 30 mins with shaking for phage adsorption. 800 μL of SOC was added to the mixture, and cells were incubated for 1 h at 37 °C with shaking. Serial dilution of the final mixture was made, and 5 μL of each dilution were platted onto LB agar. Plates were incubated at 37 °C overnight and the number of CFU were enumerated after 14 h to 16 h of incubation. Differences in the number of *S. flexneri* CFU recovered between the *cas9-shiA* and *cas9*-NT phagemid lysates treatment represented the Cas9-mediated antimicrobial effect

### Statistical analysis

Data of this study was generated from at least 2 phage lysate samples, 3 biological repeats (host cells tested) and 2 technical repeats, unless as stated otherwise. 1 lysate sample, 3 biological repeats (host cells tested) and 3 technical repeats were used for experiments investigating the role of Ca^2+^ in P1 transduction. Data of LPS inhibition assays were generated from 1 LPS sample, 1 phage lysate sample, 3 biological repeats (host cells tested) and 2 technical repeats. Statistical analysis of this study was carried out using Real Statistics Resource Pack on Microsoft excel (Microsoft, Redmond, WA, USA). *s* represented S.E.M while *n* represented the number of both biological and technical repeats. Welch’s ANOVA was used to determine the p-values, unless for comparisons involving 0 values, such as the total inhibition of transduction efficiency for LPS inhibition assay. In the latter case, a two-tailed, Student’s t-test (assuming unequal variance) was used. Graphpad Prism 6 was used to generate the graphs of this study. *p*-value of < 0.05 is considered statistically significant, for this study. The exact p-values of comparisons made are available in **Supplementary Materials 2**.

## SUPPLEMENTARY DATA

Supplementary Data are available at Cell Reports Online, including 6 Supplementary Figures and 3 Supplementary Tables.

## DATA AVAILABILITY

All relevant data are within the manuscript and supplementary information files. Data will be shared upon request to the corresponding author.

## ACKNOWLEDGEMENTS

This work was supported by the Bill and Melinda Gates Foundation under the Grand Challenges Explorations grant (OPP1139488), and UK Research and Innovation Future Leaders Fellowship [MR/S018875/1]. The authors wish to thank Professor Garry Blakely and Dr. Tahar Ait-Ali, for their kind guidance, advice and support for this study.

## AUTHOR CONTRIBUTIONS

B.W. conceived and supervised the study. B.W., Y.H., J.F. designed the experiments. Y.H. performed the experiments related to the P1 phagemid design, construction, production and characterisation in bacterial culture and P1 phage genome engineering. Y.H performed data analysis. All authors took part in the interpretation of results and preparation of materials for the manuscript. Y.H. and B.W. wrote the manuscript with input from all co-authors.

## COMPETING INTERESTS

The authors declare no competing interests.

## Notes

### Competing Interest Statement

The authors have declared no competing interest.

