## Supplementary Materials 1 for "The host range and the role of O-antigen in P1 transduction with its alternative S’ tail fibre"

**Table of Contents**

Supplementary Figures 1-8

Supplementary Tables 1-4

Supplementary Materials and Methods

Supplementary References

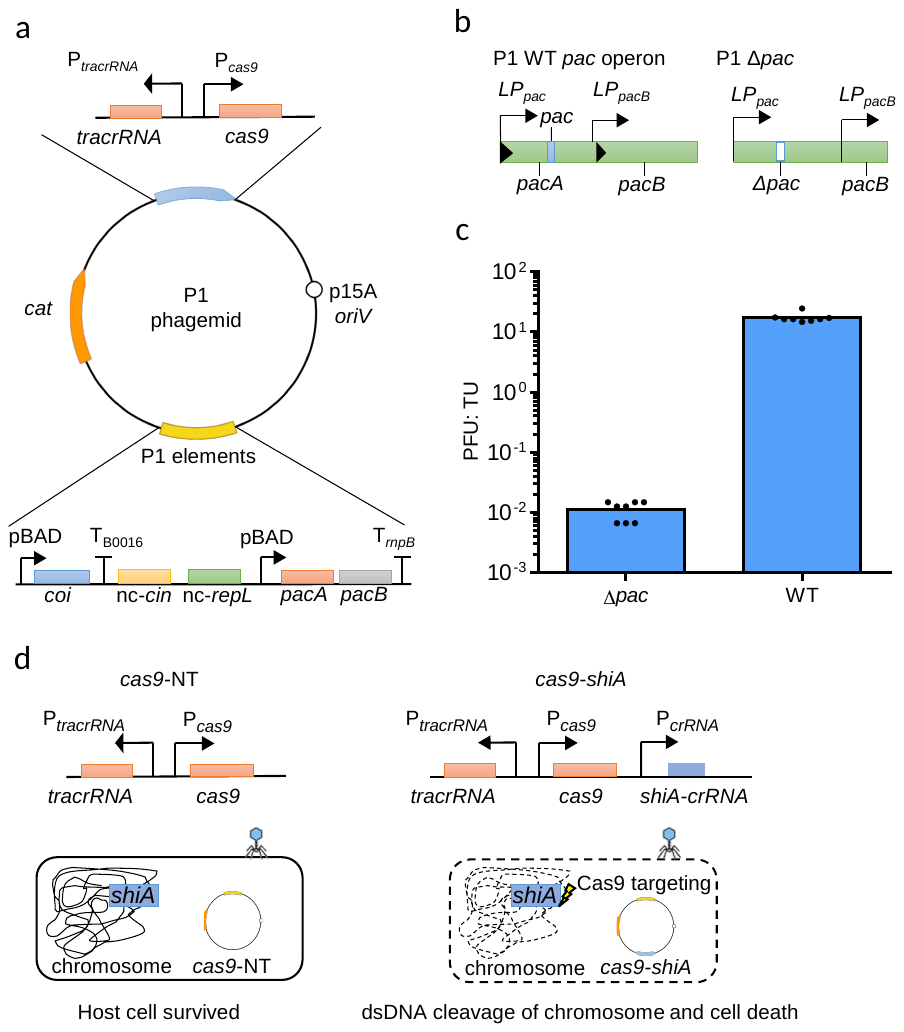

**Supplementary Figure 1: The use of a Δ*pac* *E. coli* P1 lysogen EMG16 to package an antimicrobial *cas9* phagemid.** (**a**) Schematic diagram showing the design of a P1 *cas9* phagemid, which was adapted from the phagemid (Addgene plasmid #40781) developed by Kittleson et al. (2012), with some modifications. The P1-based elements of our J72114 phagemid are *coi*, *pacA*, *pacB*, non-coding copy of *repL* and *cin* gene sequences. *coi*, whose gene product acts as a repressor antagonist of c1 repressor, hence acting as the switch that induces lytic stage replication of P1 bacteriophage (Heinzel et al., 1990). *coi* gene expression was controlled by the arabinose-inducible promote P_BAD_, which allows *trans*-activation of P1 lytic lifecycle and the packaging of phagemid into transducing particles (Kittleson et al., 2012). A non-coding copy of P1 *repL* gene, which contains the P1 lytic stage origin of replication, *ori_L_*, was included into the phagemid to allow replication of phagemid DNA during P1 lytic lifecycle (Westwater et al., 2002; Kittleson et al., 2012). A copy of both P1 *pacA* and *pacB* gene were included and their expression were controlled by the arabinose inducible P_BAD_ promoter. *pacA* and *pacB* gene product form the pacase enzyme, which is involved in processing and packaging of DNA into P1 procapsids (Sternberg and Coulby, 1988; Skorupski et al., 1992). The *pacA* coding sequence contains a *pac* site, which is recognised by the pacase enzyme, which allows packaging of phagemid DNA into transducing particles (Sternberg and Coulby, 1988; Skorupski et al., 1992). Finally, a non-coding copy of *cin* was included to increase phagemid transduction efficiency via an unknown mechanism (Kittleson et al., 2012). All our *cas9* phagemids contain a p15A origin of replication and a copy of *cat* gene, which confers chloramphenicol resistance after transduction. (**b**) Schematic diagram showing the wildtype *pac* operon of P1 genome. The 161 bps *pac* site internal to *pacA* gene sequence (in blue) was deleted (in white with white dashed lines) via lambda Red recombineering technique, and the mutant P1 lysogen is termed Δ*pac* *E. coli* EMG16. The deletion of *pac* was an attempt to reduce packaging of P1 genomic DNA, such that the purity of phagemid transducing particles could be improved. The mutation was complemented *in trans*, via the *pacA* and *pacB* genes of the P1 *cas9* phagemid, which would provide a functional pacase activity for the packaging of DNA into P1 procapsids. (**c**) Quantification of the ratio of P1 wildtype phage: phagemid transducing particles, for lysates prepared from wildtype (WT) and Δ*pac* *E. coli* EMG16 P1 lysogen. The tire of wildtype P1 progeny and phagemid transducing particles were calculated by enumerating the number of plaque forming units (PFU) and chloramphenicol resistant (Cm^R^) CFU recovered on *E. coli* K12 NCM3722 after lysates treatment, respectively. Data were generated from 4 lysate samples, 2 biological repeats (host cells tested), with each of the data point represented the average value of 2 technical repeats. Lysates prepared from the Δ*pac* P1 lysogen yielded a 1700-fold (*s* = 250, *n*= 16, p < 0.00005) lower ratio of PFU: TU when compared to lysates of wildtype P1 lysogen, which indicated that the deletion of *pac* could increase the purity of phagemid transducing particles. (**d**) Schematic diagram showing the *cas9* genetic construct of P1 J72114 phagemid, with a protospacer sequence targeting the chromosomal gene, *shiA* of *S. flexneri* (in blue). Following transduction and circularisation of the phagemid, the presence of a CRISPR guide RNA (crRNA) with spacer sequence complementary to chromosomal *shiA* gene of *S. flexneri* would cause dsDNA cleavage of the chromosome via Cas9 endonuclease, leading to cell death (shown as dashed lines). *cas9*-NT phagemid with a randomly-generated spacer sequence could be stably maintained in transduced cells (survival of host cell indicated by filled line). The DNA-sequence specific, Cas9 killing of bacteria was demonstrated previously by Bikard et al (2014), Citorik et al (2014), Gomaa et al (2014) and Cui and Bikard (2016).

| **Registry** | **Lysates** |
| --- | --- |
| 1 | P1(S) *cas9*-NT 1 |
| 2 | P1(S) *cas9*-NT 2 |
| 3 | P1(S') *cas9*-NT 1 |
| 4 | P1(S') *cas9*-NT 2 |
| 5 | P1(S) *cas9*-*shiA* 1 |
| 6 | P1(S) *cas9*-*shiA* 2 |
| 7 | P1(S') *cas9*-*shiA* 1 |
| 8 | P1(S') *cas9*-*shiA* 2 |

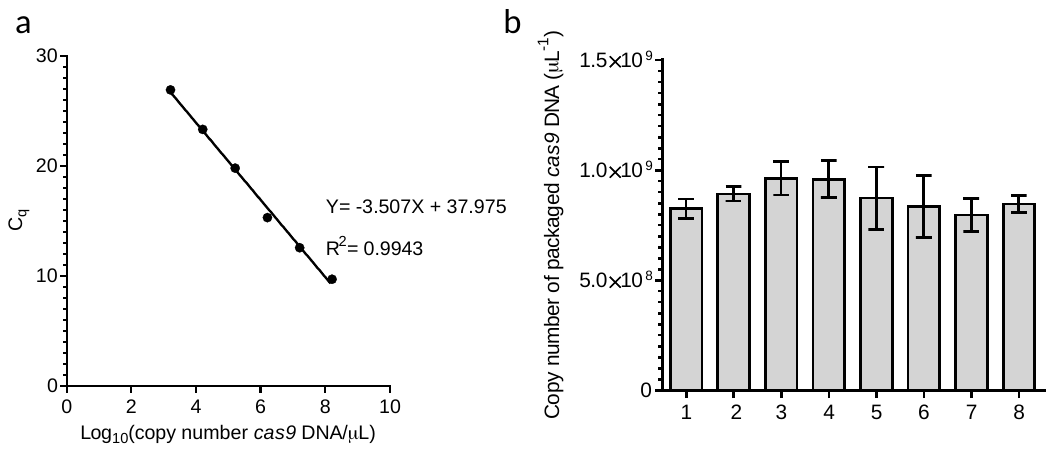

**Supplementary Figures 2: All DNase I-treated lysates yielded a similar concentration of P1-packaged phagemid DNA.** (**a**) The Standard curve of *cas9* DNA from SYBR GREEN qPCR. Log_10_(copy number/μL) was determined from serial dilutions of *cas9* DNA fragment from a concentration range of 1.28 X 10^3^ to 1.28 X 10^8^ copy number/μL. The quantification cycle (Cq) values were the SYBR GREEN qPCR results at the respective concentrations. Linear equation and correlation coefficient were shown in the graph. Efficiency of qPCR was 92.82 % or represented by an E_S_ of 0.928, as determined using the Thermofisher qPCR efficiency calculator. Standard deviation (SD) of Cq at all 6 concentrations ranged from 0.085 to 0.223, giving an average SD of 0.150 cycles. The DNA molecule deviation was calculated using the following equation reported from a study by Rutledge and Côté (2003): ± % DNA molecule = ((Es + 1)^SD^ – 1) x 100%

Based on the SD values determined earlier, the DNA molecule derivation ranged from ± 5.770 to ± 15.768%, with an average deviation of ± 10.318%. (**b**) Copy number of packaged *cas9* DNA sequence per μL phagemid lysates. 1 μL of DNase-treated phagemid lysates was used for qPCR. The names of lysates corresponding to each of the registry were listed in the table below the graphs. 3 technical repeats were carried out for each lysate. Data were represented as mean ± standard deviation (SD). Comparisons of the mean values between all lysates gave a p-value of 0.2647 using Welch’s ANOVA test, indicating no significant difference in the concentration of P1-packaged phagemid DNA between all lysates used in this study.

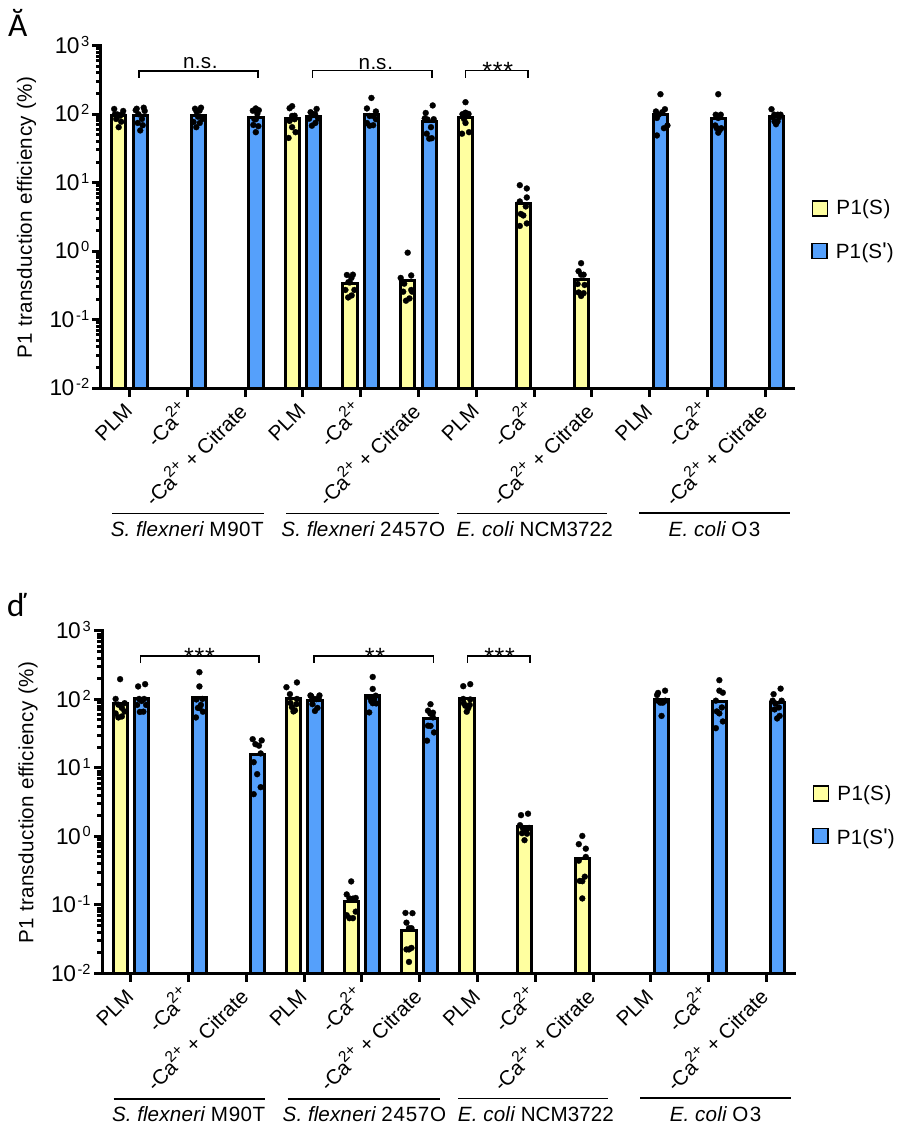

**Supplementary Figure 3: P1(S') transduction has a less stringent Ca^2+^ requirement compared to P1(S).** The transduction efficiency of P1(S) (yellow bars) and P1(S') (blue bars) when assessed on *S. flexneri* 5a M90T (*S. flexneri* M90T), *S. flexneri* 2a 2457O (*S. flexneri* 2457O), *E. coli* NCM3722 and *E. coli* O3 in PLM, without 5 mM Ca^2+^ supplement (- Ca^2+^), or with 10 mM citrate supplement (- Ca^2+^ + Citrate), at MOI (transducing units: host cell) of approximately (**a**) 0.1 and (**b**) 0.01. For these experiments, only PEG 6000-purified lysates were used. The relative transduction efficiency was calculated by comparing the number of phagemid transductant to that recovered after phage adsorption in PLM medium. Data were generated from 1 lysate sample, 3 biological repeats (host cells tested) and 3 technical repeats. The p-values were determined by Welch’s ANOVA test. n.s. represented no significant difference(s) for comparisons of P1(S') transduction efficiency in *S. flexneri* 5a M90T and *S. flexneri* 2a 2457O between PLM and with the addition of citrate, at a MOI of 0.1. p < 0.005 was shown as **, for comparison of P1(S') transduction efficiency in *S. flexneri* 2a 2457O between PLM and with the addition of citrate, at a MOI of 0.01. p < 0.0005 was shown as ***, for comparison of P1(S) transduction efficiency in *E. coli* K12 NCM3722 between PLM and without Ca^2+^ supplement at both MOIs of 0.1 and 0.01, as well as comparison of P1(S') transduction efficiency in *S. flexneri* 5a M90T between PLM and with the addition of citrate at MOI of 0.01.

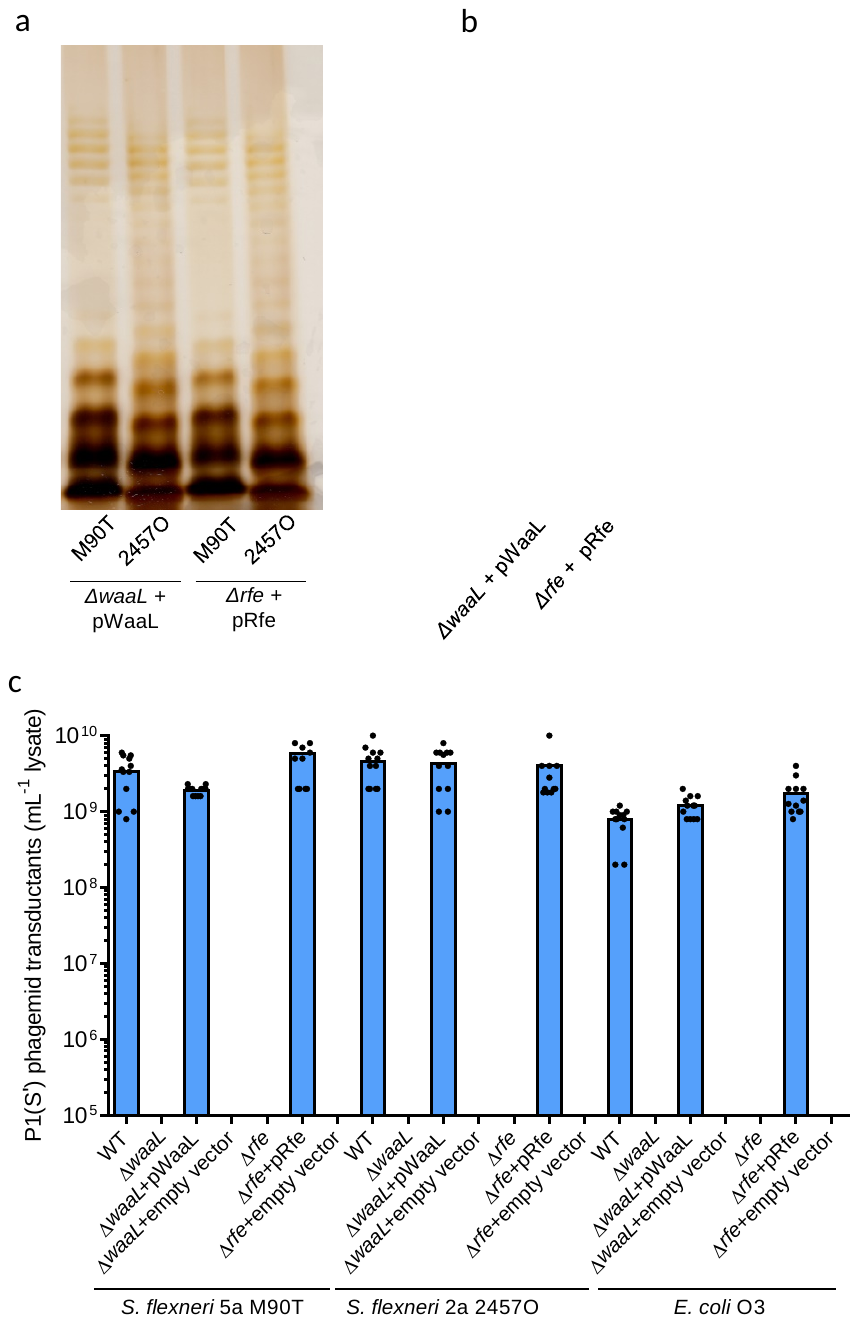

**Supplementary Figure 4: Complementation of the Δ*waaL* and Δ*rfe* mutations *in trans* restored P1(S') transduction of the mutations.** Analysis of LPS extracted from (**a**) Δ*waaL* and Δ*rfe* mutant strains of *S. flexneri* 5a M90T (M90T) and *S. flexneri* 2a 2457O (2457O) and (**b**) Δ*waaL* and Δ*rfe* mutant strains of *E. coli* O3, transformed with pWaaL (Δ*waaL* + pWaaL) and pRfe (Δ*rfe* + pRfe) respectively. Extracted LPS samples were analysed with SDS-PAGE and silver staining. All LPS samples gave a ladder like pattern of multiple bands, suggesting that the O-antigen repeats of LPS were restored in the Δ*waaL* and Δ*rfe* mutants. Δ*rfe* mutant strains of *S. flexneri* 5a M90T, *S. flexneri* 2a 2457O and *E. coli* O3 were transformed with the same pRfe plasmid since the DNA sequence of *rfe* gene was conserved between *S. flexneri* and *E. coli* O3. Δ*waaL* mutant strains of *S. flexneri* and *E. coli* O3 were transformed with pWaaL having its respective *waaL* gene sequences (pWaaL-*S. flexneri* and pWaaL-*E. coli O3* plasmids, respectively). (**c**) The number of P1(S') phagemid transductant recovered after lysates treatment of Δ*waaL* and Δ*rfe* strains of *S. flexneri* 5a M90T, *S. flexneri* 2a 2457O and *E. coli* O3 transformed with a pWaaL (+ pWaaL), a pRfe (+ pRfe) or an empty RK2 plasmid. Transformation of the Δ*waaL* and Δ*rfe* mutants with a pWaaL and a pRfe plasmid respectively restored the recovery of phagemid transductants after P1(S') infection.

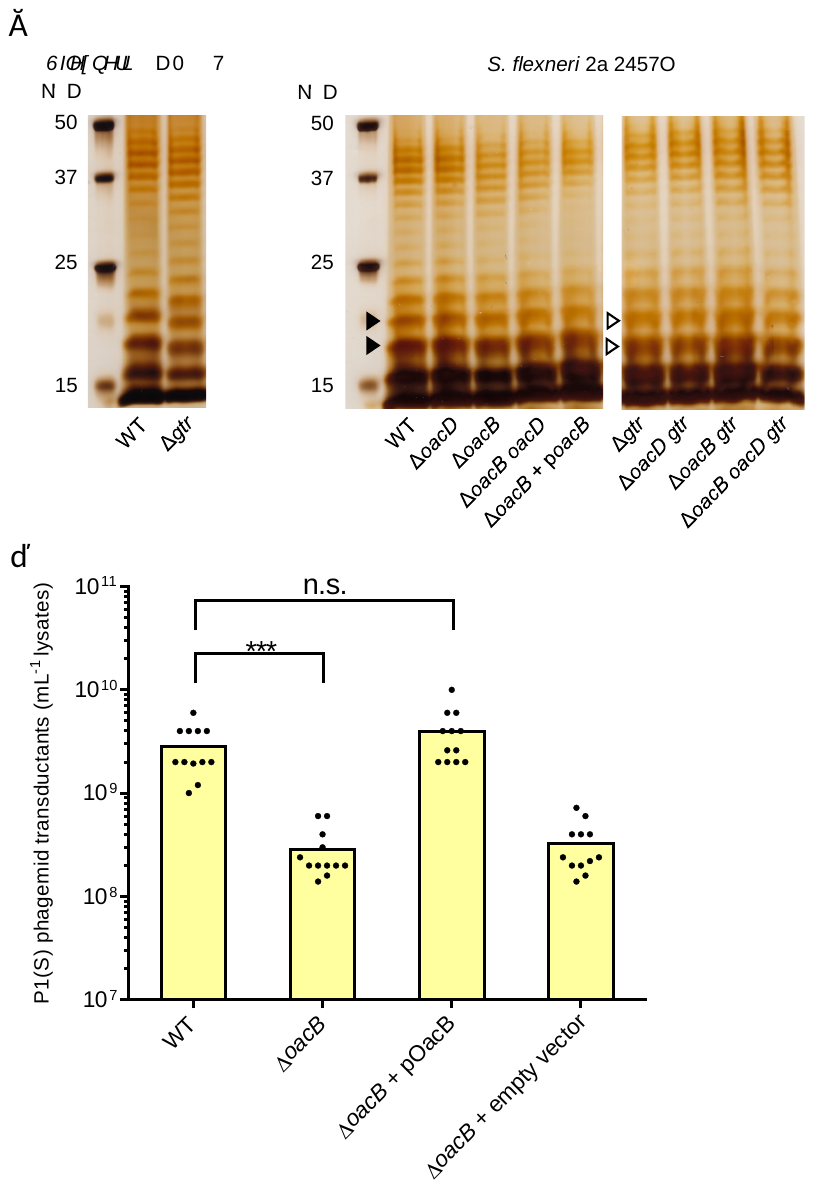

**Supplementary Figure 5: A Δ*oacB* mutation reduced the transduction efficiency of P1(S) on *S. flexneri* 2a 2457O.** (**a**) Analysis of LPS extracted from wildtype (WT) and O-antigen modifications’ mutant strains of *S. flexneri* 5a M90T and *S. flexneri* 2a 2457O with SDS-PAGE and silver staining (refer to **Figure 3a** for details of mutations). Δ*gtr* *S. flexneri* M90T lacks the glucose modification on its O-antigen, hence shift in O-antigen bands (compared to wildtype) was observed (West et al., 2005). For S*. flexneri* 2a 2457O, only the loss of glucose modification of O-antigen caused by Δ*gtr* mutation gave observable shifts in bands, especially band 3 and 4 on silver stain (Teh et al., 2020). (**b**) The number of P1(S) phagemid transductant recovered after lysates treatment of wildtype (WT) and Δ*oacB* strain of *S. flexneri* 2a 2457O, as well as the Δ*oacB* mutant transformed with a pOacB (Δ*oacB* + pOacB) or an empty RK2 vector. p < 0.0005 was shown as ***, for comparison between the number of wildtype (WT) and Δ*oacB* phagemid transductant. n.s. represented no significant difference for comparison in the number of P1(S) phagemid transductant between WT *S. flexneri* 2a 2457O and the pOacB transformant.

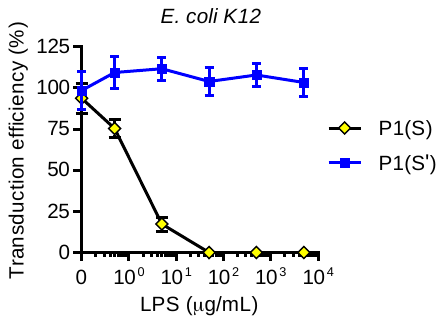

**Supplementary Figure 6: *E. coli* K12 LPS blocked P1(S) transduction but not that of P1(S').** Quantification of P1(S) (yellow diamonds, black line) and P1(S') (blue squares, blue line) transduction efficiencies when pre-incubated with 0.5 μg/mL, 5.0 μg/mL, 50 μg/mL, 500 μg/mL and 5000 μg/mL of commercially available *E. coli* K12 LPS (Invivogen, tlrl-eklps). Each data point represented results from 1 LPS sample, 1 lysates sample, 3 biological repeats (host cells used) and 2 technical repeats. Data were represented as mean ± SEM.

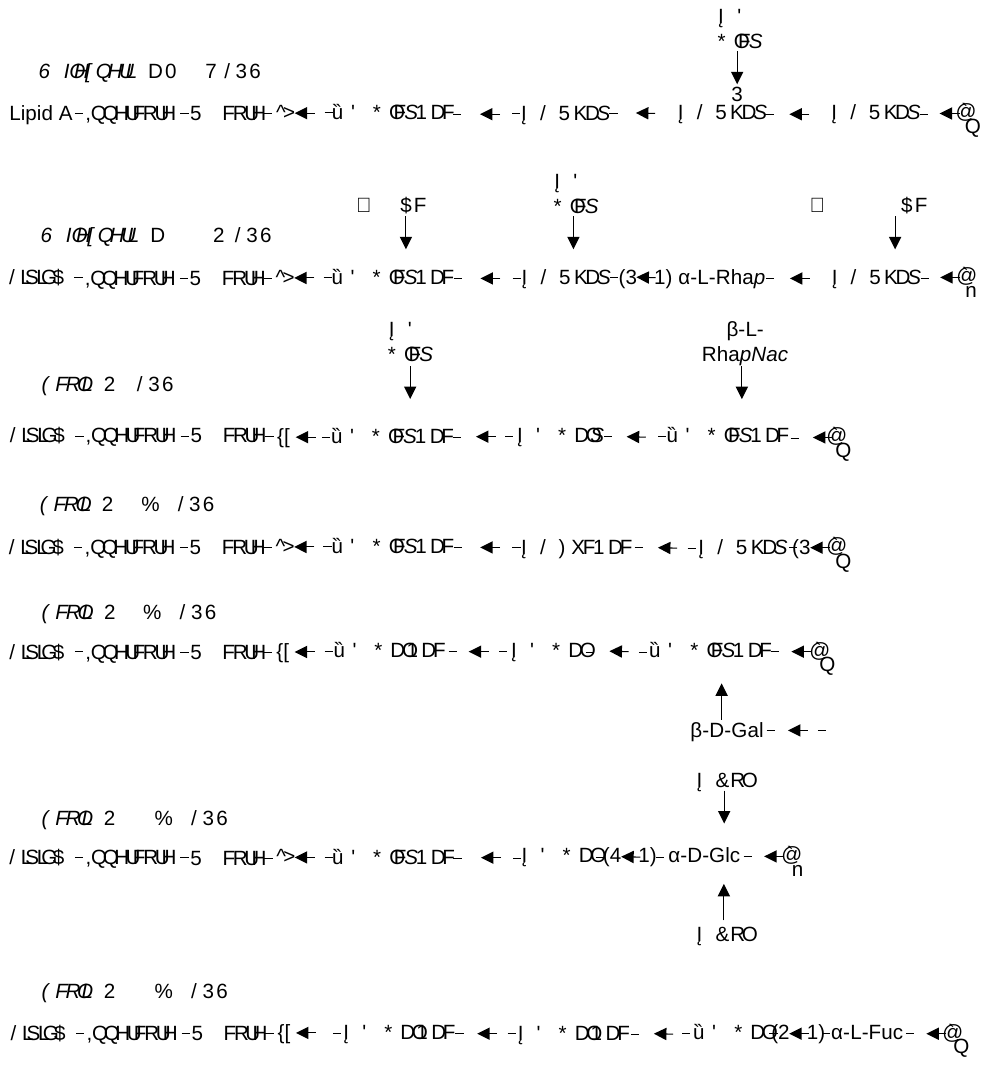

**Supplementary Figure 7: Chemical structures of *S. flexneri* 5a M90T, *S. flexneri* 2a 2457O, *E. coli* O3, *E. coli* O26:B6, *E. coli* O55:B5, *E. coli* O111:B4 and *E. coli* O127:B8 LPS.** The chemical formulas of LPS plotted were based on results of previous studies (Kenne et al., 1978; Stenutz et al., 2006; Ren et al., 2008; Huang et al., 2012; Perepelov et al., 2012). For simplification purpose, the orientation of glycosidic bond linking the O-antigen to the outer oligosaccharide core of LPS, as well as possible modifications to the inner core and lipid A were not shown. The O-antigens were linked to the terminal α-D-Glc of both R2 (Heinrichs et al., 1998) and R3 (Kaniuk et al., 2004) LPS outer oligosaccharide cores via a (O-antigen)-1,4- (terminal glucose of outer core) glycosidic bond. *p*, phosphorylation; *Ac*, acetyl group; GlcNac, *N*-acetylglucosamine; Rha, rhamnose; Glc, glucose, RhaNac; *N*-acetylrhamnosamine; Gal, galactose; GalNac, *N*-acetylgalactosamine; Col, colitose; Fuc, fucose; FucNac, *N*-acetylfucosamine.

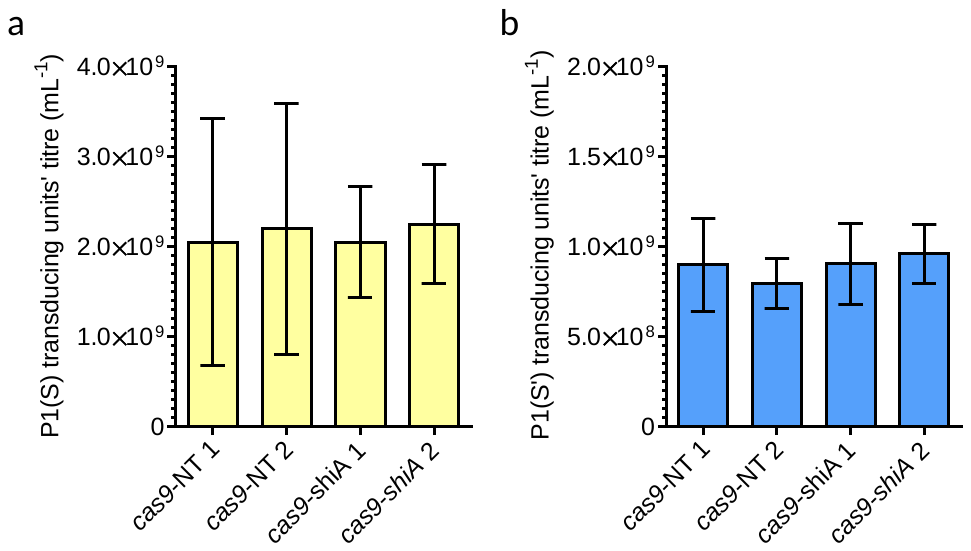

**Supplementary Figures 8: No significant differences between the number of *shiA*-targeting and non-targeting phagemid transductant.** The number of (**a**) P1(S) and (**b**) P1(S') *shiA*-targeting (*cas9*-*shiA*) and non-targeting (*cas9*-NT) phagemid transductant that were recovered after treatment of (**a**) *E. coli* NCM3722 and (**b**) *E. coli* O3, respectively. Data were collected from 2 biological repeats and 2 technical repeats for each lysate. Data were represented as mean ± standard deviation (SD). Welch’s ANOVA test gave p-values of 0.9901 and 0.7094, for comparisons of the mean values in (**a**) and (**b**) respectively, indicating no significant difference in the number of *E. coli* transductant recovered between *shiA*-targeting and non-targeting phagemid lysates treatment. The similar number of *E. coli* phagemid transductant recovered verified that *shiA*-targeting and non-targeting lysates contained a similar titre of phagemid transducing particles.

**Supplementary Table 1:** Bacterial strains used in this study

| **Registry** | **Description** | **Remarks/Source** |
| --- | --- | --- |
| EMG16 *E. coli* K12 C600 Δp*ac* P1_kc_ | *E. coli* P1 lysogen used for phage lysate preparation | Strain generated in this study |
| *E. coli* NCM3722 | Routinely used for quantifying P1(S) transducing units | CGSC#: 12355 |
| *E. coli* serotype O3 strain P1S'ab, designated as *E. coli* O3 in this study | Routinely used for quantifying P1(S') transducing units. Genomic sequencing showed the presence of O3 O-antigen gene cluster and agglutination assay verified the expression of O3 antigen. | Collection of Wang’s Lab, gift from Professor Ian Henderson |
| *ΔwaaL E. coli* O3 | Isogenic mutant strain of *E. coli* O3 with *waaL* deleted | Strain generated in this study |
| *Δrfe E. coli* O3 | Isogenic mutant strain of *E. coli* O3 with *rfe/wecA* deleted | Strain generated in this study |
| *Δwzy E. coli* O3 | Isogenic mutant strain of *E. coli* O3 with *wzy* deleted | Strain generated in this study |
| *E. coli* 24337A | Used for assessment of phagemid transduction efficiency | Collection of Wang’s Lab, gift from Professor Ian Henderson |
| *E. coli* TOP10 | Used for routine molecular cloning, | Thermo Fisher Scientific, Waltham, MA, USA |
| *E. coli* BL21 | Used for assessment of phagemid transduction efficiency | Thermo Fisher Scientific, Waltham, MA, USA |
| *S. flexneri* 2a 2457O | Avirulent mutant which contains a transposon insertion in the major-virulence plasmid which disrupts *virF* and subsequently prevents activation of the *lpa* genes essential for bacterial invasion of the host. | ATCC: 29903 |
| *S. flexneri* *5a* M90T | Virulent strain of *S. flexneri* | Gift from Dr Serge Mostowy |
| *ΔwaaL S. flexneri* 2a 2457O | Isogenic mutant strain of *S. flexneri* 2a 2457O with *waaL* deleted | Strain generated in this study |
| *Δrfe S. flexneri* 2a 2457O | Isogenic mutant strain of *S. flexneri* 2a 2457O with *rfe/wecA* deleted | Strain generated in this study |
| *Δrfc S. flexneri* 2a 2457O | Isogenic mutant strain of *S. flexneri* 2a 2457O with *rfc/wzy* deleted | Strain generated in this study |
| *ΔwaaL S. flexneri* 5a M90T | Isogenic mutant strain of *S. flexneri* 5a M90T with *waaL* deleted | Strain generated in this study |
| *Δrfe S. flexneri* 5a M90T | Isogenic mutant strain of *S. flexneri* 5a M90T with *rfe/wecA* deleted | Strain generated in this study |
| *Δrfc S. flexneri* 5a M90T | Isogenic mutant strain of *S. flexneri* 5a M90T with *rfc/wzy* deleted | Strain generated in this study |
| *Δgtr S. flexneri* 5a M90T | Isogenic mutant strain of *S. flexneri* 5a M90T with *gtrA, gtrB* and *gtrII* deleted | Strain generated in this study |
| *ΔoacD S. flexneri* 2a 2457O | Isogenic mutant strain of *S. flexneri* 2a 2457O with *oacD* deleted | Strain generated in this study |
| *Δgtr S. flexneri* 2a 2457O | Isogenic mutant strain of *S. flexneri* 2a 2457O with *gtrA, gtrB* and *gtrV* deleted | Strain generated in this study |
| *ΔoacB S. flexneri* 2a 2457O | Isogenic mutant strain of *S. flexneri* 2a 2457O with *oacB* deleted | Strain generated in this study |
| *ΔoacD gtr S. flexneri* 2a 2457O | Isogenic mutant strain of *S. flexneri* 2a 2457O with *oacD, gtrA, gtrB* and *gtrV* deleted | Strain generated in this study |
| *ΔoacB gtr S. flexneri* 2a 2457O | Isogenic mutant strain of *S. flexneri* 2a 2457O with *oacB, gtrA, gtrB, gtrV* | Strain generated in this study |
| *ΔoacB oacD S. flexneri* 2a 2457O | Isogenic mutant strain of *S. flexneri* 2a 2457O with *oacB* and *oacD* deleted | Strain generated in this study |
| *ΔoacB oacD gtr S. flexneri* 2a 2457O | Isogenic mutant strain of *S. flexneri* 2a 2457O with *oacB, oacD, gtrA, gtrB* and *gtrV* deleted | Strain generated in this study |

**Supplementary Table 2** Plasmids and phagemids used in this study

| **Registry** | **Description** | **Remarks/Source** |
| --- | --- | --- |
| P1 *cas9*-NT | P1 phagemid with constitutive *cas9* expression, tracrRNA and crRNA guide from *S. pyogenes* derived from pCas9 (Addgene plasmid #42876)*.* Cas9 chromosomal-targeting effect is absent in *S. flexneri*, hence named non-targeting (NT) | Phagemid generated in this study |
| P1 *cas9-shiA* | Cas9 antimicrobial effect on *S. flexneri* is present, due to complementary sequence of guide crRNA spacer sequence to that of chromosomal *shiA* gene sequence | Phagemid generated in this study |
| pkD46 | Lambda red-mediated recombineering, used for genetic modification of *pacA* gene of P1 bacteriophage | CGSC |
| pCP20 | Provide expression of Flippase that excise inserted kanamycin resistant cassette | CGSC |
| pP1SU | P*_sit_* regulating the expression of tail fibre gene *S*, chaperone *U*, with its cognate RBS. Constitutive expression of β-lactamase (*amp*), with pSC101 origin of replication | Plasmid generated in this study |
| pP1S'U' | P*_sit_* regulating the expression of tail fibre gene *S'*, chaperone *U'*, with its cognate RBS. Constitutive expression of β-lactamase (*amp*), with pSC101 origin of replication | Plasmid generated in this study |
| pWaaL-*S. flexneri* | Promoter Bba_J23110 and artificial RBS Bba_B0030 giving constitutive expression of *waaL* gene, cloned from *S. flexneri* 2a 2457O chromosome. Constitutive expression of *npt* conferring kanamycin resistance, with RK2 origin of replication | Plasmid generated in this study |
| pWaaL-*E. coli* O3 | Promoter Bba_J23110 and artificial RBS Bba_B0030 giving constitutive expression of *waaL* gene, cloned from *E. coli* O3 chromosome. Constitutive expression of *npt* conferring kanamycin resistance, with RK2 origin of replication | Plasmid generated in this study |
| pRfe | Promoter Bba_J23110 and artificial RBS Bba_B0030 giving constitutive expression of *rfe* gene, cloned from *S. flexneri* 2a 2457O chromosome. Constitutive expression of *npt* conferring kanamycin resistance, with RK2 origin of replication. | Plasmid generated in this study |
| pOacB | Promoter Bba_J23110 and artificial RBS Bba_B0030 giving constitutive expression of *oacB* gene, cloned from *S. flexneri* 2a 2457O chromosome. Constitutive expression of *npt* conferring kanamycin resistance, with RK2 origin of replication | Plasmid generated in this study |

**Supplementary Table 3:** Primers used for lambda Red recombineering in this study

| **Name of sequence(s)** | **Sequence (5’ to 3’)^a^** | **Remarks** |
| --- | --- | --- |
| LR∆*pac*-f | **gaagacaccaggactacggacagccgcaagccaaataagccagtcaggaagccactaaaa**attccggggatccgtcgacc | Deletion of *pac* site of P1 bacteriophage |
| LR∆*pac*-r | **caacatagcgcgcgcggccttccgcgcttcaacgttatctatcaggtaatcgccaacttc**gtgtaggctggagctgcttc | Deletion of *pac* site of P1 bacteriophage |
| ∆*pac*-f | agccagtcaggaagccacta | Primer to detect ∆*pac* mutation of P1 bacteriophage |
| ∆*pac*-r | gttctccagcataaggagatg | Primer to detect ∆*pac* mutation of P1 bacteriophage |
| LR∆*S-cin*-f | **tataccgcagttatggccataaacacgactacagcataaggatatgcttgctgtcaaacatgaga**attaattccggggatc | Deletion of *S-cin* of P1 bacteriophage |
| LR∆*S-cin*-r | **agtgaattaaatatcattgggaaacggtatgtactttgtgatttccacac**gtgtaggctggagctgcttcgaag | Deletion of *S-cin* of P1 of P1 bacteriophage |
| ∆*S-cin*-f | tggataacgagaacaagccaatc | Primer to detect ∆*S-cin* mutation of P1 bacteriophage |
| ∆*S-cin*-r | tacccttatattccgcgaaatac | Primer to detect ∆*S-cin* mutation of P1 bacteriophage |
| LR∆*waaL-S.flexneri-*f | **aataaccaataagttgacatcggagataagatgacctcaacattatttttctctctcgag**attccggggatccgtcgacc | Deletion of *waaL* of *S. flexneri* |
| LR∆*waaL-S.flexneri-*r | **ggttattcataattggtttgaataaataaaaaggccgcattatgcagccttttttatttt**gtgtaggctggagctgcttc | Deletion of *waaL* of *S. flexneri* |
| ∆*waaL-S. flexneri-*f | tgggatggcgtaactcaaag | Primer to detect ∆*waaL* of *S. flexneri* |
| ∆*waaL-S. flexneri-*r | tgcagatttgccagctcaaa | Primer to detect ∆*waaL* of *S. flexneri* |
| LR∆*waaL-E. coli* O3*-*f | **gattaatacgttatatattatttaagatggtggagaagca**attccggggatccgtcgacc | Deletion of *waaL* of *E. coli* O3 |
| LR∆*waaL-E. coli* O3-r | **tggttaaaaattattggatttgaagagcctgcctgaacttaatgtcaggcaggctttttt**gtgtaggctggagctgcttc | Deletion of *waaL* of *E. coli* O3 |
| ∆*waaL-E. coli* O3*-*f | agtcatttaactgcagcact | Primer to detect ∆*waaL* of *E. coli* O3 |
| ∆*waaL-E. coli* O3-r | atattaagcgcgttttggc | Primer to detect ∆*waaL* of *E. coli* O3 |
| LR∆*rfe-*f | **tctgaataaaggtcttcgtggttatacttctgctaataattttctctgagagcatgcatt**attccggggatccgtcgacc | Deletion of *rfe* of *S. flexneri* and *E. coli* O3 |
| LR∆*rfe-*r | **cacatcctcatttatttggttaaattggggctgccaccacgatttctacgcagtctgcgt**gtgtaggctggagctgcttc | Deletion of *rfe* of *S. flexneri* and *E. coli* O3 |
| ∆*rfe-*f | cgaaacaacccacgaatatcc | Primer to detect ∆*rfe* of *S. flexneri* and *E. coli* O3 |
| ∆*rfe-*r | tatcaaatggttgatttttgcacag | Primer to detect ∆*rfe* of *S. flexneri* and *E. coli* O3 |
| LR∆*rfc-S. flexneri-*f | **aaatttatattttattaatgttgattttaaacatattttaaaaaacatcaccttacgaat**attccggggatccgtcgacc | Deletion of *rfc* of *S. flexneri* |
| LR∆*rfc-S. flexneri*-r | **tttattttgctccagaagtgaggttattactaatttggatattttctatagaaaataccc**gtgtaggctggagctgcttc | Deletion of *rfc* of *S. flexneri* |
| ∆*rfc-S. flexneri-*f | ctggtcaataacttccctatttttaac | Primer to detect ∆*rfc* of *S. flexneri* |
| ∆*rfc-S. flexneri-*r | ggaagttaaggcggaaaaag | Primer to detect ∆*rfc* of *S. flexneri* |
| LR∆*wzy-E. coli* O3*-*f | **ttatttttctttcattgcttcctgttttaattaaagagggagctagagggaatggcaaat**attccggggatccgtcgacc | Deletion of *wzy* of *E. coli* O3 |
| LR∆*wzy-E. coli* O3-r | **cattgaattatagtacataaaaatcaaaggatttaataacgcgatccctatagaacttaa**gtgtaggctggagctgcttc | Deletion of *wzy* of *E. coli* O3 |
| ∆*wzy-E. coli* O3*-*f | aagccaggatgataattgaaaaggaattat | Primer to detect ∆*wzy* of *E. coli* O3 |
| ∆*wzy-E. coli* O3-r | taccaaatgacccagtaccaccag | Primer to detect ∆*wzy* of *E. coli* O3 |
| LR∆*gtr* M90T*-*f | **acacacttatacactgggtggtttttggtgtatgtatctatgccgcacataccagtcaggc**attccggggatccgtcgacc | Deletion of *gtr* of *S. flexneri* 5a M90T (West et al., 2005) |
| LR∆*gtr* M90T*-*r | **cgggggggatttattgatgcaaaacttagatcaataacattaagtaactaccattcaaca**gtgtaggctggagctgcttc | Deletion of *gtr* of *S. flexneri* 5a M90T (West et al., 2005) |
| ∆*gtr* M90T*-*f | ggatgaagaaagaggccg | Primer to detect ∆*gtr* of *S. flexneri* 5a M90T |
| ∆*gtr* M90T*-*r | atgttaaagttatttgtaaagtacac | Primer to detect ∆*gtr* of *S. flexneri* 5a M90T |
| LR∆*oacD* 2457O*-*f | **tacacaaatacaggtatatatgattgcgcagataggatttggaatagtcacgctgaagcc**attccggggatccgtcgacc | Deletion of *oacD* of *S. flexneri* 2a 2457O (Teh et al., 2020) |
| LR∆*oacD* 2457O*-*r | **gtcaccttgggttgggggctggcaaattgtatccttttgtttttagttatcacgtcccac**gtgtaggctggagctgcttc | Deletion of *oacD* of *S. flexneri* 2a 2457O (Teh et al., 2020) |
| ∆*oacD* 2457O*-*f | aatggcccaccgactgtc | Primer to detect ∆*oacD* of *S. flexneri* 2a 2457O |
| ∆*oacD* 2457O*-*r | actggagagcctactttaaatg | Primer to detect ∆*oacD* of *S. flexneri* 2a 2457O |
| LR∆*gtr* 2457O*-*f | **ggaggcgctctttaataataaataaatctatcaaagctaaatattaaatggaagccaccc**attccggggatccgtcgacc | Deletion of *gtr* of *S. flexneri* 2a 2457O (Teh et al., 2020) |
| LR∆*gtr* 2457O*-*r | **acacacttatacactgggtggtttttggtgtatgtatctatgccgcacataccagtcaggc**gtgtaggctggagctgcttc | Deletion of *gtr* of *S. flexneri* 2a 2457O (Teh et al., 2020) |
| ∆*gtr* 2457O*-*f | gtgattgcatccattgatg | Primer to detect ∆*gtr* of *S. flexneri* 2a 2457O |
| ∆*gtr* 2457O*-*r | atgttaaagttatttgtaaagtacac | Primer to detect ∆*gtr* of *S. flexneri* 2a 2457O |
| LR∆*oacB* 2457O*-*f | **aaatcttgttgagcacggcaggaataatcaaatagatggaatgcgggggttcttagcaat**attccggggatccgtcgacc | Deletion of *oacB* of *S. flexneri* 2a 2457O (Teh et al., 2020) |
| LR∆*oacB* 2457O*-*r | **aatataaatacattcgccgccccatttggacatatatgtagggcggcttatgtttattgt**gtgtaggctggagctgcttc | Deletion of *oacB* of *S. flexneri* 2a 2457O (Teh et al., 2020) |
| ∆*oacB* 2457O*-*f | ccgtgatattgatgtcgt | Primer to detect ∆*oacB* of *S. flexneri* 2a 2457O |
| ∆*oacB* 2457O*-*r | gtttgcatttagcattaccag | Primer to detect ∆*oacB* of *S. flexneri* 2a 2457O |

^a^DNA sequences in bold represent the 60 bps complementary sequence to the target gene.

**Supplementary Tables 4:** DNA sequences

DNA sequence of tail fibre genes and its chaperones

Promoter LP*_S_* regulating *S* and *U* or *S'* and *U'* genes expressions with the native RBS

aagttacttatcttacaatgaggcttcacaacattgattagggaaaatc

*S* tail fibre gene*, with Sc* (constant region) in black and *Sv* (variable region) in blue

atgaatgacgttacagttgttacatcggttacttacccatcatccgagtcgttggctctggtggccgatgtgcaataccacgaaccatatctgtcagccgcgctaaaccgaaaattcagggggattgttgacccgggattttatgccggtttcttacctaagcctggcggtgggatgaacctgttaattacctcagtggatggtgataaaaccgcaggcgcggcgtcggtggatattggtgaattttaccaggtaactattcagcaacgtaaggatatttctcttgcacttagtgcaggcaagaaatatgcaattgtgctgaagggaagatacctccttggagaagatacctaccaggtgaataccgcgtcacatattcatgcggctgaatttgttgccagaacctataccgattcatatcagttaggagatggggagctgcttgtttgtacggtgaatatccctgctagtgtatctgccattacccaggagatgattgatacatccgagcgtatcaaccgctcgatcggcattgatatttcagactctgtaaccagtaccagaagtgatgttgctgcaagttcgctggcagttaaaaaagcctacgatctggcgaaaagcaagtatacggcacaggatgcaagcacaacgcaaaagggattagttcagctcagtagcgcaactaacagcgacagcgaaacaatggcggctacccctaaagctgttaagtctataaaagatctggctgataccaaagcgccaatagaaagcccgagtctgacaggaacgccaaccgcgccgacggcagcgcaaggtacaaacagcacgcagatcgcaaatacagcctttgttaaggcagctataactgcacttatcaacggtgcgcctggcacactggatacgctgaaagaaatagcggctgcgatcaataacgacccgaattacagcacaactatcaacaatgccttggctctcaaagcgcctttggcaagccctgcattaacgggtgtccctactgcgcctacggctgcacagggcacaaacaatacgcagatcgctacgactgcttacgtacgggctgctatctctgcattggtcggctcatcacctgaagctcttgataccctgtatgagcttgcagcagcactgggcaatgacccgaactttgcgacaacaatgacaaatgcgctggcagggaaacagccacttgatgcaactttaaccgcgcttgctggtcttgcgacaggcgcaaataaattgccgtactttaccggtacagacactgtttctcagactgacttaacgtcagttggtcgcgatattctggccaaaacaagcattcttgctgttatccaataccttggtttaagagaactcggtaccagcggtgaaaagatccccctgttgagcacggctaacacatggagtgcacgccagactttcaacggcgggatcaccggggcgctgacagggaacgccgataccgcaacgaaattgaaaacagccagaaacattaatggcgtcaggttcgatggttctggtgacattaatatcaatactctggtatcgcgcggtcgcgtaacggccctggaggcgaatgcacagggaacatccgggattcagctgtatgaggcatacaacaatggctacccttccccctatggcaatgtgcttcaccttaaaggtgccaccgctgctggcgaaggtgagttattcattggctggagtggcacgagcggtgcccatgcgcccgtacatatccgttcgcggcgggatactgattctgccaactggtctgaatgggcgcaggtctatacgtcaaaagattcaattcccggcgtcaatgccaaaggggatcaggatacctctggtaatgcggctacagcgaccaagttgcagacagcatgtactatcaacggcgtctcgtttgacggttctaaaaatattgagctaacggcggaagatttaaatctacaggaaacggtaaacaaggctgataacgcggttcaaaagacaggcgataccttgtccggtggacttacttttgaaaacgactcaatccttgcctggattcggaatactgactgggcgaagattggttttaaaaatgatgccgacagcgatactgattcatacatgtggtttgaaacaggcgacaacggcaatgaatatttcaaatggagaagcaaacaaagtaccacaacaaaagacctgatgaatcttaaatgggatgctttgtatgttcttgtcaatgccattgtaaatggcgaagtcatatcaaaatcagcaaacggcctacgtattgcttatggtaattacggattctttattcgtaatgatggttcaaatacatacttcatgttgacaaactccggtgacaacatggggacttataacggattaaggccattatggattaataacgctactggcgctgtttcgatggggcgtggccttaatgtttcaggggagacactttcagaccgttttgctattaacagcagtaatggtatgtggattcagatgcgcgataacaacgctatctttgggaaaaatatagttaacactgatagcactcaggcgttacttcgccagaatcacgccgaccgaaagttcatgataggtggactggggaacaagcaatttggcatctacatgattaataactcaaggacagccaatggcaccgatggtcaggcgtacatggacaataacggtaactggctttgcggtgcgcaaattattcccggaaattatggcaattttgactcacgctatgtgagagatgtccgacttggcacacgtgttgttcaattgatggcgcgtggagggcgttatgaaagagccggacacgcacttaccggattaaggattattggtgaagttgatggcgatgatgacgctatcttcaggccgatacaaaaatacatcaatggcatatggtataacgtcgcacaggtgtaa

*U* after *Sv*, with bases ATT between both coding sequences

ATTatgcagcacttaaaaaatatcaggtcaggaaacccaaagacaaaagagcaataccaattaacaaagaattttgacgtaatctggttgtggtctgaagacggaaaaaactggtatgaggaagtgaaaaactttcaaccagacaccataaagattgtttacgatgaaaataatattattgtcgccatcaccaaagatgcctccacgcttaaccctgaaggttttagcgtcgttgaggttcccgacataacagccaaccgccgcgctgatgattcaggaaagtggatgtttaaggatggagctgtagttaaacggatttatacggcagacgaacagcaacaacaagccgaatcacaaaaggccgcattgctttccgaagctgaatcagtcatccagccgctggaacgcgctgtcaggctgaatatggcaacagacgaggaacgcacacgactggaagcatgggaacgctacagtgttctggtcagccgtgtggatacggcaaatcctgaatggccacaaaagcctgaataa

*S'* tail fibre gene*, with Sc* (constant region) in black and *Sv'* (variable region) in blue

atgaatgacgttacagttgttacatcggttacttacccatcatccgagtcgttggctctggtggccgatgtgcaataccacgaaccatatctgtcagccgcgctaaaccgaaaattcagggggattgttgacccgggattttatgccggtttcttacctaagcctggcggtgggatgaacctgttaattacctcagtggatggtgataaaaccgcaggcgcggcgtcggtggatattggtgaattttaccaggtaactattcagcaacgtaaggatatttctcttgcacttagtgcaggcaagaaatatgcaattgtgctgaagggaagatacctccttggagaagatacctaccaggtgaataccgcgtcacatattcatgcggctgaatttgttgccagaacctataccgattcatatcagttaggagatggggagctgcttgtttgtacggtgaatatccctgctagtgtatctgccattacccaggagatgattgatacatccgagcgtatcaaccgctcgatcggcattgatatttcagactctgtaaccagtaccagaagtgatgttgctgcaagttcgctggcagttaaaaaagcctacgatctggcgaaaagcaagtatacggcacaggatgcaagcacaacgcaaaagggattagttcagctcagtagcgcaactaacagcgacagcgaaacaatggcggctacccctaaagctgttaagtctataaaagatctggctgataccaaagcgccaatagaaagcccgagtctgacaggaacgccaaccgcgccgacggcagcgcaaggtacaaacagcacgcagatcgcaaatacagcctttgttaaggcagctataactgcacttatcaacggtgcgcctggcacactggatacgctgaaagaaatagcggctgcgatcaataacgacccgaattacagcacaactatcaacaatgccttggctctcaaagcgcctttggcaagccctgcattaacgggtgtccctactgcgcctacggctgcacagggcacaaacaatacgcagatcgctacgactgcttacgtacgggctgctatctctgcattggtcggctcatcacctgaagctcttgataccctgtatgagcttgcagcagcactgggcaatgacccgaactttgcgacaacaatgacaaatgcgctggcagggaaacagccacttgatgcaactttaaccgcgcttgctggtcttgcgacaggcgcaaataaattgccgtactttaccggtacagacactgtttctcagactgacttaacgtcagttggtcgcgatattctggccaaaacaagcattcttgctgttatccaataccttggtttaagagaactcggtaccagcggtgaaaagatccccctgttgagcacggctaacacatggagtgcacgccagactttcaacggcgggatcaccggggcgctgacagggaacgccgataccgcaacgaaattgaaaacagccagaaacattaatggcgtcaggttcgatggttctggtgacattaatatcaatactctggtatcgcgcggtcgcgtaacggccctggaggcgaatgcacagggaacatccgggattcagctgtatgaggcatacaacaatggctacccttccccctatggcaatgtgcttcaccttaaaggtgccaccgctgctggcgaaggtgagttattcattggctggagtggcacgagcggtgcccatgcgcccgtacatatccgttcgcggcgggatactgattctgccaactggtctgaatgggcgcaggtctatacgtcaaaagattcaattcccggcgtcaatgccaaaggggatcaggatacctctggtaatgcggctacagcgaccaagttgcagacagcatgtactatcaacggtgtctcgtttgatggttctaaaaatattgagctaacggctgaaaatttaaatcttgagcgaacagtagaattagccgctgggtcattgcagaaaaatcagaacggcgcggatattcctggaaaagataccttcacaaaaaatattggtgcatgtcgcgcttttcacagttctattagtacaggtgcagggaactggacaacggcacaattgattgaatggctggattctcaaggggcattcaatcacccatactggatgtgcaaatgttcatggtcgtacggcaataataaaattataaccgatactggctgtggaactattcatcttgcaggttgcgttattgaggttatgggtaataaaggtgccatgaccatccgtgtaacaacaccaagcacttccagcggtggcggaatcactaacgctcaattcacttatattaatcatggtgatgcttacgctcctggctggcgacgagactacaacacgaaaaacctgcaacctgcatttgctttagggcagacaggaaacagggttgcaaatgataaagctgttggctggaactggaatagcggtgtttatgatgcagacctaaaaggcgcatcaacattaattcttcatttcaatatgaacgcgggtagctgcccggctgtacaattacgcgtgaattataagaacggcggtatttattatcgttcagcgcgtgatggttatggatttgaggctgactggtcagagttttacaccacaacccgcaaaccctctgcgggggatgttggtgcatatacgcaggcagaatgtaactcaaggtttattacaggtattcgcctgggcggtctgtcatctgtccagacatggaatggccccggctggtctgacaggtcaggttatgtcgttacgggttcagttaacgggaaccgtgatgaattaattgatacaacacaggcaaggccaattcagtattgcattaatgggacgtggtataacgcggggagtatttaa

*U'* after *Sv'*, with bases TT between both coding sequences

TTatgatgcacttaagaaatattacagctggcaaccctaaaacaaaagagcaataccagctaacgaaacaatttaacatcaaatggctttatacagaggatggaaaaaactggtatgaggaacaaaagaatttccagtatgatacgttgaaaatggcctatgaccacaacggcgttattatttgtattgaaaaggatgtttcagcaattaatccagaaggcgcaagcgtcgttgaattacctgatattacagcaaatcgccgggctgatatttctggtaaatggatgttcaaagatggcgtagtggtaaagcgaacttataccgaggaagagcagaggcaacaagcggaaaatgaaaagcaaagtctgctacagctcgtcagggataaaacccagctatgggactcacagctacggctgggtatcatttccgccgagaataagcagaaattaaccgagtggatgctctttgcgcagaaagtcgaatccacagacacctccagcctaccagtaacatttcccgaacaacctgaatga

T7 terminator sequence after *U* or *U'* coding sequence

ctgctaacaaagcccgaaaggaagctgagttggctgctgccaccgctgagcaataactagcataaccccttggggcctctaaacgggtcttgaggggttttttgctgaaaggaggaactatatccggat

4A3 plasmid backbone with pSC101 *ori*, conferring ampicillin resistance (*bla* coding sequence highlighted in blue)

taataatactagtagcggccgctgcagtccggcaaaaaagggcaaggtgtcaccaccctgccctttttctttaaaaccgaaaagattacttcgcgttatgcaggcttcctcgctcactgactcgctgcgctcggtcgttcggctgcggcgagcggtatcagctcactcaaaggcggtaatctcgaggttacattgtcgatctgttcatggtgaacagctttaaatgcaccaaaaactcgtaaaagctctgatgtatctatcttttttacaccgttttcatctgtgcatatggacagttttccctttgatatctaacggtgaacagttgttctacttttgtttgttagtcttgatgcttcactgatagatacaagagccataagaacctcagatccttccgtatttagccagtatgttctctagtgtggttcgttgtttttgcgtgagccatgagaacgaaccattgagatcatgcttactttgcatgtcactcaaaaattttgcctcaaaactggtgagctgaatttttgcagttaaagcatcgtgtagtgtttttcttagtccgttacgtaggtaggaatctgatgtaatggttgttggtattttgtcaccattcatttttatctggttgttctcaagttcggttacgagatccatttgtctatctagttcaacttggaaaatcaacgtatcagtcgggcggcctcgcttatcaaccaccaatttcatattgctgtaagtgtttaaatctttacttattggtttcaaaacccattggttaagccttttaaactcatggtagttattttcaagcattaacatgaacttaaattcatcaaggctaatctctatatttgccttgtgagttttcttttgtgttagttcttttaataaccactcataaatcctcatagagtatttgttttcaaaagacttaacatgttccagattatattttatgaatttttttaactggaaaagataaggcaatatctcttcactaaaaactaattctaatttttcgcttgagaacttggcatagtttgtccactggaaaatctcaaagcctttaaccaaaggattcctgatttccacagttctcgtcatcagctctctggttgctttagctaatacaccataagcattttccctactgatgttcatcatctgagcgtattggttataagtgaacgataccgtccgttctttccttgtagggttttcaatcgtggggttgagtagtgccacacagcataaaattagcttggtttcatgctccgttaagtcatagcgactaatcgctagttcatttgctttgaaaacaactaattcagacatacatctcaattggtctaggtgattttaatcactataccaattgagatgggctagtcaatgataattacatgtccttttcctttgagttgtgggtatctgtaaattctgctagacctttgctggaaaacttgtaaattctgctagaccctctgtaaattccgctagacctttgtgtgttttttttgtttatattcaagtggttataatttatagaataaagaaagaataaaaaaagataaaaagaatagatcccagccctgtgtataactcactactttagtcagttccgcagtattacaaaaggatgtcgcaaacgctgtttgctcctctacaaaacagaccttaaaaccctaaaggcttaagtagcaccctcgcaagctcgggcaaatcgctgaatattccttttgtctccgaccatcaggcacctgagtcgctgtctttttcgtgacattcagttcgctgcgctcacggctctggcagtgaatgggggtaaatggcactacaggcgccttttatggattcatgcaaggaaactacccataatacaagaaaagcccgtcacgggcttctcagggcgttttatggcgggtctgctatgtggtgctatctgactttttgctgttcagcagttcctgccctctgattttccagtctgaccacttcggattatcccgtgacaggtcattcagactggctaatgcacccagtaaggcagcggtatcatcaacaggcttacccgtcttactgtccctagtgcttggattctcaccaataaaaaacgcccggcggcaaccgagcgttctgaacaaatccagatggagttctgaggtcattactggatctatcaacaggagtccaagcgagctcgtaaacttggtctgacagttaccaatgcttaatcagtgaggcacctatctcagcgatctgtctatttcgttcatccatagttgcctgactccccgtcgtgtagataactacgatacgggagggcttaccatctggccccagtgctgcaatgataccgcgagacccacgctcaccggctccagatttatcagcaataaaccagccagccggaagggccgagcgcagaagtggtcctgcaactttatccgcctccatccagtctattaattgttgccgggaagctagagtaagtagttcgccagttaatagtttgcgcaacgttgttgccattgctacaggcatcgtggtgtcacgctcgtcgtttggtatggcttcattcagctccggttcccaacgatcaaggcgagttacatgatcccccatgttgtgcaaaaaagcggttagctccttcggtcctccgatcgttgtcagaagtaagttggccgcagtgttatcactcatggttatggcagcactgcataattctcttactgtcatgccatccgtaagatgcttttctgtgactggtgagtactcaaccaagtcattctgagaatagtgtatgcggcgaccgagttgctcttgcccggcgtcaatacgggataataccgcgccacatagcagaactttaaaagtgctcatcattggaaaacgttcttcggggcgaaaactctcaaggatcttaccgctgttgagatccagttcgatgtaacccactcgtgcacccaactgatcttcagcatcttttactttcaccagcgtttctgggtgagcaaaaacaggaaggcaaaatgccgcaaaaaagggaataagggcgacacggaaatgttgaatactcatactcttcctttttcaatattattgaagcatttatcagggttattgtctcatgagcggatacatatttgaatgtatttagaaaaataaacaaataggggttccgcgcacatttcccatggtgccacctgacgtctaagaaaccattattatcatgacattaacctataaaaataggcgtatcacgaggcagaatttcagataaaaaaaatccttagctttcgctaaggatgatttctg

DNA sequence of J72114 phagemid (Kittleson et al., 2012)

P_BAD_ promoter regulating *coi* expression (Kittleson et al., 2012), and that of *pacA* and *pacB*

aagaaaccaattgtccatattgcatcagacattgccgtcactgcgtcttttactggctcttctcgctaaccaaaccggtaaccccgcttattaaaagcattctgtaacaaagcgggaccaaagccatgacaaaaacgcgtaacaaaagtgtctataatcacggcagaaaagtccacattgattatttgcacggcgtcacactttgctatgccatagcatttttatccataagattagcggatcttacctgacgctttttatcgcaactctctactgtttctccat

*pacA* gene sequence with its native RBS (Kittleson et al., 2012)

cgaaaggaagcataagtgacctgggacgatcacaagaagaattttgctcgcctggcgcgagatggtggttacaccatcgcacagtatgccgccgagtttaatcttaaccctaataccgcacgtcgttatctccgtgccttcaaagaagacaccaggactacggacagccgcaagccaaataagccagtcaggaagccactaaaaagcatgatcattgatcactctaatgatcaacatgcaggtgatcacattgcggctgaaatagcggaaaaacaaagagttaatgccgttgtcagtgccgcagtcgagaatgcgaagcgccaaaataagcgcataaatgatcgttcagatgatcatgacgtgatcacccgcgcccaccggaccttacgtgatcgcctggaacgcgacaccctggatgatgatggtgaacgctttgagttcgaagttggcgattacctgatagataacgttgaagcgcggaaggccgcgcgcgctatgttgcgtcggtccggggccgatgttctggaaaccactcttctggaaaagtctctttctcatctccttatgctggagaacgccagggatacgtgtattcgcctggtgcaggaaatgcgcgatcagcaaaaagacgatgatgaaggtactccgcctgaataccgtatcgcgagcatgctaaacagctgttccgcgcagataagcagcctgatcaacaccatttacagcatccggaataactatcgaaaagaaagccgggaggcggaaaagcacgctttatctatggggcaagctggcattgttaagctggcatacgaacgaaagcgtgaaaataactggtcagtgctggaagcggctgagttcatcgaggcgcatggaggaaaagtgccgcccctgatgctggagcaaatcaaagccgatctgcgtgctcctaagaccaataccgatgatgaggaaaaccaaacagcatctggcgctccatcacttgaggatctggataaaatcgcgcgagaacgggccgccagccgccgcgctgatgccgcattgtggattgagcatcgtagagaagaaattgccgatatcgtcgatacaggtggttatggtgatgtcgatgcggaaggcatatcaaacgaagcatggcttgaacaggatctggacgaagacgaggaggaagacgaagaagttacccgcaaactgtacggggatgatgattaa

*pacB* gene sequence with an A base overlapping with *pacA* coding sequence (in capital letter), native terminator sequence in blue

Atggccagaagttgcgtaacggacccacgttggcgcgagcttgtggcgctatatcgttatgactggattgcggccgctgatgtgttgtttgggaagacaccaacctggcagcaggatgagatcattgagtccacgcagcaggacggcagttggacaagtgtgacctccggccatggtactggtaaatcggatatgacgagtatcattgcaatactcttcatcatgtttttccccggcgctcgcgtcattctggtcgctaacaaaagacagcaagtccttgatggtattttcaaatacataaagagcaattgggctactgctgttagcagattcccgtggttgtcgaagtatttcattcttacagaaacgtctttttttgaggtgactggcaagggtgtttggacaatattgataaagtcctgtcgtcccggaaatgaggaggcgttggctggtgaacacgccgatcatctcttgtatatcatcgacgaagcgtcgggtgtgagtgataaagcattcagtgtgataacaggtgcgctgaccggtaaggataaccgtattctgcttctttcccagcctacgcgaccttcaggctatttctacgattcacaccacagactagctattcgcccgggaaatcctgatggattgtttactgcgataatactgaatagtgaagaatctccgcttgtagatgcaaaatttatacgagcaaaacttgcggagtatggcggtcgtgataaccccatgtacatgatcaaagtacgtggtgaatttcccaaatctcaagatggctttcttcttggtcgtgatgaggttgagcgggcgacgcggcgaaaggtcaagattgccaaaggatggggctgggttgcatgtgttgacgttgctggtggcacaggacgagataagtccgttattaatatcatgatggtgtccggccagcgaaataaacgccgtgtaatcaactatcgtatgctggaatacacagacgttacagaaacgcagttagccgccaagattttcgcagaatgtaacccagaacggttcccgaacataaccatagctattgatggcgatggcttggggaaatcgacggctgatctaatgtacgaacgctatggcattaccgtccagcgtatccgctggggtaaaaagatgcacagccgtgaagataaaagcctttatttcgatatgcgcgctttcgcgaatattcaggcggcagaagctgtaaaatcagggcgtatgaggcttgataagggggctgcgactatagaggaagcatcaaagataccggtagggataaattccgcaggtcaatggaaggtgatgtcaaaggaagatatgaagaaaaaactcaacctgcactcaccggaccattgggatacatattgtttcgctatgttggcgaactatgttccccaagatgaagtgcttagcgtcgaagacgaagcgcaggttgatgaagctctggcatggcttaatgaataactcattaaccatgccggatggaaa

Non-coding *repL* sequence (without start codon) (Kittleson et al., 2012)

ttaccctctgaatcctgccggtataccccattgttcgttatctttatttttggctaaaaccgcattaagagcttcgtttaccgtcatgcaatgcggtaggttatcgaagtttgatatcccgccaatatcaggcgaacgcttgttTttcaggtaagcatatttccgcgcagccgcctctactttctgcttgaactcatgtttttgagtgcgttttttggataaccgcagattgtcagcctttgcttttgccttagcgatccatgaagtcaattttttgaggctggttgttccggcaccgccggaaactgatctttttgtttttttaacttgtgacttcttattctttattgccacgtcatcctgacagggggagggggtatcattttgacatgggggtgtggataaaaaattaaataaagccaatgtcttagcgagaacagctttaaccttggttgccgctgaCgaaatctttaatttgctttctatcagcgcatttttggcttgttgtgcgaaggccaaaaaggatggtgtaaaccggtacaggttagcgcgacgttcacggtgatcgccgataacaatctctacagacagaatacctttgtttacagcttcacggaatgcacgaacgacggttgattggctataaccagtttctgccgcgatcaggcggtgaggcttgtgaatgaagtattcactggttgttgccgcgagatttgcacattgcgacaggatatgcccggcgctacgggatagaccggagtgtgttacaaagcaggccaattcatagccagaaaaagtaaaatcgcttta

Non-coding *cin* sequence (without start codon) (Kittleson et al., 2012)

ccgagttctcttaaaccaaggtttaggattgaaatgatgacgccggaaacttcttataaagcgtggaaacagccacatcatagatgattgcaacctgcttacgggggatgcccttctccagcaatcgccgcatttgctgccatgtttcttcttggtatttaggccgacgcccacctatacgaccttctgcgcgagctgcatcaagtccagcgcgtgtacgttcaacgataagctcacgttccatttctgccagcgcccccattacgtgaaagaaaaagcgccccattggtgtactggtgtcgatggagtcagtgagactccggaagttaatgcctctgtcacgcagctcttccaccagcacaactaagtgacgcatgctgcgcccaagacggtctaacttccatacgaccagggtatcacctctggaaagTtacggagtaccttttttaacccagggcgctcagcctttttgccgctcgccttgtcctcaaaaattagctcacatcctgcgctttcaagagcgtttcgttgtaaagcagtgttttgttcatttgttgatacgcgtacatagcctattag

*araC* with its native promoter and RBS (Kittleson et al., 2012)

ctctgaatggcgggagtatgaaaagtatggctgaagcgcaaaatgatcccctgctgccgggatactcgtttaatgcccatctggtggcgggtttaacgccgattgaggccaacggttatctcgatttttttatcgaccgaccgctgggaatgaaaggttatattctcaatctcaccattcgcggtcagggggtggtgaaaaatcagggacgagaatttgtttgccgaccgggtgatattttgctgttcccgccaggagagattcatcactacggtcgtcatccggaggctcgcgaatggtatcaccagtgggtttactttcgtccgcgcgcctactggcatgaatggcttaactggccgtcaatatttgccaatacggggttctttcgcccggatgaagcgcaccagccgcatttcagcgacctgtttgggcaaatcattaacgccgggcaaggggaagggcgctattcggagctgctggcgataaatctgcttgagcaattgttactgcggcgcatggaagcgattaacgagtcgctccatccaccgatggataatcgggtacgcgaggcttgtcagtacatcagcgatcacctggcagacagcaattttgatatcgccagcgtcgcacagcatgtttgcttgtcgccgtcgcgtctgtcacatcttttccgccagcagttagggattagcgtcttaagctggcgcgaggaccaacgtatcagccaggcgaagctgcttttgagcaccacccggatgcctatcgccaccgtcggtcgcaatgttggttttgacgatcaactctatttctcgcgggtatttaaaaaatgcaccggggccagcccgagcgagttccgtgccggttgtgaagaaaaagtgaatgatgtagccgtcaagttgtcataa

*coi* with its native RBS (Kittleson et al., 2012)

tacagtgaggcataattatggctttcattccaccaaccatcgacgacgttagacattgctctaacgctttatctgtagaccccgccgaaaccgacgctgcccgcgccattgctgaacactactcaaagatatccaatcaggagtaccgcatcacccaagacgacctggatgatctcactgacacaatcgaatatctcatggccactaaccagccagactcacaataa

rrnB T1 terminator after *coi* coding sequence (Kittleson et al., 2012)

caaataaaacgaaaggctcagtcgaaagactgggcctttcgttttatctgttgtttgtcggtgaacgctctc

J72114 vector backbone, with p15A ori, *cat* gene in blue conferring chloramphenicol resistance (Kittleson et al., 2012)

agtcagagtagaatagaagtatcattctcaccaataaaaaacgcccggcggcaaccgagcgttctgaacaaatccagatggagttctgaggtcattactggatctatcaacaggagtccaagcgagctcgatatcaaattacgccccgccctgccactcatcgcagtactgttgtaattcattaagcattctgccgacatggaagccatcacaaacggcatgatgaacctgaatcgccagcggcatcagcaccttgtcgccttgcgtataatatttgcccatggtgaaaacgggggcgaagaagttgtccatattggccacgtttaaatcaaaactggtgaaactcacccagggattggctgagacgaaaaacatattctcaataaaccctttagggaaataggccaggttttcaccgtaacacgccacatcttgcgaatatatgtgtagaaactgccggaaatcgtcgtggtattcactccagagcgatgaaaacgtttcagtttgctcatggaaaacggtgtaacaagggtgaacactatcccatatcaccagctcaccgtctttcattgccatacgaaattccggatgagcattcatcaggcgggcaagaatgtgaataaaggccggataaaacttgtgcttatttttctttacggtctttaaaaaggccgtaatatccagctgaacggtctggttataggtacattgagcaactgactgaaatgcctcaaaatgttctttacgatgccattgggatatatcaacggtggtatatccagtgatttttttctccattttagcttccttagctcctgaaaatctcgataactcaaaaaatacgcccggtagtgatcttatttcattatggtgaaagttggaacctcttacgtgcccgatcaaagatccgcaccgccggacatcagcgctagcggagtgtatactggcttactatgttggcactgatgagggtgtcagtgaagtgcttcatgtggcaggagaaaaaaggctgcaccggtgcgtcagcagaatatgtgatacaggatatattccgcttcctcgctcactgactcgctacgctcggtcgttcgactgcggcgagcggaaatggcttacgaacggggcggagatttcctggaagatgccaggaagatacttaacagggaagtgagagggccgcggcaaagccgtttttccataggctccgcccccctgacaagcatcacgaaatctgacgctcaaatcagtggtggcgaaacccgacaggactataaagataccaggcgtttccccctggcggctccctcgtgcgctctcctgttcctgcctttcggtttaccggtgtcattccgctgttatggccgcgtttgtctcattccacgcctgacactcagttccgggtaggcagttcgctccaagctggactgtatgcacgaaccccccgttcagtccgaccgctgcgccttatccggtaactatcgtcttgagtccaacccggaaagacatgcaaaagcaccactggcagcagccactggtaattgatttagaggagttagtcttgaagtcatgcgccggttaaggctaaactgaaaggacaagttttggtgactgcgctcctccaagccagttacctcggttcaaagagttggtagctcagagaaccttcgaaaaaccgccctgcaaggcggttttttcgttttcagagcaagagattacgcgcagaccaaaacgatctcaagaagatcatcttattaatcagataaaatatttctaaggcctcccctgattctgtggataaccgggatctgtaaggatcaaccactttgtacaagaaagctgggtcgaattgagatccgaacggtttattacgtacatcaggtaaaactgaccgataagccgctttcttttgggtatagtgtcgtggacagtcattcatctttctgcccctccaaaagtaaaaacccgccgaagcgggtttttacgtaaaacaggtgaaactgaccgataagccgctttcttttgggtatagtgtcgtggacagtcattcatctttctgcccctccaaaagcaaaaacccgccgaagcgggtttttacgtaaaccaggtgaaactgaccgataagccgctttcttttgggtatagcgtcgtggacagtcattcatctttccgcccctccaaaagcaaaaacccgccgaagcgggtttttacgtaaatcaggtgaaactgaccgataagccgggttctgtcgtggacagtcattcatctaggccagcaatcgctcagatcc

DNA sequence of CRISPR Cas9 construct derived from pCas9 plasmid (Addgene plasmid #42876)

Promoter sequence of *trans*-activating *crRNA*

agtattaagtattgttttatggctgataaatttctttgaatttctccttgattatttgttataaaagttataaaataatcttgtt

*Trans*-activating crRNA

ggaaccattcaaaacagcatagcaagttaaaataaggctagtccgttatcaacttgaaaaagtggcaccgagtcggtgc

*cas9* native promoter and RBS

atagaatgataacaaaataaactactttttaaaagaattttgtgttataatctatttattattaagtattgggtaatattttttgaagagatattttgaaaaagaaaaattaaagcatattaaactaatttcggaggtcattaaaactattattgaaatcatcaaactcattatggatttaatttaaactttttattttaggaggcaaaa

crRNA guide leader sequence in upper case, direct repeats (in blue) and protospacer sequence (in black), BsaI sites are underlined

TATTTCTTAATAACTAAAAATATGGTATAATACTCTTAATAAATGCAGTAATACAGGGGCTTTTCAAGACTGAAGTCTAGCTGAGACAAATAGTGCGATTACGAAATTTTTTAGACAAAAATAGTCTACGAGgttttagagctatgctgttttgaatggtcccaaaactgagaccagtctcggaagctcaaaggtctcgttttagagctatgctgttttgaatggtcccaaaac

*cas9*, DNA in upper case represent the region between *cas9* ORF and crRNA leader sequence

atggataagaaatactcaataggcttagatatcggcacaaatagcgtcggatgggcggtgatcactgatgaatataaggttccgtctaaaaagttcaaggttctgggaaatacagaccgccacagtatcaaaaaaaatcttataggggctcttttatttgacagtggagagacagcggaagcgactcgtctcaaacggacagctcgtagaaggtatacacgtcggaagaatcgtatttgttatctacaggagattttttcaaatgagatggcgaaagtagatgatagtttctttcatcgacttgaagagtcttttttggtggaagaagacaagaagcatgaacgtcatcctatttttggaaatatagtagatgaagttgcttatcatgagaaatatccaactatctatcatctgcgaaaaaaattggtagattctactgataaagcggatttgcgcttaatctatttggccttagcgcatatgattaagtttcgtggtcattttttgattgagggagatttaaatcctgataatagtgatgtggacaaactatttatccagttggtacaaacctacaatcaattatttgaagaaaaccctattaacgcaagtggagtagatgctaaagcgattctttctgcacgattgagtaaatcaagacgattagaaaatctcattgctcagctccccggtgagaagaaaaatggcttatttgggaatctcattgctttgtcattgggtttgacccctaattttaaatcaaattttgatttggcagaagatgctaaattacagctttcaaaagatacttacgatgatgatttagataatttattggcgcaaattggagatcaatatgctgatttgtttttggcagctaagaatttatcagatgctattttactttcagatatcctaagagtaaatactgaaataactaaggctcccctatcagcttcaatgattaaacgctacgatgaacatcatcaagacttgactcttttaaaagctttagttcgacaacaacttccagaaaagtataaagaaatcttttttgatcaatcaaaaaacggatatgcaggttatattgatgggggagctagccaagaagaattttataaatttatcaaaccaattttagaaaaaatggatggtactgaggaattattggtgaaactaaatcgtgaagatttgctgcgcaagcaacggacctttgacaacggctctattccccatcaaattcacttgggtgagctgcatgctattttgagaagacaagaagacttttatccatttttaaaagacaatcgtgagaagattgaaaaaatcttgacttttcgaattccttattatgttggtccattggcgcgtggcaatagtcgttttgcatggatgactcggaagtctgaagaaacaattaccccatggaattttgaagaagttgtcgataaaggtgcttcagctcaatcatttattgaacgcatgacaaactttgataaaaatcttccaaatgaaaaagtactaccaaaacatagtttgctttatgagtattttacggtttataacgaattgacaaaggtcaaatatgttactgaaggaatgcgaaaaccagcatttctttcaggtgaacagaagaaagccattgttgatttactcttcaaaacaaatcgaaaagtaaccgttaagcaattaaaagaagattatttcaaaaaaatagaatgttttgatagtgttgaaatttcaggagttgaagatagatttaatgcttcattaggtacctaccatgatttgctaaaaattattaaagataaagattttttggataatgaagaaaatgaagatatcttagaggatattgttttaacattgaccttatttgaagatagggagatgattgaggaaagacttaaaacatatgctcacctctttgatgataaggtgatgaaacagcttaaacgtcgccgttatactggttggggacgtttgtctcgaaaattgattaatggtattagggataagcaatctggcaaaacaatattagattttttgaaatcagatggttttgccaatcgcaattttatgcagctgatccatgatgatagtttgacatttaaagaagacattcaaaaagcacaagtgtctggacaaggcgatagtttacatgaacatattgcaaatttagctggtagccctgctattaaaaaaggtattttacagactgtaaaagttgttgatgaattggtcaaagtaatggggcggcataagccagaaaatatcgttattgaaatggcacgtgaaaatcagacaactcaaaagggccagaaaaattcgcgagagcgtatgaaacgaatcgaagaaggtatcaaagaattaggaagtcagattcttaaagagcatcctgttgaaaatactcaattgcaaaatgaaaagctctatctctattatctccaaaatggaagagacatgtatgtggaccaagaattagatattaatcgtttaagtgattatgatgtcgatcacattgttccacaaagtttccttaaagacgattcaatagacaataaggtcttaacgcgttctgataaaaatcgtggtaaatcggataacgttccaagtgaagaagtagtcaaaaagatgaaaaactattggagacaacttctaaacgccaagttaatcactcaacgtaagtttgataatttaacgaaagctgaacgtggaggtttgagtgaacttgataaagctggttttatcaaacgccaattggttgaaactcgccaaatcactaagcatgtggcacaaattttggatagtcgcatgaatactaaatacgatgaaaatgataaacttattcgagaggttaaagtgattaccttaaaatctaaattagtttctgacttccgaaaagatttccaattctataaagtacgtgagattaacaattaccatcatgcccatgatgcgtatctaaatgccgtcgttggaactgctttgattaagaaatatccaaaacttgaatcggagtttgtctatggtgattataaagtttatgatgttcgtaaaatgattgctaagtctgagcaagaaataggcaaagcaaccgcaaaatatttcttttactctaatatcatgaacttcttcaaaacagaaattacacttgcaaatggagagattcgcaaacgccctctaatcgaaactaatggggaaactggagaaattgtctgggataaagggcgagattttgccacagtgcgcaaagtattgtccatgccccaagtcaatattgtcaagaaaacagaagtacagacaggcggattctccaaggagtcaattttaccaaaaagaaattcggacaagcttattgctcgtaaaaaagactgggatccaaaaaaatatggtggttttgatagtccaacggtagcttattcagtcctagtggttgctaaggtggaaaaagggaaatcgaagaagttaaaatccgttaaagagttactagggatcacaattatggaaagaagttcctttgaaaaaaatccgattgactttttagaagctaaaggatataaggaagttaaaaaagacttaatcattaaactacctaaatatagtctttttgagttagaaaacggtcgtaaacggatgctggctagtgccggagaattacaaaaaggaaatgagctggctctgccaagcaaatatgtgaattttttatatttagctagtcattatgaaaagttgaagggtagtccagaagataacgaacaaaaacaattgtttgtggagcagcataagcattatttagatgagattattgagcaaatcagtgaattttctaagcgtgttattttagcagatgccaatttagataaagttcttagtgcatataacaaacatagagacaaaccaatacgtgaacaagcagaaaatattattcatttatttacgttgacgaatcttggagctcccgctgcttttaaatattttgatacaacaattgatcgtaaacgatatacgtctacaaaagaagttttagatgccactcttatccatcaatccatcactggtctttatgaaacacgcattgatttgagtcagctaggaggtgactgaAGTATATTTTAGATGAAGAT

Coding sequences of *waaL*, *rfe*, *oacB*

J23110 regulating the expressions of *waaL, rfe, oacB*

tttacggctagctcagtcctaggtacaatgctagc

RBS 30 for translation of WaaL, Rfe and OacB

attaaagaggagaaa

*waaL* of *S. flexneri*

atgacctcaacattatttttctctctcgagaaaaaaaactggatagcgtactggaacagagctctcgtattcttattcattaccacctattttttgggtgggattacaaggtataaacatcttattgttattcttatgacaataacgacaatcgtctatctctgcaaacggccaaaacactatctctcactatttaaaacatttctttttggtagtgttgccatattaactattgctgcattgctgtcacttcttcaatcccctgatgcaggtgctagcatgaaggaagttttcaaagctattattgagaatactttactatgcacaatagcaataccggtcatattgagagacgagaaaagagaagatgtcgaaaaaatcgttttcttctcatttattagtgcgttgggcttacgctgtttttctgaattgattacctattataaggactatcaacaagggataatgccattcgcagattatagacaccgtagcatttctgactcgatggtctttttattccctgcattgttaaatctctggcttatcaaatcagcaaaataccgcatttcttttgtggttctaagcgttatttttatttttctgatattaggaactttatccagaggggcctggctttccgtgttagtcattggattaatatggattctgatgtttaaacaatggaagttactattagtaggagtaatggtttccatcattgcattgtcggttattttcacacataaggagatgaccgcaaagctaacgtataaacttcaacaaactaatagttcttatcgctatgcaaatggtactcaaggcagcgcactcgatctaatattagaaaatcctgttattggttatggttacggtaacgttgcatataaagatgtctataataaacgtgtcattgattatccagaatggacctttagacaatcaatagggccacataattttgcgctattcatctggtttggcactggtttattagggctggtaagtcttatgatgctatactgtgcaatattgaaagagtgtataaaaaatggcgtcaagaataaatatcgctcaccatataatgcatattatataatcttactatcttttataggttattttgttatccgtggaaacgtagaacaaattgaaccaaatttattaggcgtttacgccggcttattattagcgatgaaaaacaagtaa

*waaL* of *E. coli* O3

atgttaacagcctcgctggcgctacgtaataaagaaaagtggaaaccgtactggaataaagcgttggtattcctctttattgcaacctttttccttgatggtataacccgttataaacacatcatatctatactaatgattattacggtaatctatcaagtttctcgtgctccgggaacatttaaagtactctataaaaataatcttttttatagtgtcctggcactatcactcattcttttatatgctacttttatatcacctgatctgaaaatcagttttaaagaattcagtaatacggtacttaaaggatttttggcctatagtttacttatacctgcattattaaaagatgaagataatgaaagcattggtaaaattgtactatactcattagttaccggacttggcttacgttgtcttgttgaacttattctttatattcaggattataataaaggaataatgccattttccacctatgaacaccgtagtatttccgattcaatggtgtttctgtttccagcattattaaacctttggcttatcaagaaaacctcttataaaatagcattcgtaatttttagtgcagttttcttatttctgctattaggaacgctgtctcgtggtgcatggctggcagtattcatagtgactctgctatggttaatcttaaatcgtcagtggaaattattgatgctgacttccatcgttatttctgttgccgcagtgggtgtatttacctataaaggtgatcatgctggtaaagacaggcttatttataaactgcaacagaccgacagttcttctcgctataccaatggtactcaaggtacggcgtggacattaattatggaaaatcctcttaagggatatggctacggcgatgatatttaccatgcaatatataataagcgtgttgtagattttccgtcatggaagttcaggcaatccattgggccacacaacgttgttttgtcaatttggtttgcagcgggtttggctggactactcgctttactttatctgtatggctctattatcaaagagacagctaatgcaacgtttaaaacagttgtcgtcactccttataatggtcaattactactatttttaacgcttgtaagcttttatattattcgaggtaacttcgaggaagttgatctcaaaccaattggcttgattgttggtttactgttagcaatgcggaataaataa

*rfe* of *E. coli* MG1655

gtgaatttactgacagtgagtactgatctcatcagtatttttttattcacgacactgtttctgttttttgcccgtaaggtggcaaaaaaagtcggtttagtggataaaccaaacttccgcaaacgtcaccagggattgatacctctcgttggggggatttcggtttacgcagggatttgcttcacgttcggaattgtcgattactatattccgcatgcatctctctatctcgcttgtgccggtgtgcttgttttcattggcgcgctggatgatcgttttgatatcagcgtaaaaatccgtgccaccatacaggccgctgttggcattgttatgatggtgttcggcaaactttatctcagtagcctgggttatatctttggctcctgggagatggtgctcggaccgtttggttacttcctgacgctatttgccgtctgggcggccattaatgcgttcaacatggttgatggcattgatggcttgctgggcgggttgtcctgcgtctcgtttgcagcaatcggtatgattttgtggttcgacgggcaaaccagcctcgcaatctggtgctttgcgatgatcgccgccatcctgccatacatcatgcttaaccttggtatcctgggtcgccgctacaaagtctttatgggtgatgcgggcagtacgctgattggttttaccgttatctggatcctgctcgaaacgacccagggcaaaacccatcccatcagcccggttaccgctttgtggataatcgccattccgctaatggatatggtggcgattatgtaccgtcgcctgcgtaaaggcatgagcccattctctcctgaccgtcagcatattcaccatttgatcatgcgtgccgggtttacttcccgtcaggcgtttgtgctgattacccttgccgcagcactgctcgcttccattggcgtgctggcagaatattctcattttgtcccggagtgggtcatgctggtgctctttttgctagcattcttcctctatggatattgcattaagcgtgcctggaaagttgctcgctttattaagcgcgtaaaacgcagactgcgtagaaatcgtggtggcagccccaatttaaccaaataa

*oacB* of *S. flexneri* 2a 2457O

atgcatatgattgaaattaacagtttattattaataacatccgtgatattgatgtcgttattagctgtggggttatttgataaaatttccccaataaatcttgttgagcacggcaggaataatcaaatagatggaatgcgggggttcttagcaattttcgtgcttattcatcacgcagcaatttggaatggatacttgtcatctggagtatgggaagcaccttcatcaaatctgttagcaaatttaggccaagttggtgtgtcattcttttttatgattactggttatctgttcttttcaaagattatctctggagatcaggactggacaaggctttatgtatcaagattactacgattaaccccaatgttcatagttagcctatgtctcatttttatcattgtaggttttaagtctggatggagaatgcaggtatccacagaagagctttttgtgtcaataatgaagtggttgccattcactgcgctaggtatgccgaacattaatgacgtaaaagattcatttactatcaatgccgctgtaacatggacacttgtatatgaatggttcttttatttttctcttccggtaatttccgcgctcataaaaagaaaagtcagtatttacatggttatgattagcgcaatatcgctatttgttttcattttatttttcagtaaaatacacatagcatcatttctttttgggctcctggcatttttgctaaataaatcaaagatagtcaatggaattgccaaagcaaaagtaacaccaattataataactgcgataatggtttttgaaatgacgtatttcaaaacaacttacgcaccgctaccactgattctttgtggaataacatttatcattatcgcatcaggttgtgatttgtatggaatattaagattaaatataaccagaaagctaggagaaactacctacagcgtttaccttctgcatgggatattcctttattgcctaatgacgtggattattcctaataattacactgaaaatacatttatcatactggtaagcacaactgcatttcttattacgttcacatcatgcctaacttttaaattaatagagacgccattcatcaaattaacaaaacaaaccacaactttagtaaaagaattaatacccacattaacaaacaacaatcaataa

RK2 plasmid backbone providing kanamycin resistance (*npt*, coding sequence in blue)

cttcttggtcgtcatagttcctcgcgtgtcgatggtcatcgacttcgccaaacctgccgcctcctgttcgagacgacgcgaacgctccacggcggccgatggcgcgggcagggcagggggagccagttgcacgctgtcgcgctcgatcttggccgtagcttgctggactatcgagccgacggactggaaggtttcgcggggcgcacgcatgacggtgcggcttgcgatggtttcggcatcctcggcggaaaaccccgcgtcgatcagttcttgcctgtatgccttccggtcaaacgtccgattcattcaccctccttgcgggattgccccggaattaattccccggatcgatccgtcgatcttgatcccctgcgccatcagatccttggcggcaagaaagccatccagtttactttgcagggcttcccaaccttaccagagggcgccccagctggcaattccggttcgcttgctgtccataaaaccgcccagtctagctatcgccatgtaagcccactgcaagctacctgctttctctttgcgcttgcgttttcccttgtccagatagcccagtagctgacattcatccggggtcagcaccgtttctgcggactggctttctacgtggctgccatttttggggtgaggccgttcgcggccgaggggcgcagcccctggggggatgggaggcccgcgttagcgggccgggagggttcgagaagggggggcaccccccttcggcgtgcgcggtcacgcgcacagggcgcagccctggttaaaaacaaggtttataaatattggtttaaaagcaggttaaaagacaggttagcggtggccgaaaaacgggcggaaacccttgcaaatgctggattttctgcctgtggacagcccctcaaatgtcaataggtgcgcccctcatctgtcagcactctgcccctcaagtgtcaaggatcgcgcccctcatctgtcagtagtcgcgcccctcaagtgtcaataccgcagggcacttatccccaggcttgtccacatcatctgtgggaaactcgcgtaaaatcaggcgttttcgccgatttgcgaggctggccagctccacgtcgccggccgaaatcgagcctgcccctcatctgtcaacgccgcgccgggtgagtcggcccctcaagtgtcaacgtccgcccctcatctgtcagtgagggccaagttttccgcgaggtatccacaacgccggcggccctacatggctctgctgtagtgagtgggttgcgctccggcagcggtcctgatcccccgcagaaaaaaaggatctcaagaagatcctttgatcttttctacggcgcgcccagctgtctagggcggcggctcggtaccaaattccagaaaagaggcctcccgaaaggggggccttttttcgttttggtccctgcagcggccgctactagtatataaacgcagaaaggcccacccgaaggtgagccagtgtgactctagtagagagcgttcaccgacaaacaacagataaaacgaaaggcccagtctttcgactgagcctttcgttttatttgatgcctggctctagtattactctagaagcggccgcgaattcgtcgtgactgggaaaaccctggcgactagtcttggactcctgttgatagatccagtaatgacctcagaactccatctggatttgttcagaacgctcggttgccgccgggcgttttttattggtgagaatccaggggtccccaataattacgatttaaatttgtgtctcaaaatctctgatgttacattgcacaagataaaaatatatcatcatgaacaataaaactgtctgcttacataaacagtaatacaaggggtgttatgagccatattcagcgtgaaacgagctgtagccgtccgcgtctgaacagcaacatggatgcggatctgtatggctataaatgggcgcgtgataacgtgggtcagagcggcgcgaccatttatcgtctgtatggcaaaccggatgcgccggaactgtttctgaaacatggcaaaggcagcgtggcgaacgatgtgaccgatgaaatggtgcgtctgaactggctgaccgaatttatgccgctgccgaccattaaacattttattcgcaccccggatgatgcgtggctgctgaccaccgcgattccgggcaaaaccgcgtttcaggtgctggaagaatatccggatagcggcgaaaacattgtggatgcgctggccgtgtttctgcgtcgtctgcatagcattccggtgtgcaactgcccgtttaacagcgatcgtgtgtttcgtctggcccaggcgcagagccgtatgaacaacggcctggtggatgcgagcgattttgatgatgaacgtaacggctggccggtggaacaggtgtggaaagaaatgcataaactgctgccgtttagcccggatagcgtggtgacccacggcgattttagcctggataacctgattttcgatgaaggcaaactgattggctgcattgatgtgggccgtgtgggcattgcggatcgttatcaggatctggccattctgtggaactgcctgggcgaatttagcccgagcctgcaaaaacgtctgtttcagaaatatggcattgataatccggatatgaacaaactgcaatttcatctgatgctggatgaatttttctaataattaattggaccgcggtccgcgcgttgtccttttccgctgcataaccctgcttcggggtcattatagcgattttttcggtatatccatcctttttcgcacgatatacaggattttgccaaagggttcgtgtagactttccttggtgtatccaacggcgtcagccgggcaggataggtgaagtaggcccacccgcgagcgggtgttccttcttcactgtcccttattcgcacctggcggtgctcaacgggaatcctgctctgcgaggctggccgtaggccggccgcgatgcaggtggctgctgaacccccagccggaactgaccccacaaggccctagcgtttgcaatgcaccaggtcatcattgacccaggcgtgttccaccaggccgctgcctcgcaactcttcgcaggcttcgccgacctgctcgcgccacttcttcacgcgggtggaatccgatccgcacatgaggcggaaggtttccagcttgagcgggtacggctcccggtgcgagctgaaatagtcgaacatccgtcgggccgtcggcgacagcttgcggtacttctcccatatgaatttcgtgtagtggtcgccagcaaacagcacgacgatttcctcgtcgatcaggacctggcaacgggacgttttcttgccacggtccaggacgcggaagcggtgcagcagcgacaccgattccaggtgcccaacgcggtcggacgtgaagcccatcgccgtcgcctgtaggcgcgacaggcattcctcggccttcgtgtaataccggccattgatcgaccagcccaggtcctggcaaagctcgtagaacgtgaaggtgatcggctcgccgataggggtgcgcttcgcgtactccaacacctgctgccacaccagttcgtcatcgtcggcccgcagctcgacgccggtgtaggtgatcttcacgtccttgttgacgtggaaaatgaccttgttttgcagcgcctcgcgcgggattttcttgttgcgcgtggtgaacagggcagagcgggccgtgtcgtttggcatcgctcgcatcgtgtccggccacggcgcaatatcgaacaaggaaagctgcatttccttgatctgctgcttcgtgtgtttcagcaacgcggcctgcttggcttcgctgacctgttttgccaggtcctcgccggcggtttttcg

Lambda Red recombineering template, flippase recognition motifs (FRT) in red, kanamycin resistance gene (*npt*) in blue

attccggggatccgtcgacctgcagttcgaagttcctattctctagaaagtataggaacttcagagcgcttttgaagctcacgctgccgcaagcactcagggcgcaagggctgctaaaggaagcggaacacgtagaaagccagtccgcagaaacggtgctgaccccggatgaatgtcagctactgggctatctggacaagggaaaacgcaagcgcaaagagaaagcaggtagcttgcagtgggcttacatggcgatagctagactgggcggttttatggacagcaagcgaaccggaattgccagctggggcgccctctggtaaggttgggaagccctgcaaagtaaactggatggctttcttgccgccaaggatctgatggcgcaggggatcaagatctgatcaagagacaggatgaggatcgtttcgcatgattgaacaagatggattgcacgcaggttctccggccgcttgggtggagaggctattcggctatgactgggcacaacagacaatcggctgctctgatgccgccgtgttccggctgtcagcgcaggggcgcccggttctttttgtcaagaccgacctgtccggtgccctgaatgaactgcaggacgaggcagcgcggctatcgtggctggccacgacgggcgttccttgcgcagctgtgctcgacgttgtcactgaagcgggaagggactggctgctattgggcgaagtgccggggcaggatctcctgtcatctcaccttgctcctgccgagaaagtatccatcatggctgatgcaatgcggcggctgcatacgcttgatccggctacctgcccattcgaccaccaagcgaaacatcgcatcgagcgagcacgtactcggatggaagccggtcttgtcgatcaggatgatctggacgaagagcatcaggggctcgcgccagccgaactgttcgccaggctcaaggcgcgcatgcccgacggcgaggatctcgtcgtgacccatggcgatgcctgcttgccgaatatcatggtggaaaatggccgcttttctggattcatcgactgtggccggctgggtgtggcggaccgctatcaggacatagcgttggctacccgtgatattgctgaagagcttggcggcgaatgggctgaccgcttcctcgtgctttacggtatcgccgctcccgattcgcagcgcatcgccttctatcgccttcttgacgagttcttctaataaggggatcttgaagttcctattccgaagttcctattctctagaaagtataggaacttcgaagcagctccagcctacac

**Supplementary Materials and Methods**

**DNase I treatment of phagemid lysates**

Phagemid lysates were treated with Amplification Grade DNase I (Sigma, AMPD1) following manufacturer’s protocol to eliminate DNA that was not packaged into phage particles. Briefly, 1 μL of DNase was used to treat 1 μL of phagemid lysate in a final volume of 10 μL reaction mix, for 20 mins at room temperature. 1 μL of stop solution was added to inhibit DNase activity, followed by heat-inactivation of the enzyme at 70 °C for 15 mins. Ultrapure deionized water was added to the reaction mix, giving a final 100 X diluted phagemid lysate.

**Preparation of standards for qPCR**

Linear DNA fragment of *cas9* sequence was used as standards, which was generated with PCR, using Q5® High-Fidelity DNA Polymerase (NEB, M0491) and following manufacturer’s protocol. The primer pairs used for amplifying the *cas9* DNA fragment was 5’-CTCAATAGGCTTAGATATCGGCACAAATAGC-3’ and 5’- CTTCAGTCACCTCCTAGCTGACTCAAATC-3’, giving an amplicon of 4096 bps in size. PCR product was run on 1 % agarose gel and DNA was eluted Monarch® DNA Gel Extraction Kit (NEB, T1020), following manufacturer’s protocol. The quality and quantity of DNA sample was assessed on a DeNovix DS-11 Series Spectrophotometer (Denovix), followed by sequence verification via Sanger sequencing. Serial dilutions of the *cas9* DNA were made from 30 ng/μL to 3 X 10^-1^ ng/μL in ultrapure deionized water. The concentrations of diluted samples were confirmed using Qubit™ 1 X dsDNA High Sensitivity (HS) on a Qubit 4 Fluorometer (Thermo). Further serial dilutions of the *cas9* DNA fragment were made from 3 X 10^-1^ ng/μL to 3 X 10^-6^ ng/μL.

**SYBR Green qPCR**

SYBR™ Green PCR Master Mix (Thermo, 4309155) was used to quantify the copy number of packaged phagemid DNA on a LightCycler® 96 Instrument (Roche). The primer set used for qPCR was 5’-CTAGTGCCGGAGAATTACAA-3’ and 5’- CTTCTGGACTACCCTTCAAC-3’, which amplified a 108 bps region of *cas9* DNA sequence on the phagemid. A final primer concentration of 300 nM was used for qPCR. A concentration range of 3 X 10^-1^ ng/μL to 3 X 10^-6^ ng/μL of the *cas9* DNA fragment was used to generate the standard curve. The 10 reaction mix was prepared with 5 μL of SYBR Green PCR Master Mix, 1 μL of DNase I treated phagemid lysate, 300 nM primer pair and 3.4 μL of ultrapure deionized water. 3 technical repeats were performed for the standards at each concentration, as well as each phagemid lysate. PCR cycling conditions were pre-incubation step at 95 °C for 10 mins; 45 cycles of 95 °C for 20 s, 60 °C for 20 s, 72 °C for 20 s; melting at 95 °C for 10 s, 65 °C for 60 s, 97 °C for 1 s.

**Quantification of plaque-forming units (PFU)**

For results presented in **Supplementary Figure 1c**, plaque assay was carried out to estimate the population of wildtype P1 phage in lysates, using a protocol established by Kropinski et al., (2009). Briefly, stationary phase culture of *E. coli* NCM3722 was diluted in fresh PLM (LB broth supplemented with 5 mM Ca^2+^ and 10 mM Mg^2+^) by 1/100, cultured at 37 °C with shaking until it reached an OD_600_ of 0.5. From our preliminary results, phage lysates would have to be diluted by 10^6^ to 10^5^ to give a reasonable amount of 10 to 100 PFU for enumeration. 1 mL of cells suspension was added to 100 μL of diluted phage lysate, vortexed, and incubated at 37 °C with shaking for 10 mins. Cells and phage mixtures were added to 3 mL of melted LC top agar, then poured immediately onto LC bottom agar plates. Agar plates were dried at room temperature for 10 mins, then incubated at 37 °C for at least 16 h, before the enumeration of PFU.
